# Structural basis of CD28 and CTLA-4 interactions with CD80, CD86, and the CD80-PD-L1 heterodimer on artificial and cellular membranes

**DOI:** 10.64898/2026.03.30.715283

**Authors:** Gu Min Han, Chan Seok Lim, Jie-Oh Lee

## Abstract

The opposing actions of the co-stimulatory receptor CD28 and the co-inhibitory receptor CTLA-4, mediated by their interactions with B7-family ligands, govern the balance between T-cell activation and immune tolerance. We determined the cryo-EM structures of CD28 and CTLA-4 bound to CD80, CD86, and the CD80-PD-L1 heterodimer under two-dimensional membrane confinement. We show that CD28-CD80, CD28-CD86, and CTLA-4-CD86 form discrete complexes with closed-leg, cross-leg, and open-leg configurations, respectively, whereas CTLA-4 assembles into extended linear clusters and two-dimensional lattices with CD80. PD-L1 binding to CD80 disrupts CTLA-4-CD80 clustering by preventing CD80 homodimerization, while preserving CD28-CD80 binding through a pronounced architectural rearrangement. We further confirm the formation of linear and two-dimensional CTLA-4-CD80 clusters in cell-derived membrane fragments using cryo-EM. Together, these findings demonstrate that immune checkpoint signaling is governed not only by receptor-ligand affinity but also by membrane-imposed geometry and the competitive reorganization of higher-order assemblies at the immunological synapse.

## INTRODUCTION

T cell activation and immune tolerance are tightly regulated by the coordinated engagement of co-stimulatory and co-inhibitory receptors at the immunological synapse ^1–3^. Among these, the CD28 family receptors CD28 and CTLA-4 play central and opposing roles by interacting with the B7 family ligands CD80 and CD86 expressed on antigen-presenting cells ^4,5^. CD28 delivers a potent co-stimulatory signal that lowers the activation threshold of naïve T cells, whereas CTLA-4 functions as a dominant negative regulator that suppresses immune activation and maintains peripheral tolerance. Dysregulation of this pathway underlies a wide range of pathological conditions, including autoimmunity, chronic infection, and cancer, and therapeutic targeting of CD28, CTLA-4, and their ligands has transformed clinical immuno-oncology ^6,7^. Adding further complexity, PD-L1, best known as the ligand for PD-1, has emerged as an important modulator of the CD28/CTLA-4 pathway through its ability to bind CD80 ^8–11^. This interaction suppresses CTLA-4-mediated transendocytosis of CD80 without impairing CD28-mediated co-stimulatory signaling, consistent with a modulatory role for PD-L1 in organizing immune checkpoint interactions at the immunological synapse. However, the structural basis for how PD-L1 engagement reshapes CD80 architecture, alters receptor stoichiometry, and disrupts higher-order CTLA-4 assemblies has remained unclear, largely due to a lack of structural information obtained in a membrane-bound context.

CD28 and CTLA-4 are type I transmembrane receptors of the CD28 family. Each consists of a single extracellular V-type immunoglobulin (IgV) domain, a transmembrane helix, and a cytoplasmic tail, forming disulfide-linked homodimers that engage B7 ligands through the conserved MYPPPY motif ^12–14^. In contrast, the B7 family ligands CD80, CD86, and PD-L1 are type I transmembrane proteins with two extracellular Ig-like domains: an N-terminal IgV domain responsible for receptor binding, and a membrane-proximal IgC-like domain that provides structural spacing from the membrane. These are followed by a single transmembrane helix and a short cytoplasmic tail ^13–16^. Structurally, CD80 has a strong intrinsic tendency to homodimerize, whereas CD86 is mostly monomeric on the cell membrane ^15,17,18^. Together, this domain architecture supports a geometrically constrained, bivalent receptor-ligand system that underlies the balance between CD28-mediated co-stimulation and CTLA-4-mediated inhibition at the immunological synapse ^19,20^.

Despite extensive biochemical, cellular, and structural studies, the molecular mechanisms by which CD28 and CTLA-4 differentially engage CD80, CD86, and PD-L1 at the cell surface remain incompletely understood. High-resolution structures of the CD28-CD80, CD28-CD86, CD28-CD80-PD-L1, and CTLA-4-CD80-PD-L1 complexes remain undetermined. Furthermore, while crystal structures of the soluble ectodomains have provided important insights into receptor-ligand recognition, these studies remove the proteins from their native membrane environment ^12–16^. Increasing evidence indicates that membrane confinement, lateral diffusion, and receptor clustering critically influence signal initiation and strength at the immunological synapse. In particular, CTLA-4 is predicted to form higher-order assemblies and mediate transendocytosis of CD80 and CD86, processes that cannot be readily explained by isolated, solution-state structures alone ^13,14,20–24^.

In this study, we used a lipid monolayer-based cryo-electron microscopy (cryo-EM) approach to determine the structures of CD28 and CTLA-4 bound to CD80, CD86, and PD-L1 under two-dimensional membrane confinement ^25^. This strategy enables the visualization of receptor-ligand complexes in a geometry that closely mimics the cell surface, revealing distinct stoichiometries, clustering behaviors, and higher-order assemblies that are not observed in solution. Our structures uncover how CD28 and CTLA-4 engage the same ligands in fundamentally different architectural modes, how CTLA-4 drives linear and two-dimensional lattice formation with CD80, and how PD-L1 binding selectively disrupts these assemblies without impairing CD28 engagement. Together, these findings provide a structural framework for understanding signal integration at the immunological synapse and offer new principles for the rational design of immunomodulatory therapeutics.

## RESULTS

### Structure of the CD28-CD86 complex

To determine the structure of the CD28-CD86 complex in a membrane-bound state, we employed a lipid monolayer method ^26–28^. In this approach, octa-histidine-tagged ectodomains of receptor-ligand complexes are tethered to a lipid monolayer containing Ni^2+^-NTA lipids (Figure S1A). Using this method, we have recently determined high-resolution structures of four TNF-TNFR family complexes assembled on a lipid monolayer ^25^. The lipid monolayer profoundly influences protein assembly by promoting lateral interactions between proteins and suppressing the formation of physiologically irrelevant three-dimensional assemblies (see DISCUSSION).

For structure determination of the CD28-CD86 complex, an octa-histidine tag was fused to the C-terminus of the CD86 ectodomain (Figure 1A, Figure S1B, and Table S1). The CD28-CD86 complex was assembled by mixing purified ectodomains of CD28 and CD86, which were subsequently tethered to a Ni^2+^-NTA lipid monolayer. The lipid layer was then transferred onto an EM grid, and cryo-EM data were collected (Figure 1B). The refined cryo-EM map revealed that CD28 and CD86 homodimers assemble into a complex adopting a “cross-leg” configuration (Figure 1C), in which the left CD86 monomer crosses over the right monomer. The three-dimensional structure of the CD28-CD86 complex was determined by initially fitting atomic models of CD28 and CD86 into the cryo-EM density map, followed by refinement to an overall resolution of 4.0 Å (Table S2). The initial atomic models of CD28 and CD86 were derived from the crystal structure of a CD28-antibody complex (PDB ID: 1YJD) ^12^ and a CTLA-4-CD86 complex (this study; described in the next section), respectively.

**Figure 1.**
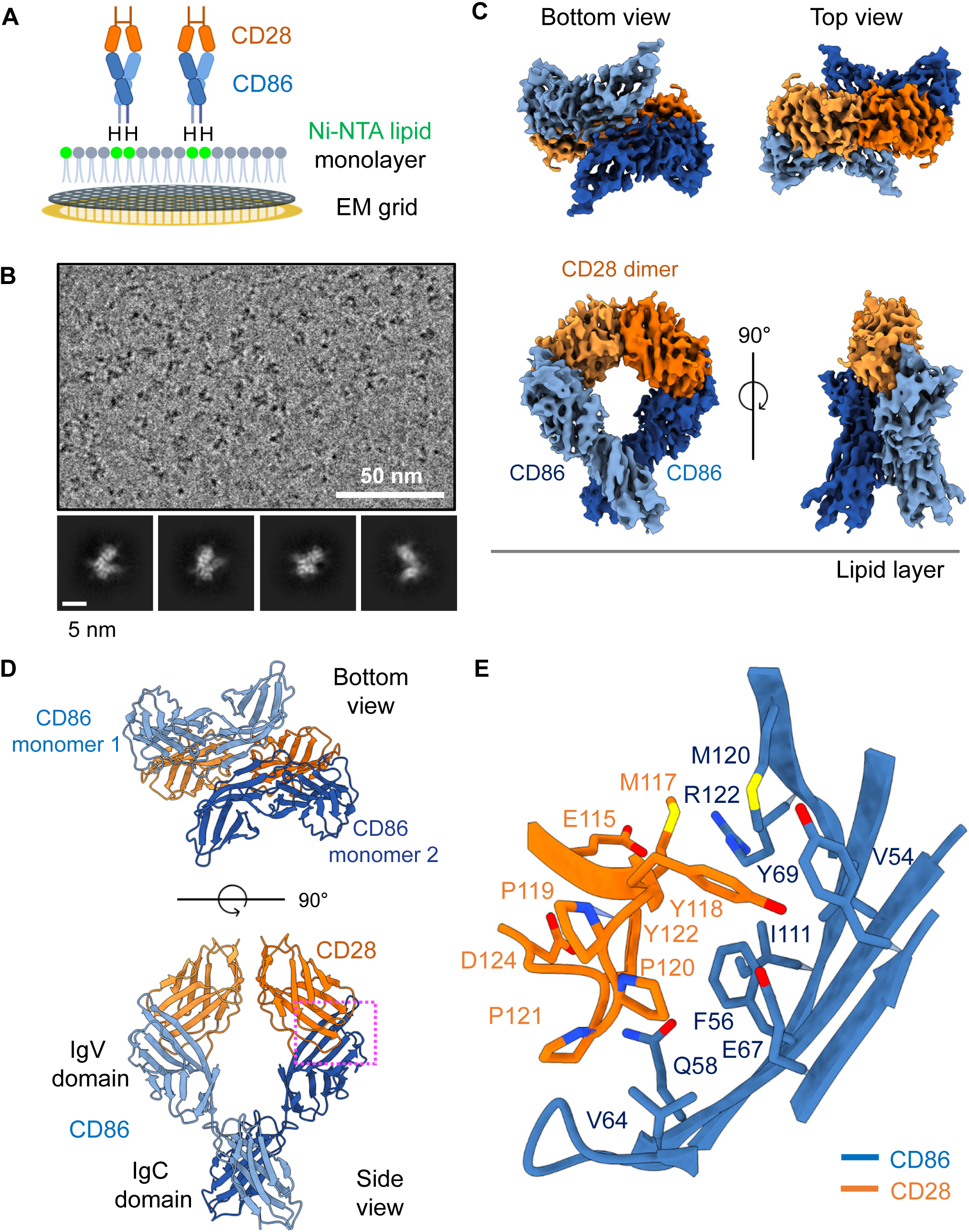
Structure of the CD28-CD86 complex on the lipid layer. (A) Schematic of the lipid monolayer experiment. An octa-histidine tag was fused to the C-terminus of the CD86 ectodomain, enabling tethering of the CD28-CD86 complex to the Ni^2+^-NTA lipid monolayer. “H” denotes the octa-histidine tag, and Ni^2+^-NTA lipids are shown in green. (B) Representative cryo-electron microscopy (cryo-EM) micrograph, with 2D class averages shown below. (C) Cryo-EM density map of the CD28-CD86 complex. CD28 is shown in dark and light orange, whereas CD86 is shown in dark and light blue. (D) Overall structure of the CD28-CD86 complex. (E) Close-up view of the CD28-CD86 heterodimerization interface. The region outlined by the dashed pink box in (D) is enlarged.

The ectodomain of CD86 consists of an immunoglobulin V-type domain linked to a C-type domain (Figure S1B), whereas the ectodomain of CD28 contains a single V-type immunoglobulin domain. In the CD28-CD86 structure, both CD28 and CD86 adopt homodimeric arrangements (Figure 1C and D). Homodimerization of CD28 is mediated by a hydrophobic interface formed by residues L59, I109, and I132 (Figure S2A and B). This hydrophobic interaction is further stabilized by a hydrogen bond between the imidazole ring of H134 and the backbone carbonyl oxygen of T107. The conserved disulfide bond connecting the two monomers in the CD28 homodimer is located near the C-terminal end of the ectodomain and is not visible in the electron density map, presumably due to structural flexibility (Figure S2A). The homodimeric structure of CD28 remains unchanged upon CD86 binding and can be superimposed with that observed in the CD28-antibody complex previously determined by X-ray crystallography (Figure S2C) ^12^. Homodimerization of CD86 is mediated by edge-to-edge interactions between the D β-strands of adjacent CD86 monomers (Figure S3A). This homodimeric interface is small, involving backbone atoms of only three amino acids, and appears to be unstable in the absence of bound CD28. The CD86 monomer itself adopts a rigid conformation that is not perturbed by CD28 binding and can be superimposed with CD86 in the CTLA-4-CD86 complex determined by X-ray crystallography (Figure S3B)^14^.

The binding between CD28 and CD86 is mediated exclusively by the IgV-type immunoglobulin domains of the two proteins. The conserved “MYPPPY” motif in CD28 plays a central role in the CD28-CD86 interaction (Figure 1E and S2A). The three prolines within this motif adopt a *cis-trans-cis* backbone conformation and form a loop connecting the F and G β-strands. This FG loop corresponds to the CDR3 loop, which plays a major role in antibody-antigen recognition in immunoglobulins. Within this loop, P120 and P121 form hydrophobic interactions with F56 and V64 of CD86. The preceding residues, M117 and Y118, engage in hydrophobic interactions with V54, F56, Y69, and M120 of CD86, and Y118 additionally forms a hydrogen bond with E67 of CD86. The residue Y122, located immediately downstream of the “MYPPPY” motif, makes hydrophobic contacts with F56 and I111 of CD86. In addition, E115 of CD28 forms an electrostatic interaction with R122 of CD86, further stabilizing the CD28-CD86 complex.

The MYPPPY motif is highly conserved in both CD28 and CTLA-4, and several site-directed mutations within this motif disrupt CD28 binding to CD86 (Figure S2A) ^29^. Likewise, mutations of CD86 residues F56, Q58, and V64 completely abolish CD86 binding to CD28 ^30^. The CD28 D124 mutation, most commonly to a valine or glutamate, is a recurrent activating alteration identified in angioimmunoblastic T-cell lymphoma (AITL) ^31^. This mutation enhances CD28-mediated co-stimulatory signaling and leads to increased downstream NF-κB activation, thereby promoting aberrant T-cell activation. Substitution at D124 has been shown to significantly increase binding affinity for CD86. The D124 residue is located adjacent to the CD28-CD86 binding interface, and although it is not directly involved in ligand contact, the mutation likely enhances electrostatic complementarity ^31^ or indirectly increases ligand affinity by altering the conformation of the conserved MYPPPY loop (Figure 1E).

### Structure of the CTLA-4-CD86 complex

To determine the structure of the CTLA-4-CD86 complex, an octa-histidine tag was fused to the C-terminal region of the CD86 ectodomain (Figures 2A and S1B, and Table S1). The purified ectodomains of CTLA-4 and CD86 were mixed, and the assembled complex was tethered to a Ni^2+^-NTA lipid monolayer. The lipid layer was then transferred onto an EM grid, and cryo-EM data were collected (Figure 2B). The refined cryo-EM map revealed that one CTLA-4 homodimer binds two CD86 monomers in an “open-leg” configuration (Figure 2C). The three-dimensional structure of the CTLA-4-CD86 complex was determined by initially fitting atomic models of CTLA-4 and CD86 into the cryo-EM density map, followed by refinement to an overall resolution of 3.2 Å (Table S2). The initial atomic models of CTLA-4 and CD86 were derived from a previously reported crystal structure of the CTLA-4-CD80 complex (PDB ID: 1I8L) and a model predicted by AlphaFold3, respectively ^13,32^.

**Figure 2.**
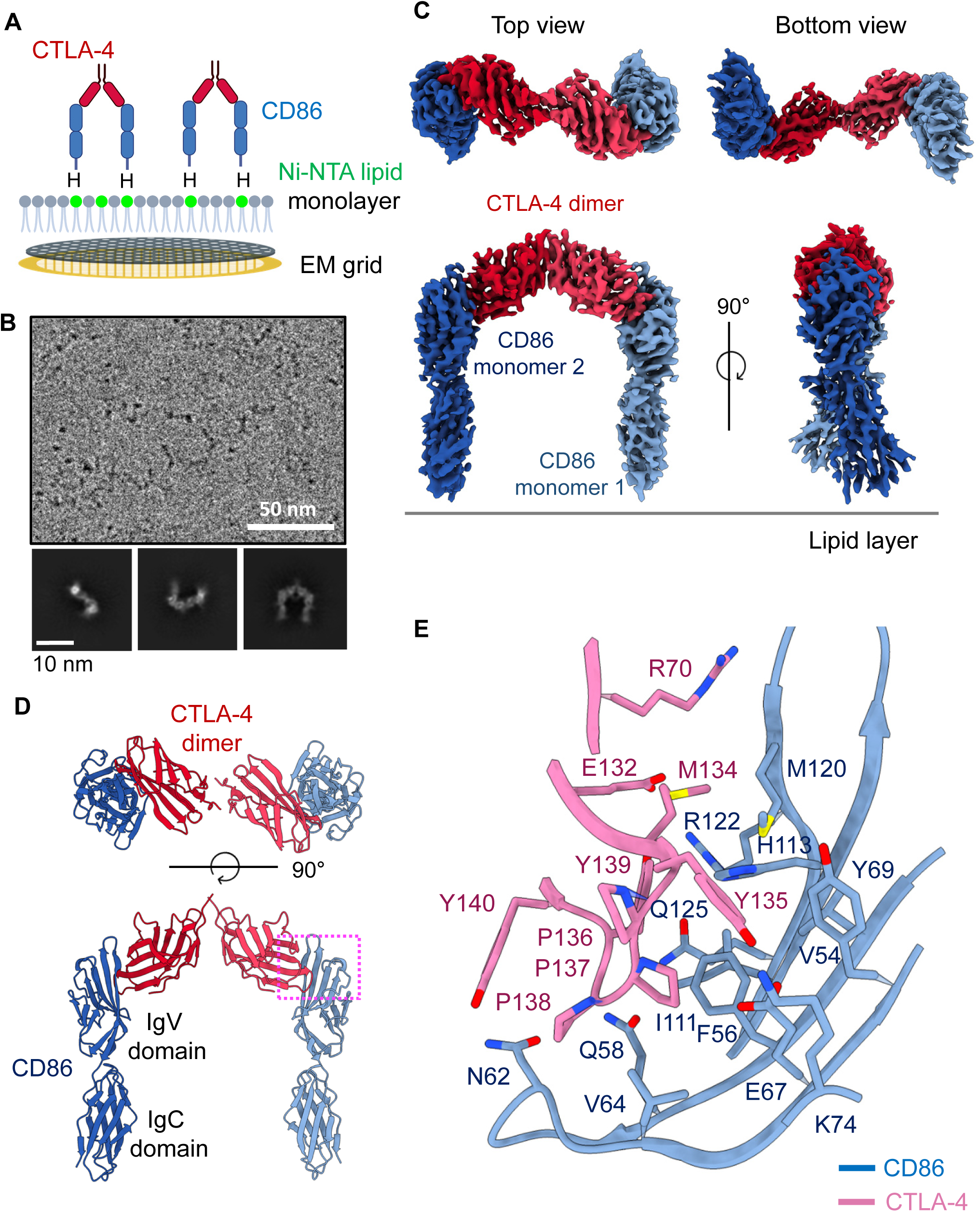
Structure of the CTLA-4-CD86 complex on the lipid layer. (A) Schematic of the lipid monolayer experiment. An octa-histidine tag was fused to the C-terminus of the CD86 ectodomain, enabling tethering of the CTLA-4-CD86 complex to the Ni^2+^-NTA lipid monolayer. “H” denotes the octa-histidine tag, and Ni^2+^-NTA lipids are shown in green. (B) Representative cryo-EM micrograph, with 2D class averages shown below. (C) Cryo-EM density map of the CTLA-4-CD86 complex. CTLA-4 is shown in dark and light red, whereas CD86 is shown in dark and light blue. (D) Overall structure of the CTLA-4-CD86 complex. (E) Close-up view of the CTLA-4-CD86 heterodimerization interface. The region outlined by the dashed pink box in (D) is enlarged.

In the CTLA-4-CD86 structure, two CD86 monomers bind to a CTLA-4 homodimer, yielding a 2:2 stoichiometry (Figure 2D). Hydrophobic residues V45, L47, Y150, and I152 play key roles in stabilizing the CTLA-4 homodimer (Figure S4A). As observed for CD28, the cysteine residues that form inter-subunit disulfide bonds in CTLA-4 are located within a structurally flexible region near the C-terminal end of the ectodomain and are not resolved in the electron density map (Figure S2A). Unlike the CD28-CD86 complex, the two CD86 monomers in the CTLA-4-CD86 complex are widely separated and do not make direct contact with one another (Figure 2D). This separation does not arise from structural changes in CD86, as the CD86 structures in the CTLA-4-CD86 and CD28-CD86 complexes superimpose with a Cα RMSD of 1.12 Å (Figure S4B). The interaction between the IgV and IgC domains of CD86 is mediated primarily by hydrophobic residues F32, I102, A134, Y163, and L196. An ionic interaction between K103 and E195 further stabilizes this hydrophobic interface (Figure S4C). The CTLA-4 homodimer in our cryo-EM structure closely matches that observed in the previously reported crystal structure of the CTLA-4-CD86 complex, with a Cα RMSD of 1.52 Å (Figure S5A). Similarly, the individual IgV domains of CD86 in the cryo-EM and X-ray structures adopt nearly identical conformations and superimpose with a Cα RMSD of 0.69 Å (Figure S5B). In contrast, the relative orientation of CD86 with respect to CTLA-4 is rotated by ∼7.4° (Figure S5C). This small but noticeable structural alteration is likely due to the attachment of CD86 to the lipid membrane.

Similar to the CD28-CD86 interface, the conserved MYPPPY motif in CTLA-4 plays a central role in mediating the CTLA-4-CD86 interaction (Figure 2E). Within this loop, residues M134, P137, P138, and Y139 of CTLA-4 form extensive hydrophobic interactions with residues V54, F56, V64, Y69, I111, and M120 of CD86. This hydrophobic core is further stabilized by surrounding polar interactions: Y135 of CTLA-4 contacts K74 and E67 of CD86, while the backbone atoms of Y139 in CTLA-4 and Q58 in CD86 form strong hydrogen bonds. In addition, Y140 of CTLA-4 interacts with N62 of CD86, and E132 and Y139 of CTLA-4 engage in hydrophilic interactions with R122 and Q125 of CD86. Consistent with these structural observations, site-directed mutagenesis of CD86 residues F56, Q58, or V64 completely abolishes CD86 binding to CTLA-4, while several mutations within the MYPPPY motif of CTLA-4 completely disrupt its interaction with CD86 ^29,30,33–36^. R70 of CTLA-4 is located in the interaction interface, and its mutation to a glutamine reduces binding affinity tenfold ^33^ (Figure 2E).

### Structure of the CD28-CD80 complex

For structure determination of the CD28-CD80 complex, an octa-histidine tag was fused to the C-terminal region of the CD80 ectodomain (Figures 3A, S1B, and Table S1). The purified ectodomains of CD28 and CD80 were mixed, and the assembled complex was tethered to a Ni²⁺-NTA lipid monolayer. The lipid layer was then transferred onto an EM grid, and images were collected by cryo-EM (Figure 3B). The refined cryo-EM map revealed that two CD28 homodimers and one CD80 homodimer assemble into a complex with a “closed-leg” configuration (Figure 3C). The three-dimensional structure of the CD28-CD80 complex was determined by fitting atomic models of CD28 and CD80 into the cryo-EM density map, followed by refinement to an overall resolution of 3.9 Å (Table S2). The initial atomic models of CD28 and CD80 were derived from the crystal structures of a CD28-antibody complex (PDB ID: 1YJD) and a CTLA-4-CD80 complex (PDB ID: 1I8L), respectively ^12,13^. Notably, shifting the octa-histidine membrane anchoring site from CD80 to CD28 did not alter the overall architecture of the complex (Figure S6), indicating that complex formation is an intrinsic property of the protein assembly under two-dimensional membrane confinement.

**Figure 3.**
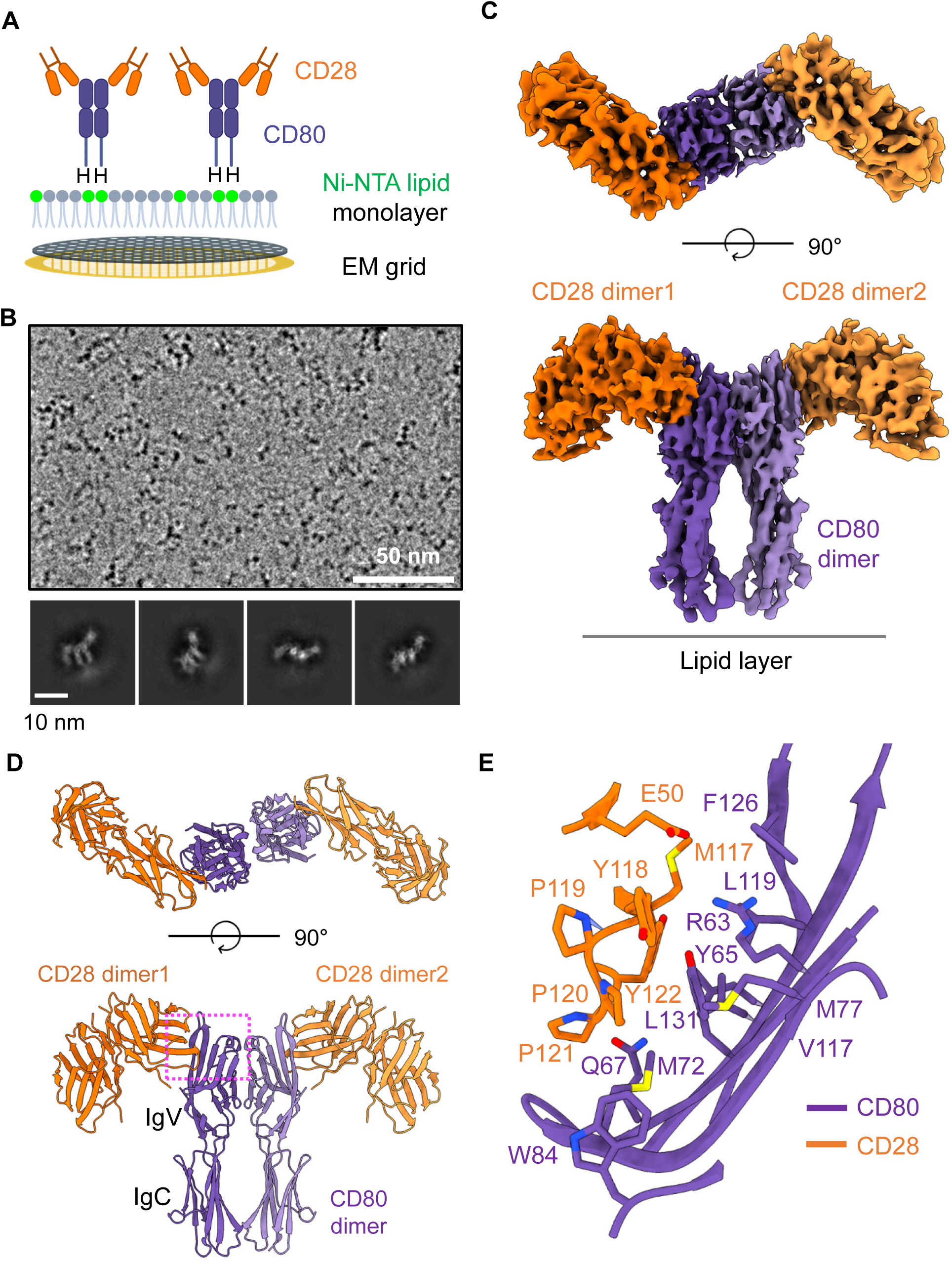
Structure of the CD28-CD80 complex on the lipid layer. (A) Schematic of the lipid monolayer experiment. An octa-histidine tag was fused to the C-terminus of the CD80 ectodomain, enabling tethering of the CD28-CD80 complex to the Ni^2+^-NTA lipid monolayer. “H” denotes the octa-histidine tag, and Ni^2+^-NTA lipids are shown in green. (B) Representative cryo-EM micrograph, with 2D class averages shown below. (C) Cryo-EM density map of the CD28-CD80 complex. CD28 is shown in dark and light orange, whereas CD80 is shown in dark and light purple. (D) Overall structure of the CD28-CD80 complex. The IgV and IgC domains of CD80 are labeled. (E) Close-up view of the CD28-CD80 heterodimerization interface. The region outlined by the dashed pink box in (D) is enlarged.

In the CD28-CD80 structure, two CD28 homodimers bind to a single CD80 homodimer, resulting in a complex with a 4:2 molar ratio of CD28 to CD80 (Figure 3C and D). This multimeric CD28-CD80 complex cannot extend into a linear cluster, unlike the CTLA-4-CD80 complex (see next section), because the inter-subunit angle of the CD28 dimer prevents engagement of additional CD80 subunits due to steric clashes. The structure of the CD28 homodimer in the CD28-CD80 complex is essentially identical to that in the CD28-CD86 complex, and the two structures can be superimposed with a Cα RMSD of 1.08 Å (Figure S7A). Homodimerization of CD80 is mediated by two hydrophobic interfaces within the IgV-type immunoglobulin domain. The first interface is formed by residues V56, L59, M76, and I95 (Figure S7B), whereas the second interface, comprising residues V45, I92, V102, and L104, stabilizes the primary interface and supports dimer formation. The resulting CD80 homodimer closely matches the previously reported crystal structure (PDB ID: 1I8L) (Figure S8), and the two structures can be superimposed with a Cα RMSD of 1.25 Å ^13^. Notably, CD80 in the crystal structure is bound to CTLA-4 rather than CD28. Together, these observations demonstrate that the CD80 homodimer is structurally rigid and is not readily perturbed by different binding partners or crystallization conditions.

Similar to the CD28-CD86 complex, the conserved “MYPPPY” motif in CD28 plays a central role in mediating the CD28-CD80 interaction (Figure 3E). Within this loop, residues M117, Y118, P120, P121, and Y122 form extensive hydrophobic interactions with residues Y65, M72, M77, W84, V117, L119, F126, and L131 of CD80. P119 within the “MYPPPY” motif plays an important role in stabilizing the conformation of the loop but does not make direct contact with CD80. The backbone atoms of Y122 form a strong hydrogen bond with the side chain of Q67 in CD80, further stabilizing the hydrophobic interface. In addition, E50, located in a loop adjacent to the “MYPPPY” motif, forms a charge interaction with R63 of CD80, further stabilizing the CD28-CD80 complex. Consistent with these structural observations, mutations in the MYPPPY motif of CD28 or in CD80 residues R63, Y65, Q67, M72, and W84 completely abolished complex formation ^29,30,37^ (Figure 3E).

### Structure of the CTLA-4-CD80 linear and two-dimensional clusters

For structure determination of the CTLA-4-CD80 complex under membrane-confined conditions, an octa-histidine tag was fused to the C-terminal region of the CD80 ectodomain (Figures 4A, S1B, and Table S1). The purified ectodomains of CTLA-4 and CD80 were mixed, and the assembled complex was tethered to a Ni^2+^-NTA lipid monolayer. The lipid monolayer was transferred onto an EM grid, and cryo-EM data were collected (Figure 4B). Raw cryo-EM micrographs and corresponding 2D class averages revealed the emergence of prominent linear clusters of CTLA-4-CD80 complexes on the lipid membrane (Figure 4B and C). The refined three-dimensional reconstruction shows that CTLA-4 homodimers engage CD80 homodimers in a 2:2 molar ratio (Figure 4D and E). Atomic models of CTLA-4 and CD80 were fitted into the cryo-EM density map and refined to an overall resolution of 3.8 Å (Table S3). Initial models were derived from the previously reported crystal structure of the CTLA-4-CD80 complex (PDB ID: 1I8L) ^13^. The repeating unit, consisting of one CTLA-4 dimer bound to two CD80 monomers, spans approximately 98 Å (Figure 4E). Homodimeric interactions between adjacent CD80 molecules mediate lateral assembly of these repeating units, giving rise to extended linear arrays that typically comprise several tens of units. Notably, shifting the octa-histidine membrane anchoring site from CD80 to CTLA-4 did not alter the overall architecture of the linear cluster (Figure S9), indicating that clustering is an intrinsic property of the protein assembly under two-dimensional membrane confinement.

**Figure 4.**
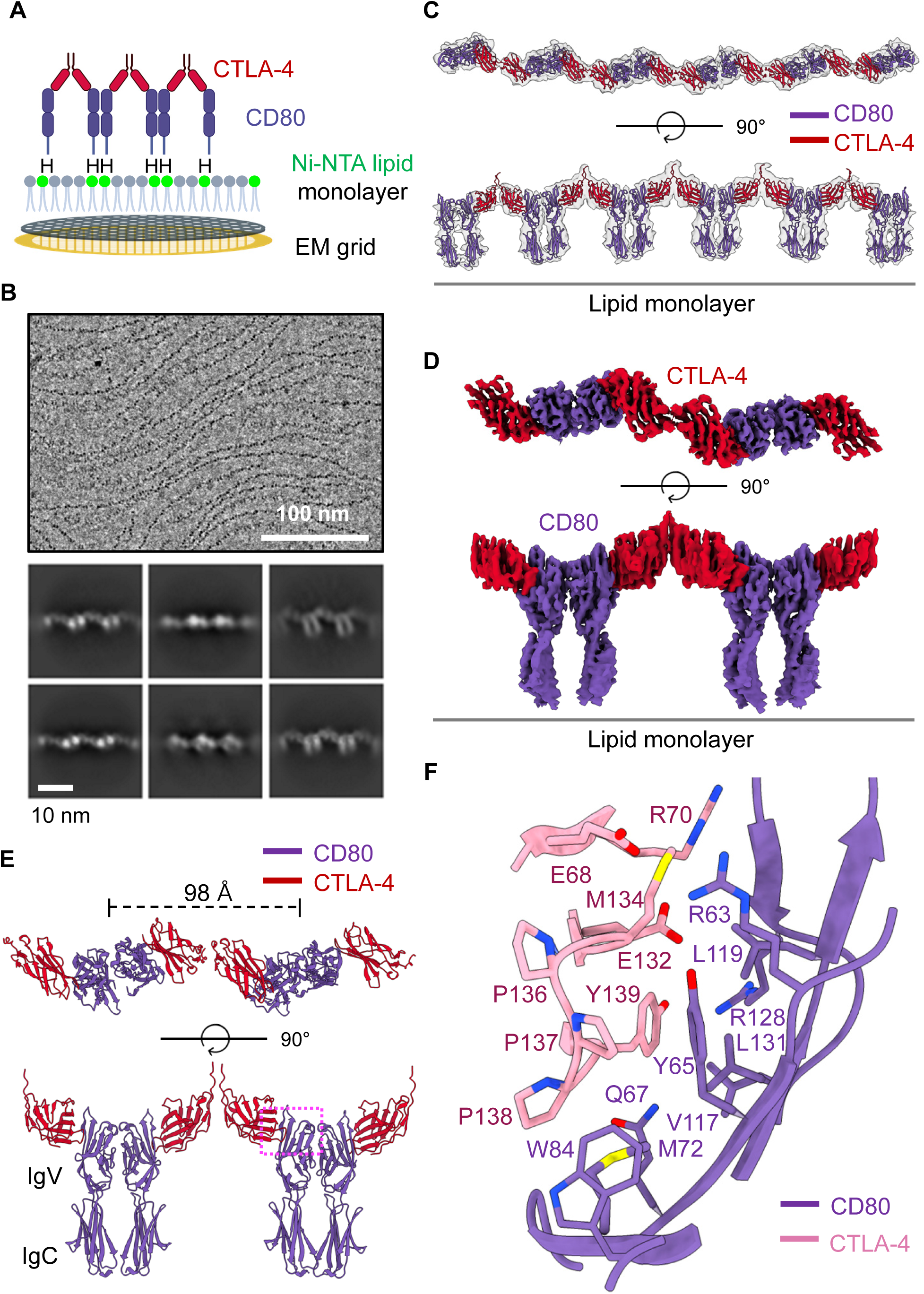
Linear clustering of the CTLA-4-CD80 complex on the lipid layer. (A) Schematic of the lipid monolayer experiment. An octa-histidine tag was fused to the C-terminus of the CD80 ectodomain, enabling tethering of the CTLA-4-CD80 complex to the Ni^2+^-NTA lipid monolayer. “H” denotes the octa-histidine tag, and Ni^2+^-NTA lipids are shown in green. (B) Representative cryo-EM micrograph, with 2D class averages shown below. (C) Cryo-EM density map of the CTLA-4-CD80 linear cluster, shown in grey. Refined atomic models of the CTLA-4-CD80 complex are fitted into the map. CTLA-4 is shown in red, whereas CD80 is shown in purple. (D) Three-dimensional refined cryo-EM map of the CTLA-4-CD80 linear cluster. (E) Overall structure of the CTLA-4-CD80 complex within the linear cluster. The IgV and IgC domains of CD80 are labeled. (F) Close-up view of the CTLA-4-CD80 heterodimerization interface. The region outlined by the dashed pink box in (E) is enlarged.

Formation of the linear cluster is not due to structural changes in either the CTLA-4 or CD80 homodimers. The homodimeric structure of CTLA-4 within the CTLA-4-CD80 linear cluster is essentially identical to that observed in the CTLA-4-CD86 complex; the two structures can be superimposed with a Cα RMSD of 0.44 Å (Figure S10A). The homodimerization interfaces are indistinguishable, and the same amino acid residues mediate CTLA-4 homodimerization in both complexes (Figure S4A). Likewise, the immunoglobulin domains of CD80 adopt nearly identical conformations across complexes. The CD80 homodimers in the CTLA-4-CD80 and CD28-CD80 complexes can be superimposed with a Cα RMSD of 1.00 Å (Figure S10B). Consistent with this observation, the same hydrophobic residues, V45, V56, L59, M76, I92, I95, V102, and L104, that mediate CD80 homodimerization in the CD28-CD80 complex also mediate CD80 homodimerization within the CTLA-4-CD80 linear cluster (Figure S7B).

Similar to the heterodimerization interface observed in the CTLA-4-CD86 structure, the conserved “MYPPPY” motif of CTLA-4 plays a central role in mediating the CTLA-4-CD80 interaction (Figure 4F). Within this loop, residues M134, P137, P138, and Y139 of CTLA-4 form extensive hydrophobic interactions with residues Y65, M72, V117, L119, and L131 of CD80. This hydrophobic core is further stabilized by surrounding hydrophilic interactions. Specifically, the backbone of CTLA-4 Y139 forms a hydrogen bond with the side chain of CD80 Q67. Additionally, strong salt bridges are formed between CTLA-4 E68 and CD80 R63, as well as between CTLA-4 E132 and CD80 R128. Consistent with these structural observations, mutations in key residues, including the MYPPPY motif of CTLA-4 and CD80 residues R63, Y65, Q67, M72, and W84, completely abolish CD80 binding to CTLA-4 ^29,30,33,36,37^. Interestingly, mutation of Y135 within the MYPPPY motif of CTLA-4 to phenylalanine has no effect on the CTLA-4-CD80 interaction but selectively abolishes CTLA-4 binding to CD86 ^34,35^. This residue is not involved in CTLA-4 binding to CD80; however, it plays an important role in CD86 binding by forming a hydrophobic interaction with Y69 of CD86 (Figure 2E). R70 of CTLA-4 is located in the interaction interface, and its mutation to a glutamine reduces binding affinity by twenty-five fold ^33^ (Figure 4F).

CTLA-4 haploinsufficiency causes an autosomal dominant immune dysregulation syndrome characterized by autoimmunity and lymphoproliferation, including lymphadenopathy and hepatosplenomegaly, reflecting impaired negative regulation of T cell activation ^38–41^. Immunologically, patients with CTLA-4 haploinsufficiency show reduced CTLA-4 expression and defective transendocytosis, suggesting disruption of the CTLA-4-CD80 interaction. Our structural analysis supports this genetic observation, as several disease-associated mutations map to the CTLA-4-CD80 interaction interface. These include the R70W, P136A/S/L, P137R/L, and Y139C mutations. R70 is located adjacent to the ligand-binding interface, whereas P136, P137, and Y139 are integral components of the conserved MYPPPY motif (Figure 4F).

Linear clustering of the CTLA-4-CD80 complex was previously proposed based on crystal-packing interactions observed in the crystal structure of the complex. In that structure, the asymmetric unit contains one CD80 homodimer bound to two CTLA-4 monomers, and crystal packing interactions between CTLA-4 molecules from neighboring asymmetric units generate an apparent linear arrangement of CTLA-4-CD80 complexes. Although the crystal-packing-derived model and our membrane-tethered cryo-EM structure share a similar overall linear architecture, direct comparison reveals notable differences. Superposition of the first CD80 molecules from each structure reveals relative rotations of approximately 8° (Figure S11A). These deviations arise from subtle adjustments at the CTLA-4-CD80 heterodimerization interface (Figure S11B), reflecting the distinct geometric constraints imposed by two-dimensional membrane confinement versus three-dimensional crystal packing.

At lower protein concentrations, most CTLA-4-CD80 complexes assemble into linear clusters. However, at higher protein concentrations on the lipid layer, these linear clusters laterally associate and stack to form a two-dimensional lattice (Figure 5A). Each lattice typically comprises multiple linear clusters. Two-dimensional class averages from cryo-EM reveal that the lattice adopts a highly ordered architecture (Figure 5B). A three-dimensional reconstruction of the lattice, calculated to an overall resolution of 7.7 Å, shows that CTLA-4 molecules from neighboring linear clusters form close lateral contacts that mediate assembly of the two-dimensional structure (Figure 5C). This interface involves a limited number of residues, including S108, G109, and N110 in the DE loop of CTLA-4 (Figure 5D and E). Although these lateral interactions are individually weak and not detectable in solution, their cooperative engagement is promoted by two-dimensional membrane confinement, which restricts diffusion and thereby stabilizes the lattice.

**Figure 5.**
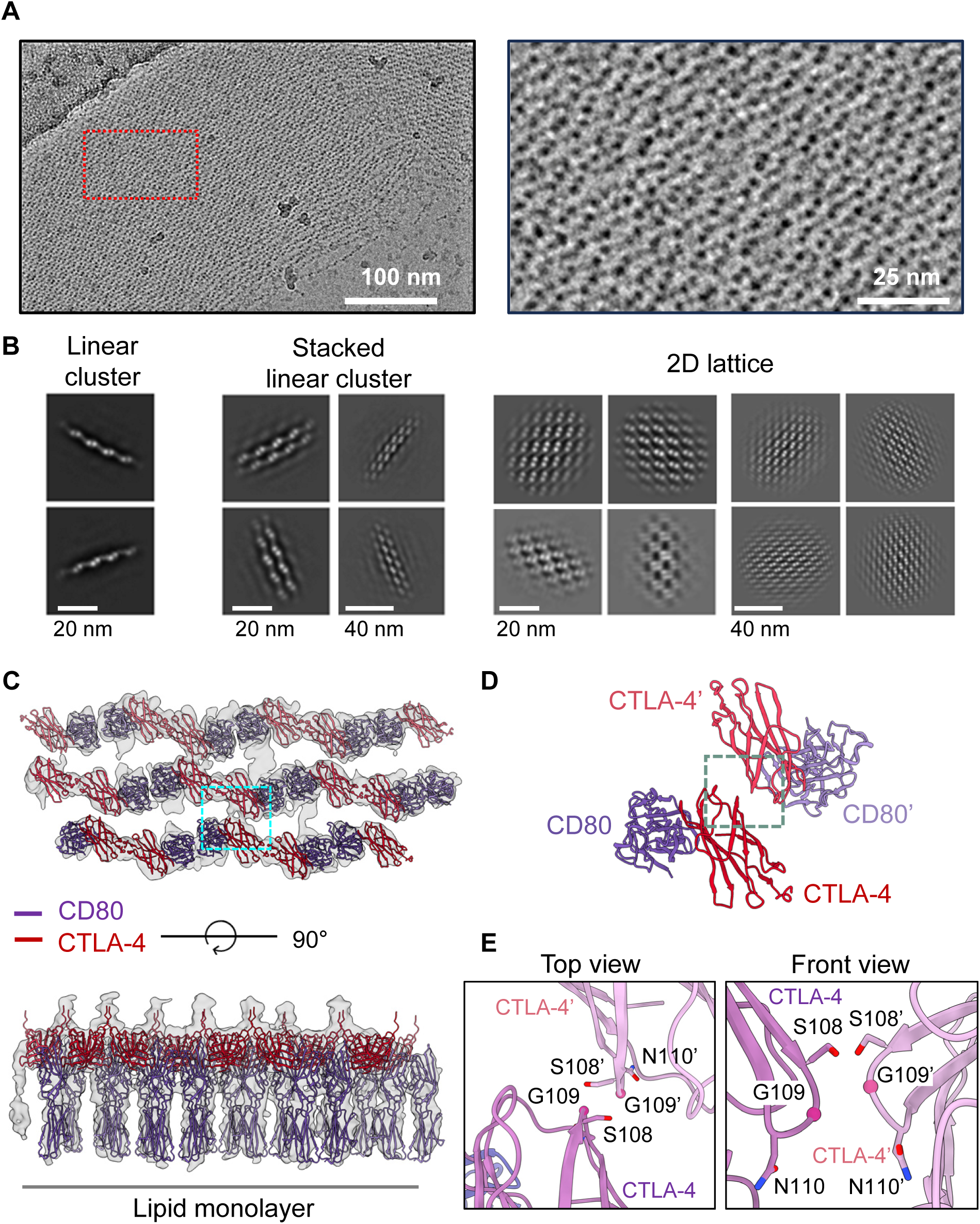
Formation of a two-dimensional lattice of the CTLA-4-CD80 complex on the lipid layer. (A) Representative cryo-EM micrograph of the CTLA-4-CD80 lattice. The region outlined by the red box is enlarged in the right panel. (B) 2D class averages of the CTLA-4-CD80 linear clusters and two-dimensional lattices. (C) Cryo-EM density map of the CTLA-4-CD80 lattice, with refined atomic models of the CTLA-4-CD80 complex fitted into the map. CTLA-4 is shown in red, whereas CD80 is shown in purple. (D) Close-up view of the CTLA-4-CD80 complex within the two-dimensional lattice. The region outlined by the dashed cyan box in (C) is enlarged. The second CTLA-4 and CD80 monomers are indicated by primes. (E) Close-up view of the CTLA-4-CTLA-4 interaction interface within the CTLA-4-CD80 lattice. The region outlined by the dashed green box in (D) is enlarged. Residue numbers for the second CTLA-4 monomer are indicated by prime symbols.

### Structure of the CTLA-4-CD80 complex in a cellular membrane environment

To test whether CTLA-4-CD80 clusters form in a cellular membrane environment, HEK293 cells were transfected with a recombinant baculovirus encoding full-length CTLA-4 (CTLA-4fl). Following gentle cell lysis in a hypotonic buffer, a CD80 ectodomain bearing an octa-histidine tag was added to the membrane preparation (Figure S12). After removal of unbound CD80, membrane fragments containing flat membrane sheets and vesicles were generated by sonication. Membrane fragments bearing bound CD80 were then tethered to a Ni^2+^-NTA lipid monolayer for cryo-EM imaging (Figure 6A). The lipid monolayer serves two purposes in this approach: first, it mimics the antigen-presenting cell membrane by displaying CD80 on its surface; second, it selectively enriches membrane fragments containing CTLA-4fl-CD80 complexes, thereby facilitating cryo-EM analysis.

**Figure 6.**
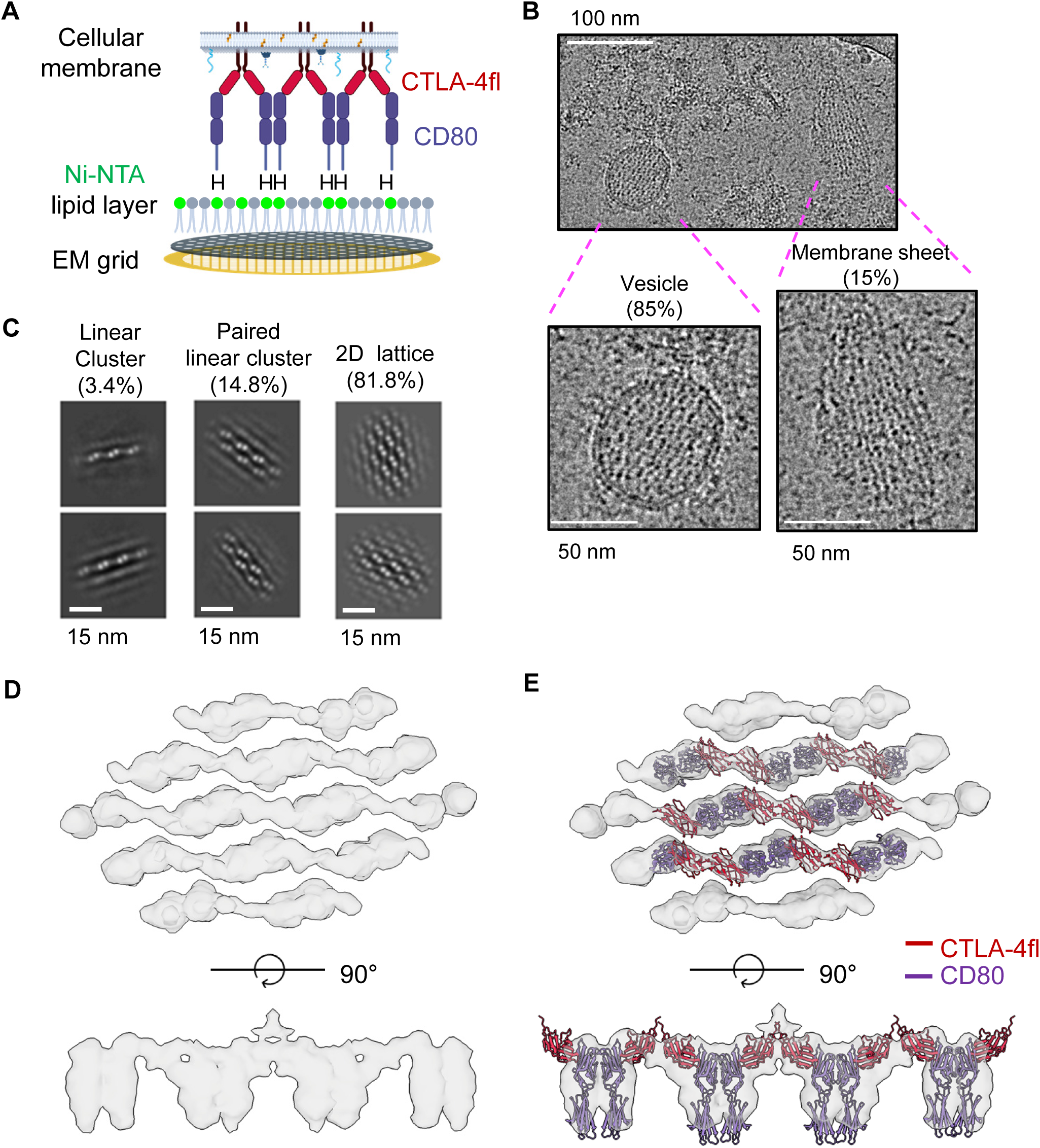
Formation of linear and two-dimensional clusters of the CTLA-4fl-CD80 complex on cellular membrane fragments. (A) Schematic of the lipid monolayer experiment using cellular membrane fragments. An octa-histidine tag was fused to the C-terminus of the CD80 ectodomain, enabling the tethering of the CTLA-4fl-CD80 complex to the Ni²⁺-NTA lipid monolayer. “H” denotes the octa-histidine tag, and the Ni²⁺-NTA lipids are shown in green. Other proteins, carbohydrates, and cholesterols on the cellular membrane fragments are schematically shown as cyan, blue, and orange lines, respectively. (B) Representative cryo-EM micrograph of the CTLA-4fl-CD80 lattice. The regions outlined by magenta lines are enlarged in the lower panels. (C) 2D class averages of linear CTLA-4fl-CD80 clusters and two-dimensional lattices. (D) Cryo-EM density map of the CTLA-4fl-CD80 lattice. (E) Structure of the CTLA-4fl-CD80 lattice on the cellular membrane fragment. The atomic model of the CTLA-4-CD80 complex is fitted into the cryo-EM density map. CTLA-4 is shown in red, whereas CD80 is shown in purple.

As shown in Figures 6B and C, the raw micrographs and 2D class averages clearly reveal the formation of 2D lattices of CTLA-4fl-CD80 ectodomain complexes. Most CTLA-4fl-CD80 clusters were observed within vesicle-like membrane structures (Figure 6B). It remains unclear whether these structures represent closed spherical vesicles or membrane sheets curled at the edges. The vast majority of observed CTLA-4fl-CD80 clusters adopted the 2D lattice organization, whereas linear-type clusters were rarely detected (Figure 6C). A 3D reconstruction of the cryo-EM map demonstrates that the structure of the CTLA-4fl-CD80 cluster is essentially identical to that formed on artificial lipid monolayers using only the CTLA-4 and CD80 ectodomains (Figures 6D and E; Figure S13). This indicates that the formation of the CTLA-4-CD80 cluster is driven primarily by ectodomain interactions between CTLA-4 and CD80, with other membrane proteins and lipids on the cell membrane having minimal impact on the overall structure.

### Disruption of the CTLA-4-CD80 linear cluster by PD-L1 binding

Previous studies have shown that PD-L1 engages in a *cis* interaction with CD80 when both proteins are expressed on the same cell membrane ^8–10,42^. This interaction has been proposed to weaken the CTLA-4-CD80 interaction by disrupting CTLA-4-CD80 clustering and to inhibit CTLA-4-mediated transendocytosis ^10^. To investigate the structural basis of PD-L1-mediated disruption of CTLA-4-CD80 clusters, the purified PD-L1 ectodomain containing an octa-histidine tag was mixed with CD80. CTLA-4 was subsequently added to the mixture to form the CTLA-4-CD80 complex. The C-terminus of the CD80 ectodomain was also tagged with an octa-histidine motif to enable the formation of CTLA-4-CD80 clusters on the lipid layer (Figure 7A). When PD-L1 was added to the layer at an equimolar ratio relative to the CTLA-4-CD80 complex, both linear and two-dimensional CTLA-4-CD80 clusters became shorter but remained largely intact (Figure 7B). In contrast, addition of PD-L1 at a threefold molar excess resulted in substantial disruption of CTLA-4-CD80 clusters, as observed in cryo-EM micrographs (Figure 7C). 2D class averages revealed that most CTLA-4-CD80 clusters were disassembled into isolated repeating units consisting of one CTLA-4 homodimer bound to two CD80-PD-L1 heterodimers. In addition, partially disrupted assemblies were observed, in which short linear CTLA-4-CD80 clusters persisted but were capped by PD-L1 molecules at their termini.

**Figure 7.**
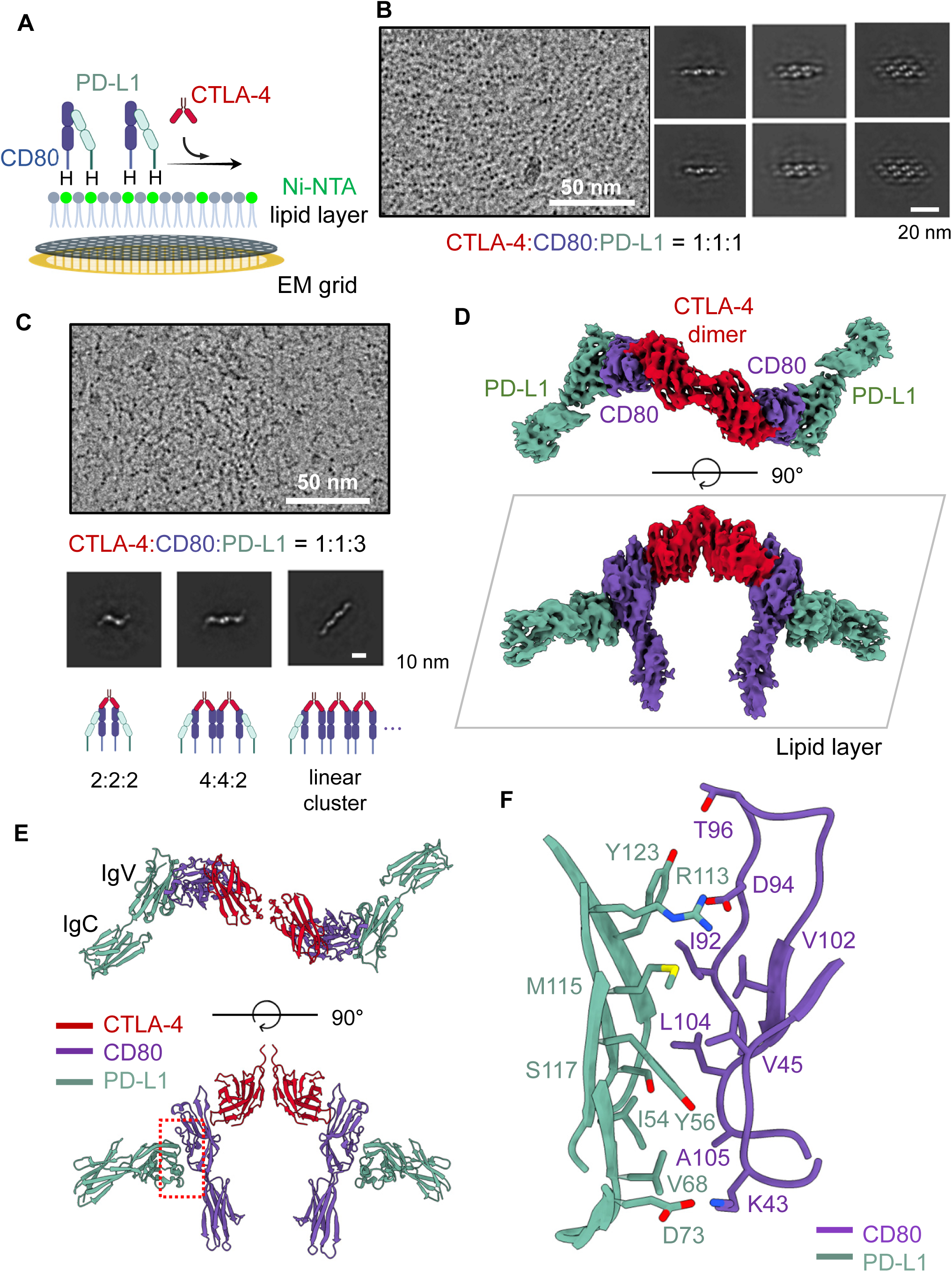
Structure of the CTLA-4-CD80-PD-L1 complex on a lipid layer. (A) Schematic of the lipid monolayer experiment. An octa-histidine tag was fused to the C-termini of the CD80 and PD-L1 ectodomains. Purified CD80 and PD-L1 were mixed and tethered to a Ni²⁺-NTA lipid monolayer, followed by the addition of CTLA-4 to form the final CTLA-4-CD80-PD-L1 complex. “H” denotes the octa-histidine tag, and the Ni²⁺-NTA lipids are shown in green. (B) Representative cryo-EM micrograph and 2D class averages of the CTLA-4-CD80-PD-L1 complex assembled at a 1:1:1 molar ratio. (C) Representative cryo-EM micrograph and 2D class averages of the CTLA-4-CD80-PD-L1 complex assembled at a 1:1:3 molar ratio. Schematic representations of the CTLA-4-CD80-PD-L1 complexes corresponding to each 2D class are shown below the class averages. (D) Cryo-EM density map of the CTLA-4-CD80-PD-L1 complex. CTLA-4, CD80, and PD-L1 are shown in red, purple and light green, respectively. The planar lipid layer is drawn as a grey parallelogram. (E) Overall structure of the CTLA-4-CD80-PD-L1 complex. The IgV and IgC domains of PD-L1 are labeled. (F) Close-up view of the CD80-PD-L1 interaction interface. The region outlined by the dashed red box in (E) is enlarged.

The cryo-EM map, refined to 3.7 Å resolution, reveals that PD-L1 binding disrupts CD80 homodimerization and that a CD80 monomer can simultaneously engage PD-L1 and CTLA-4, forming a 2:2:2 complex of CTLA-4, CD80, and PD-L1 in an “open-leg” configuration (Figure 7D and Table S4). The interaction between PD-L1 and CD80 is mediated exclusively by their IgV-type immunoglobulin domains (Figure 7E). Hydrophobic residues V45, I92, V102, L104, and A105 of CD80 and residues I54, Y56, V68, and M115 of PD-L1 form the core of the heterodimerization interface (Figure 7F). This interface is further stabilized by hydrogen bonds between Y56 of PD-L1 and the backbone of K43 of CD80, as well as between Y123 of PD-L1 and T96 of CD80. Furthermore, electrostatic interactions, specifically between R113 of PD-L1 and D94 of CD80, and between D73 of PD-L1 and K43 of CD80, enhance PD-L1 binding to CD80. Notably, the binding site for CD80 on PD-L1 overlaps with that of PD-1, indicating direct competition between PD-1 and CD80 for PD-L1 binding (Figure S14). Structural comparison of our CTLA-4-CD80-PD-L1 cryo-EM structure with a previously reported CD80-PD-L1 complex determined in the absence of CTLA-4 ^43^ reveals an ∼7.7° rotation of PD-L1 relative to CD80 (Figure S15). In this crystal structure, CD80 has only the IgV domain and was engineered to contain seven mutations to enhance its affinity for PD-L1. Therefore, the small but noticeable difference in PD-L1-CD80 geometry may arise either from the engineered mutations or from the confinement of the two proteins to a flat lipid membrane.

### Structural changes in the CD28-CD80 complex induced by PD-L1 binding

To investigate the structural basis of PD-L1 binding to the CD28-CD80 complex, the purified PD-L1 ectodomain was mixed with CD80. CD28 was subsequently added to the mixture to form the ternary CD28-CD80-PD-L1 complex (Figure 8A). Octa-histidine tags were fused to the C-termini of both the PD-L1 and CD80 ectodomains to enable membrane anchoring. The electron density map and two-dimensional class averages indicate that the resulting complex adopts a more compact shape than the CD28-CD80 4:2 complex described in Figure 3 (Figure 8B). A three-dimensional map of the CD28-CD80-PD-L1 complex at 3.3 Å resolution reveals pronounced structural rearrangements relative to the CD28-CD80 complex (Figure 8C and Table S4). In the absence of PD-L1, CD28 engages CD80 with a 4:2 stoichiometry in a “closed-leg” configuration (Figure 3D). Upon PD-L1 binding, the complex undergoes a transition to a “reversed cross-leg” configuration, in which the left CD80 monomer crosses beneath the right CD80 monomer (Figure 8D). This rearrangement reduces the CD28:CD80 stoichiometry from 4:2 to 2:2, yielding a ternary 2:2:2 complex of CD28, CD80, and PD-L1. This structural transition appears to be driven by steric clashes between PD-L1 and the second CD80 monomer in the CD80 homodimer when the CD28-CD80 complex adopts the closed-leg configuration in the absence of PD-L1 (Figure 8E). Notably, this rearrangement does not involve substantial conformational changes within the individual protein components or their binding interfaces. The CD28 dimer and CD80 monomers in the CD28-CD80 and CD28-CD80-PD-L1 complexes can be superimposed with Cα RMSDs of 1.14 Å and 0.83 Å, respectively (Figure S16). In contrast, the relative orientations of CD80 with respect to CD28 and of PD-L1 with respect to CD80 are both rotated by ∼7.3°. This small but noticeable structural alteration is likely due to the simultaneous attachment of CD80 and PD-L1 to a flat lipid membrane.

**Figure 8.**
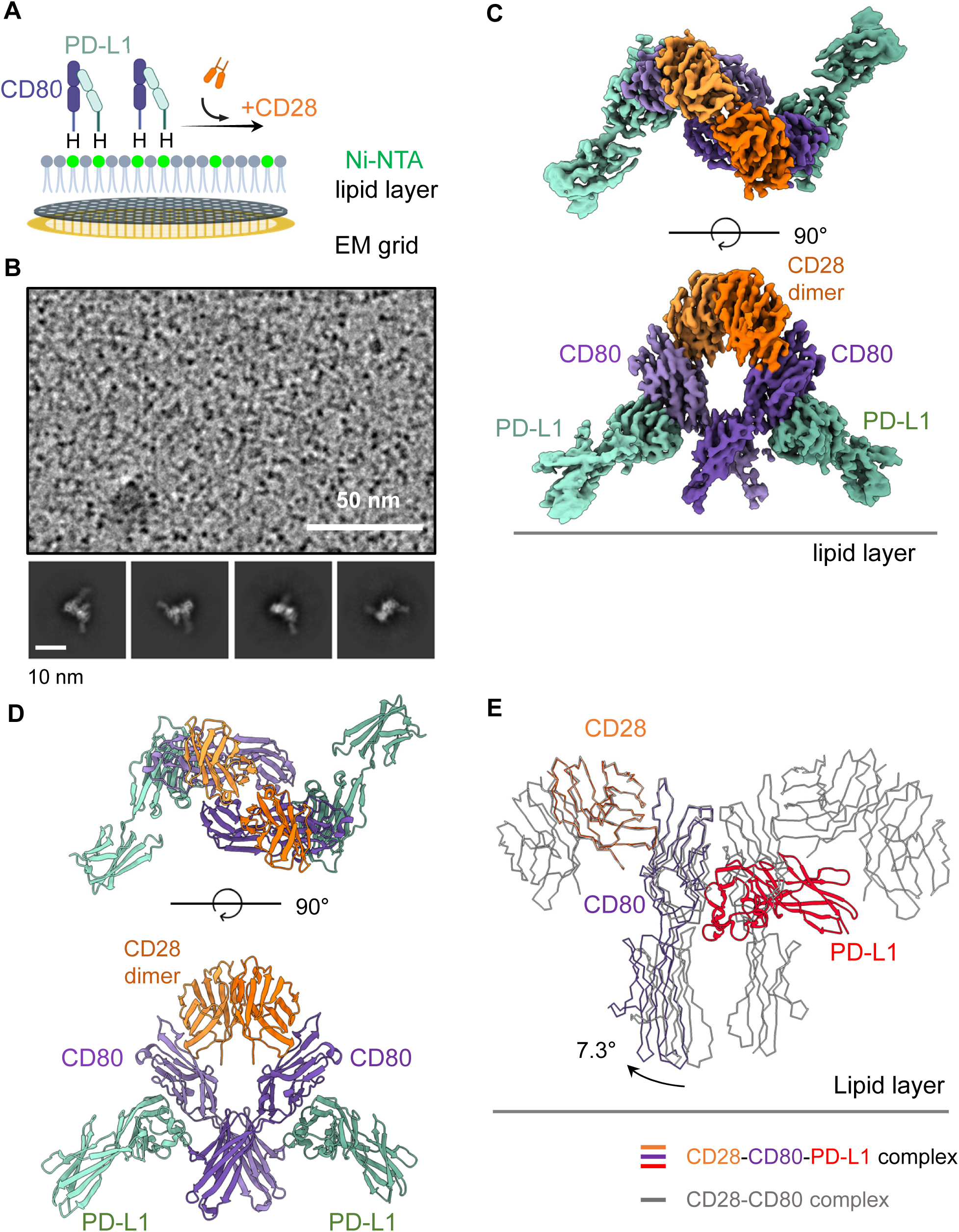
Structure of the CD28-CD80-PD-L1 complex on the lipid layer. (A) Schematic of the lipid monolayer experiment. An octa-histidine tag was fused to the C termini of the CD80 and PD-L1 ectodomains, enabling tethering of the CD28-CD80-PD-L1 complex to a Ni^2+^-NTA lipid monolayer. “H” denotes the octa-histidine tag, and Ni^2+^-NTA lipids are shown in green. (B) Representative cryo-EM micrograph and 2D class averages of the CD28-CD80-PD-L1 complex. (C) Cryo-EM density map of the CD28-CD80-PD-L1 complex. CD28, CD80, and PD-L1 are shown in light and dark shades of orange, purple, and green, respectively. (D) Overall structure of the CD28-CD80-PD-L1 complex. (E) Modeling of the CD28-CD80-PD-L1 complex on the CD28-CD80 structure (Figure 3D). The CD28 monomer structures in the two complexes are superimposed.

## DISCUSSION

In the present study, we used a lipid monolayer-based cryo-EM approach to determine the structures of CD28 and CTLA-4 bound to the B7 family ligands CD80, CD86, and PD-L1 under two-dimensional membrane confinement. Our structures reveal that, despite sharing overlapping ligand-binding interfaces, CD28 and CTLA-4 organize fundamentally distinct receptor-ligand architectures on the membrane (Figure 9). We further show that PD-L1 binding to CD80 selectively disrupts CTLA-4-mediated clustering by preventing CD80 homodimerization, while preserving CD28 engagement through a reorganization of the complex rather than through changes in the binding interface. Together, these findings provide a structural framework for understanding how membrane confinement and ligand competition govern the organization and signaling of immune checkpoint receptors at the immunological synapse.

**Figure 9.**
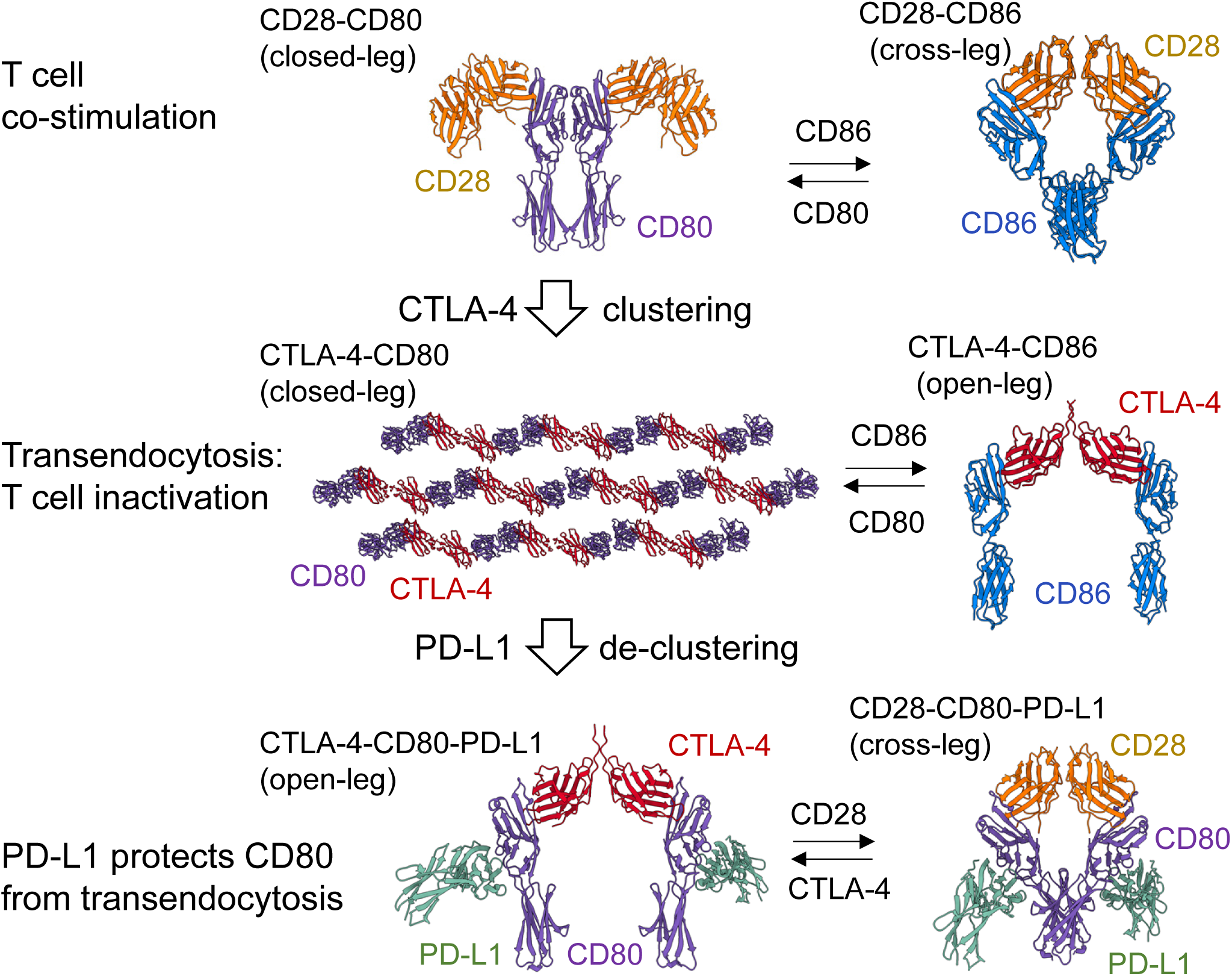
Structural basis of immune regulation by CD28, CTLA-4, and their ligands. CD28 (orange), CTLA-4 (red), CD80 (purple), CD86 (blue), and PD-L1 (green) are illustrated above. Structural data indicate that CD28 forms a discrete complex with CD80, whereas CTLA-4 displaces CD28 from this complex to promote the formation of higher-order linear and two-dimensional CD80 clusters. PD-L1 binding to CD80 disrupts CD80 homodimerization and remodels the CTLA-4–CD80 assembly into a distinct, “open-leg” CTLA-4–CD80–PD-L1 complex. These observations suggest a model in which CTLA-4-mediated clustering facilitates inhibitory receptor organization and CD80 capture, while PD-L1-mediated de-clustering attenuates CTLA-4–CD80 clustering and alters downstream T-cell signaling. Notably, the PD-L1–CD80 heterodimer preserves the ability of CD80 to bind to CD28.

To mimic a membrane-bound environment, we employed the lipid monolayer method. We recently showed that four TNF family receptor-ligand complexes, TNFα-TNFR1, BAFF-BAFFR, BAFF-BCMA, and BAFF-TACI, form highly ordered clusters on a lipid monolayer^25^. Disruption of these clusters by mutations or nanobody binding inhibits receptor activation, demonstrating that the protein clusters observed on the lipid monolayer have physiological significance. Restricting protein complexes to a lipid monolayer can have multiple effects on protein multimerization. First, it facilitates the formation of protein clusters by enhancing lateral interactions among proteins tethered to the lipid layer ^44,45^. Due to reduced dimensionality, the clustering of proteins already confined to a two-dimensional surface incurs a substantially lower entropic cost. As a result, even weak interactions can have a significant impact on protein clustering on the lipid layer. For example, the interactions that stabilize CTLA-4 and CD80 in a two-dimensional lattice appear to be very weak, and such clustering cannot be reproduced in solution, where free three-dimensional diffusion is allowed. Second, tethering receptors to a flat lipid layer prevents the formation of physiologically irrelevant protein structures ^25^. Some receptor-ligand complexes can form three-dimensional assemblies in solution that are clearly incompatible with proteins tethered to a two-dimensional biological membrane surface ^46,47^. Because of these beneficial effects, the lipid monolayer method enables the structural stabilization of B7 and CD28 family receptor-ligand complexes, as reported here.

Furthermore, using cryo-EM, we visualized the CTLA-4fl-CD80 complex on membrane fragments isolated from transfected cells. Remarkably, the clusters were clearly resolved despite the low molecular weight of the individual components: full-length CTLA-4 is approximately 20 kDa, and the CD80 ectodomain is ∼26 kDa. These findings demonstrate that cryo-EM can be successfully applied to membrane receptor assemblies previously considered too small for structural analysis, even within highly heterogeneous environments such as cellular membrane fragments. Thus, tethering membrane fragments to a lipid monolayer provides a simple and efficient strategy for determining high-resolution structures of clustered receptors in a native-like membrane environment. This approach is likely to be broadly applicable to diverse membrane protein assemblies, including ion channels, transporters, and GPCRs.

The co-inhibitory role of CTLA-4 largely depends on its unique ability to mediate transendocytosis. In this contact-dependent mechanism, CTLA-4 expressed on Tregs and activated conventional T cells extracts its ligands, CD80 and CD86, from the surface of antigen-presenting cells (APCs), physically depleting the available ligand pool ^8–10,20,33^. This depletion of co-stimulatory ligands subsequently limits access for CD28 on neighboring T cells, thereby raising the threshold for productive T-cell priming, a process critical for maintaining peripheral tolerance. Recent findings have highlighted a novel regulatory interaction involving PD-L1 that specifically intersects with the CTLA-4/CD28 pathways at the level of the CD80 ligand. PD-L1 was found to physically interact with CD80 on APCs, and this interaction blocks CTLA-4-mediated transendocytosis of CD80, suggesting that PD-L1 can protect CD80 from CTLA-4-driven depletion. Furthermore, PD-L1 can also bind to the CD28-CD80 complex; notably, however, this interaction does not interfere with CD28-mediated co-stimulatory signaling, suggesting a non-inhibitory, possibly stabilizing or protective, role for PD-L1 within the CD28-CD80 signaling synapse. Our structural observations support these immunological and biochemical studies by demonstrating that PD-L1 blocks both linear and two-dimensional clustering of the CTLA-4-CD80 complex through binding to CD80. Disruption of these clusters should reduce the avidity of CTLA-4 for CD80, as monomeric CD80 exhibits substantially lower affinity for CTLA-4 ^10^. In contrast, PD-L1 does not perturb the binding interface between CD80 and CD28, although the binding stoichiometry and overall architecture of the complex are markedly altered.

Our findings, together with previous immunological studies, underscore the importance of higher-order spatial organization as a regulatory layer in immune checkpoint signaling. The ability of CTLA-4, but not CD28, to drive extended linear and two-dimensional CD80 assemblies provides a structural explanation for its dominant inhibitory function, despite these receptors sharing overlapping binding sites. In a membrane-confined environment, repeated engagement of CD80 homodimers by CTLA-4 homodimers creates multivalent interactions that dramatically enhance overall avidity, favoring ligand sequestration and facilitating transendocytosis. This mode of regulation is fundamentally distinct from simple affinity-based competition; instead, it relies on receptor clustering that arises under two-dimensional confinement. Such a mechanism offers a unifying framework for understanding how relatively modest differences in binding interfaces and dimer geometry translate into profound functional divergence between co-stimulatory and co-inhibitory receptors at the immunological synapse.

Finally, our study has broad implications for the design and interpretation of immunotherapeutic strategies targeting the CD28-CTLA-4-PD-L1 pathway. Antibodies or biologics that disrupt CTLA-4-CD80 clustering, rather than merely blocking monomeric binding interfaces, have proven especially effective in modulating immune suppression ^25,43,48,49^. Conversely, interventions aimed at stabilizing PD-L1-CD80 interactions could selectively attenuate CTLA-4-mediated inhibition while preserving CD28 co-stimulation, consistent with recent functional studies. More broadly, the lipid monolayer-based cryo-EM approach employed here provides a highly useful platform for visualizing membrane receptor organization confined to a two-dimensional lipid layer. Applying this strategy to other immune checkpoint systems and costimulatory pathways should reveal additional layers of structural regulation that are invisible in solution-based structural analyses, ultimately guiding the rational development of next-generation immunomodulatory therapeutics.

## SUPPLEMENTAL INFORMATION

Document S1. Tables S1-S4, Figures S1-S16. Methods S1

## ACKNOWLEDGMENTS

This work was funded by the National Research Foundation of Korea (RS-2023-00260454 and RS-2024-00344154), the National Research Facilities & Equipment Center of Korea (RS-2024-00436298).

## AUTHOR CONTRIBUTIONS

J-O.L. conceived of and supervised the project. G.M.H. and C.S.L. expressed and purified the protein, prepared the sample for the cryo-EM study, collected and processed cryo-EM data, and built and refined the atomic models. G.M.H., C.S.L. and J-O.L. prepared and edited the manuscript.

## STAR METHODS

### KEY RESOURCES TABLE

| REAGENT or RESOURCE | SOURCE | IDENTIFIER |
| --- | --- | --- |
| Bacterial and virus strains |  |  |
| *E. coli* DH5α | Invitrogen | Cat# 18265017 |
| *E. coli* DH10bac | Thermo Fisher Scientific | Cat# EC0113 |
| Chemicals, peptides, and recombinant proteins |  |  |
| BestBac 2.0, Linearized Baculovirus DNA | Expression Systems | Cat# 91-002 |
| Cellfectin II Reagent | Thermo Fisher Scientific | Cat# 10362100 |
| Sodium butyrate | Sigma-Aldrich | Cat# B5887-250MG |
| Tris-HCl | LPS solution | Cat# TRI05 |
| NaCl | LPS solution | Cat# NACL05 |
| Imidazole | LPS solution | Cat# IMID500 |
| Thrombin | Lee Biosolutions | Cat# 528-50H |
| 18:1 DGS-NTA(Ni<sup>2+</sup>) | Avanti Polar Lipids | Cat# 790404C |
| 16:0-18:1 PC (POPC) | Avanti Polar Lipids | Cat# 850457C |
| FreeStyle™ 293 Expression Medium | Thermo Fisher Scientific | Cat# 12338018 |
| Sf-900™ II SFM | Thermo Fisher Scientific | Cat# 10902088 |
| ESF 921 Insect Cell Culture Medium | Expression Systems | Cat# 96-001-01 |
| Fetal Bovine Serum (U.S origin) | ATLAS biologicals | Cat# FP-0500-A |
| Deposited data |  |  |
| CTLA-4-CD86(B7-2) complex on the lipid layer | PDB<br>(http://www.rcsb.org) | PDB ID: 22KJ |
| CTLA-4-CD80(B7-1) complex on the lipid layer(CTLA-4 focused) | PDB<br>(http://www.rcsb.org) | PDB ID: 22LR |
| CTLA-4-CD80(B7-1) complex on the lipid layer(CD80 focused) | PDB<br>(http://www.rcsb.org) | PDB ID: 22LQ |
| Linear cluster of CTLA-4-CD80(B7-1) complex on the lipid layer | PDB<br>(http://www.rcsb.org) | PDB ID: 22LT |
| Two-dimensional cluster of CTLA-4-CD80(B7-1) complex on the lipid layer | PDB<br>(http://www.rcsb.org) | PDB ID: 22LU |
| CTLA-4-CD80-PD-L1 complex on the lipid layer | PDB<br>(http://www.rcsb.org) | PDB ID: 22LS |
| CD28-CD86(B7-2) complex on the lipid layer | PDB<br>(http://www.rcsb.org) | PDB ID: 22KL |
| CD28-CD80(B7-1) complex on the lipid layer | PDB<br>(http://www.rcsb.org) | PDB ID: 22KI |
| CD28-CD80-PD-L1 complex on the lipid layer | PDB<br>(http://www.rcsb.org) | PDB ID: 22KY |
| Two-dimensional cluster of CTLA-4-CD80(B7-1) complex on a cellular membrane fragment | PDB<br>(http://www.rcsb.org) | EMDB ID:<br>EMD-69318 |
| Experimental models: Cell lines |  |  |
| Sf9 insect cell | Thermo Fisher Scientific | Cat# 11496015 |
| High Five insect cell | Thermo Fisher Scientific | Cat# B85502 |
| HEK293 GnTI<sup>-</sup> cell | ATCC | Cat# CRL-3022 |
| Recombinant DNA |  |  |
| pVL1393 transfer vector | Thermo Fisher Scientific | Cat# V1392-20 |
| pAcGP67a transfer vector | Thermo Fisher Scientific | 554756 |
| pEG BacMam transfer vector | Goehring *et al.* | DOI: 10.1038/nprot.2014.173 |
| Human CD80/pVL1393 | This paper | N/A |
| Human CD86/pVL1393 | This paper | N/A |
| Human CTLA-4/pVL1393 | This paper | N/A |
| Human CD28ecto-Fc/pAcGP67A | This paper | N/A |
| Human PD-L1/pVL1393 | This paper | N/A |
| Human CTLA-4fl/pEG BacMam | This paper | N/A |
| Software and algorithms |  |  |
| EPU software for Single Particle Analysis | Thermo Fisher Scientific | https://www.thermofisher.com/ |
| Cryosparc v4.7.1 | Punjani *et al.* | https://cryosparc.com/ |
| Topaz | Bepler *et al.* | https://cb.csail.mit.edu/cb/topaz/ |
| DeepEMhancer | Sanchez-Garcia *et al.* | https://github.com/rsanchezgarc/deepEMhancer |
| PHENIX v1.21 | Adams *et al.* | http://www.phenix-online.org/ |
| MolProbity | Williams *et al.* | http://molprobity.biochem.duke.edu/ |
| Coot v0.9.8 | Emsley *et al.* | https://www2.mrc-lmb.cam.ac.uk/personal/pemsley/coot/ |
| USCF chimeraX v1.11 | Pettersen *et al.* | https://www.cgl.ucsf.edu/chimerax/ |
| Other |  |  |
| cOmplete His-Tag Purification Resin | Roche | Cat# 05893682001 |
| Protein A agarose resin | Goldbio | Cat# P-400-5 |
| Superdex 200 Increase 10/300 GL | Cytiva | Cat# 28990944 |
| Quantifoil R1.2/1.3 300-mesh Gold grids | SPI Supplies | Cat# 4230G-CF |
| Teflon chamber for lipid monolayer preparation | Lim *et al.* | DOI: 10.1038/s41467-025-61271-6 |
| Micro Syringe Pipette | Hamilton | Cat# 84250 |
| AMICON MWCO 10 kDa | Millipore | Cat# UFC901008 |
| AMICON MWCO 30 kDa | Millipore | Cat# UFC903008 |

### RESOURCE AVAILABILITY

#### Lead contact

Further information and requests for resources and reagents will be directed and fulfilled by Jie-Oh Lee, the lead contact.

#### Materials availability

Plasmids generated in this study are freely available on request.

#### Data and code availability

The cryo-EM maps and the atomic coordinates were deposited in the Electron Microscopy Data Bank and the Protein Data Bank. Their code numbers are summarized in Supplementary Tables S2-S4. This paper does not report the original code. Any additional information required to reanalyze the data reported in this paper is available from the lead contact upon request.

### EXPERIMENTAL MODEL AND SUBJECT DETAILS

DH10Bac (Thermo Fisher Scientific, EC0113) was cultured in lysogeny broth (LB) at 37℃ to amplify plasmids.

HEK293 GnTI^−^ cells (ATCC, CRL-3022) were cultured in FreeStyle 293 Medium (Thermo Fisher Scientific, 12338026)

Sf9 cells (Thermo Fisher Scientific, 11496015) were cultured in Sf-900™ II SFM (Thermo Fisher Scientific, 10902088) at 27℃

High Five cells (Thermo Fisher Scientific, B85502) were cultured in ESF 921 Medium (Expression Systems) at 27℃

### METHOD DETAILS

#### Gene cloning and protein expression

Genes encoding the extracellular domains of CTLA-4, CD28, CD80, CD86, and PD-L1, together with the indicated affinity tags, were synthesized (Twist Bioscience) and cloned into the pAcGP67A or pVL1393 baculovirus transfer vectors (BD Biosciences) (Table S1). Recombinant baculoviruses were generated by co-transfection of Sf9 insect cells with the linearized baculovirus genome BestBac2.0 (Expression Systems). For protein expression, High Five insect cells cultured in ESF 921 medium (Expression Systems) were infected with the recombinant baculovirus at 3% (v/v) and incubated at 21°C for 72 hours. The gene encoding full-length CTLA-4 with an ALFA affinity tag was synthesized (Twist Bioscience) and cloned into the pEG BacMam transfer vector (Table S1) ^50^. The plasmid was transformed into DH10Bac *E. coli* to generate the recombinant bacmid via transposition. Recombinant baculovirus was produced by transfecting Sf9 insect cells with the bacmid DNA. For protein expression, HEK293 GnTI⁻ cells cultured in FreeStyle™ 293 Expression Medium (Thermo Fisher Scientific) were infected with the recombinant baculovirus at 8% (v/v). After 12∼16 hours of infection, 10 mM of sodium butyrate was added to enhance protein expression, and the cells were subsequently incubated at 30°C for an additional 48 hours.

#### Protein purification

The extracellular domains of CTLA-4, CD80, CD86, and PD-L1 were expressed as secreted proteins with octa-histidine tags and purified using cOmplete His-Tag Purification Resin (Roche). Bound proteins were eluted with buffer containing 50 mM Tris-HCl, 200 mM NaCl, and 300 mM imidazole, pH 8.0. For CTLA-4, the histidine tag was removed by thrombin digestion at 4°C for 16 hours, followed by passage over the resin to remove uncleaved protein. All proteins were concentrated using a 10 kDa MWCO centrifugal filter and further purified by size-exclusion chromatography on a Superdex 200 Increase 10/300 GL column (Cytiva) equilibrated in 50 mM Tris-HCl and 200 mM NaCl, pH 8.0. Peak fractions were pooled and concentrated to 3 mg/ml using a 10 kDa MWCO centrifugal filter. The CD28 extracellular domain was expressed as a fusion with the human IgG Fc domain and captured on Protein A agarose resin. On-column thrombin cleavage was performed at 4°C for 16 hours in buffer containing 50 mM Tris-HCl and 200 mM NaCl, pH 8.0. Cleaved CD28 was eluted, concentrated, and further purified by size-exclusion chromatography on a Superdex 200 Increase 10/300 GL column equilibrated with the same buffer. Fractions corresponding to the CD28 homodimer were pooled and concentrated to 4 mg/ml.

For preparation of the CD28-CD86 complex, 30 µM of histidine-tag-free CD28 homodimer was mixed with 50 µM of CD86 and incubated at 37°C for 4 hours. For preparation of the CD80-PD-L1 heterodimer complex, 30 µM of histidine-tagged CD80 was mixed with 40 µM of histidine-tagged PD-L1 and incubated at 37°C for 1 hour. Each mixture was subjected to size-exclusion chromatography on a Superdex 200 Increase 10/300 GL column equilibrated in 50 mM Tris-HCl and 200 mM NaCl, pH 8.0, to remove excess unbound proteins. Fractions containing the complexes were pooled and concentrated to 1 mg/ml using a 30 kDa MWCO centrifugal filter for further analysis.

#### Preparation of cellular membrane fragments

HEK293 GnTI⁻ cells expressing full-length human CTLA-4 were resuspended in a hypotonic buffer containing 1 mM Na₂CO₃, 1 mM MgCl₂, and 1 mM CaCl₂, and lysed by passing the suspension through a fine-gauge syringe needle 10 times. To remove cell debris and nuclei, the lysate was clarified by centrifugation twice at 1,000 x g for 10 min, followed by centrifugation twice at 2,000 x g for 10 min; the supernatant was collected after each step. The resulting supernatant was then incubated with the His-tagged CD80 extracellular domain at a final concentration of 1 μM for 1 hour at 30°C. Membrane fragments were subsequently pelleted by centrifugation at 12,000 x g for 1 hour. The pellet was resuspended in buffer containing 50 mM Tris-HCl and 200 mM NaCl, pH 8.0, and subjected to gentle sonication at 70% power for 6 cycles with 5 seconds on and 40 seconds off. The prepared membrane fragments were then applied to a lipid monolayer for further characterization.

#### Preparation of lipid monolayers and cryo-EM grids

Lipid monolayers were prepared as previously described ^25^. Briefly, a Teflon block containing an 80 µl well with a side injection tunnel was used. The wells were washed sequentially with ethanol and distilled water, and the block was placed in a humidified Petri dish containing wet filter paper. Each well was filled with 80 µl of buffer consisting of 50 mM Tris-HCl and 200 mM NaCl, pH 8.0. A lipid solution, 1 µl, in chloroform containing 0.9 mg/ml POPC (Sigma) and 0.1 mg/ml DGS-NTA(Ni²⁺) (Sigma) was then gently added to the surface of the buffer. The Teflon block was left undisturbed at room temperature for 30 min to allow chloroform evaporation and spontaneous formation of a lipid monolayer.

Cryo-EM grids for the CD28-CD80 complex were prepared by removing 20 µl of buffer from the side tunnel of a Teflon well, followed by injection of 10 µl of 33 µM CD28 without a histidine tag and 10 µl of 16.7 µM histidine-tagged CD80. The solution was gently mixed by pipetting through the side tunnel and incubated at room temperature for 2 hours to allow binding to the lipid monolayer and complex formation. A Quantifoil 1.2/1.3 gold grid with carbon side down was then placed on the well for 30 min to transfer the lipid layer. After addition of 15 µl buffer to the side tunnel, the grid was retrieved, supplemented with 2 µl of 50 mM Tris-HCl and 200 mM NaCl, pH 8.0, and plunge-frozen using a Vitrobot Mark IV (Thermo Fisher Scientific) at 4°C and 100% humidity with 2.5 seconds blotting and no applied force.

Cryo-EM grids for the CD28-CD86 complex were prepared similarly, except that 10 µl of buffer was removed from the well and 10 µl of the preformed, histidine-tagged CD28-CD86 complex was injected into the well before incubating for 2 hours at room temperature. For the CTLA-4-CD80 and CTLA-4-CD86 complexes, 20 µl of buffer was removed, followed by the injection of 10 µl of 33 µM untagged CTLA-4 and 10 µl of 16.7 µM histidine-tagged CD80 or CD86. After gentle mixing and a 2-hour incubation at room temperature, grids were prepared and vitrified as described above. For the cellular membrane fragments containing the full-length CTLA-4 and CD80 extracellular domain complex, 30 µl of buffer was removed from the well. Then, 30 µl of the membrane fragment solution was injected into the well and incubated for 1 hour. Grids were then transferred, blotted, and plunge-frozen under the conditions described above.

For the CTLA-4-CD80-PD-L1 complex, 22 µl of buffer was removed, and 3 µl of 16.7 µM CD80 and 9 µl of 16.7 µM histidine-tagged PD-L1 were injected and incubated for 2 hours to allow heterodimer formation and lipid binding. A grid was placed on the well for 30 min, after which 10 µl of 33 µM CTLA-4 without a histidine tag was added and incubated for an additional 10 minutes. Grids were then processed and vitrified as described above. For the CD28-CD80-PD-L1 complex, 20 µl of buffer was removed and 10 µl of 16.7 µM histidine-tagged CD80-PD-L1 heterodimer was injected and incubated for 2 hours. Subsequently, 10 µl of 33 µM CD28 without a histidine tag was added and incubated for 1 hour. Grids were then transferred, blotted, and plunge-frozen under the same conditions as described above.

#### Cryo-EM data collection and processing

Data collection statistics are summarized in Tables S2-S4. All cryo-EM data processing was performed using CryoSPARC v4.7.1 ^51^ and is summarized in Methods S1. Briefly, cryo-EM movies were preprocessed using patch motion correction and patch-based contrast transfer function (CTF) estimation. Low-quality micrographs were manually excluded based on CTF fit, ice thickness, and total frame motion. Initial particle picking was performed using blob picking, followed by particle cleaning through 2D classification. High-quality 2D class averages were used to train a Topaz particle-picking model ^52^. Particle picking was further optimized through 2-3 rounds of Topaz training and iterative 2D classification. Initial models were generated either by *ab initio* reconstruction or by fitting known protein structures into the maps. Electron density maps were refined using combinations of 3D refinement procedures. Final maps were obtained by unbinning particles followed by 3D refinement and were sharpened and denoised using DeepEMhancer ^53^.

#### Model building

Structure templates used for initial model building were summarized in Tables S2-S4. Initial models were generated by rigid-body fitting of these reference atomic coordinates into the corresponding cryo-EM density maps using ChimeraX ^54^. These initial atomic models were subsequently refined through iterative rounds of manual model building in Coot and real-space refinement in Phenix ^55^.

## REFERENCES

1. Chambers, C.A., Kuhns, M.S., Egen, J.G., and Allison, J.P. (2001). CTLA-4-mediated inhibition in regulation of T cell responses: mechanisms and manipulation in tumor immunotherapy. Annu. Rev. Immunol. 19, 565–594. 10.1146/annurev.immunol.19.1.565.

2. Lenschow, D.J., Walunas, T.L., and Bluestone, J.A. (1996). CD28/B7 system of T cell costimulation. Annu. Rev. Immunol. 14, 233–258. 10.1146/annurev.immunol.14.1.233.

3. Sharpe, A.H., and Freeman, G.J. (2002). The B7-CD28 superfamily. Nat. Rev. Immunol. 2, 116–126. 10.1038/nri727.

4. Greenwald, R.J., Freeman, G.J., and Sharpe, A.H. (2005). The B7 family revisited. Annu. Rev. Immunol. 23, 515–548. 10.1146/annurev.immunol.23.021704.115611.

5. Tivol, E.A., Borriello, F., Schweitzer, A.N., Lynch, W.P., Bluestone, J.A., and Sharpe, A.H. (1995). Loss of CTLA-4 leads to massive lymphoproliferation and fatal multiorgan tissue destruction, revealing a critical negative regulatory role of CTLA-4. Immunity 3, 541–547. 10.1016/1074-7613(95)90125-6.

6. Pardoll, D.M. (2012). The blockade of immune checkpoints in cancer immunotherapy. Nat. Rev. Cancer 12, 252–264. 10.1038/nrc3239.

7. Rowshanravan, B., Halliday, N., and Sansom, D.M. (2018). CTLA-4: a moving target in immunotherapy. Blood 131, 58–67. 10.1182/blood-2017-06-741033.

8. Butte, M.J., Keir, M.E., Phamduy, T.B., Sharpe, A.H., and Freeman, G.J. (2007). Programmed death-1 ligand 1 interacts specifically with the B7-1 costimulatory molecule to inhibit T cell responses. Immunity 27, 111–122. 10.1016/j.immuni.2007.05.016.

9. Sugiura, D., Maruhashi, T., Okazaki, I.M., Shimizu, K., Maeda, T.K., Takemoto, T., and Okazaki, T. (2019). Restriction of PD-1 function by cis-PD-L1/CD80 interactions is required for optimal T cell responses. Science 364, 558–566. 10.1126/science.aav7062.

10. Zhao, Y., Lee, C.K., Lin, C.H., Gassen, R.B., Xu, X., Huang, Z., Xiao, C., Bonorino, C., Lu, L.F., Bui, J.D., and Hui, E. (2019). PD-L1:CD80 Cis-Heterodimer Triggers the Co-stimulatory Receptor CD28 While Repressing the Inhibitory PD-1 and CTLA-4 Pathways. Immunity 51, 1059–1073 e1059. 10.1016/j.immuni.2019.11.003.

11. Butte, M.J., Pena-Cruz, V., Kim, M.J., Freeman, G.J., and Sharpe, A.H. (2008). Interaction of human PD-L1 and B7-1. Mol. Immunol. 45, 3567–3572. 10.1016/j.molimm.2008.05.014.

12. Evans, E.J., Esnouf, R.M., Manso-Sancho, R., Gilbert, R.J., James, J.R., Yu, C., Fennelly, J.A., Vowles, C., Hanke, T., Walse, B., et al. (2005). Crystal structure of a soluble CD28-Fab complex. Nat. Immunol. 6, 271–279. 10.1038/ni1170.

13. Stamper, C.C., Zhang, Y., Tobin, J.F., Erbe, D.V., Ikemizu, S., Davis, S.J., Stahl, M.L., Seehra, J., Somers, W.S., and Mosyak, L. (2001). Crystal structure of the B7-1/CTLA-4 complex that inhibits human immune responses. Nature 410, 608–611. 10.1038/35069118.

14. Schwartz, J.C., Zhang, X., Fedorov, A.A., Nathenson, S.G., and Almo, S.C. (2001). Structural basis for co-stimulation by the human CTLA-4/B7-2 complex. Nature 410, 604–608. 10.1038/35069112.

15. Ikemizu, S., Gilbert, R.J., Fennelly, J.A., Collins, A.V., Harlos, K., Jones, E.Y., Stuart, D.I., and Davis, S.J. (2000). Structure and dimerization of a soluble form of B7-1. Immunity 12, 51–60. 10.1016/s1074-7613(00)80158-2.

16. Lin, D.Y., Tanaka, Y., Iwasaki, M., Gittis, A.G., Su, H.P., Mikami, B., Okazaki, T., Honjo, T., Minato, N., and Garboczi, D.N. (2008). The PD-1/PD-L1 complex resembles the antigen-binding Fv domains of antibodies and T cell receptors. Proc. Natl. Acad. Sci. U S A 105, 3011–3016. 10.1073/pnas.0712278105.

17. van der Merwe, P.A., and Davis, S.J. (2003). Molecular interactions mediating T cell antigen recognition. Annu. Rev. Immunol. 21, 659–684. 10.1146/annurev.immunol.21.120601.141036.

18. Bhatia, S., Edidin, M., Almo, S.C., and Nathenson, S.G. (2005). Different cell surface oligomeric states of B7-1 and B7-2: implications for signaling. Proc. Natl. Acad. Sci. U S A 102, 15569–15574. 10.1073/pnas.0507257102.

19. Dustin, M.L. (2014). The immunological synapse. Cancer Immunol. Res. 2, 1023–1033. 10.1158/2326-6066.CIR-14-0161.

20. Qureshi, O.S., Zheng, Y., Nakamura, K., Attridge, K., Manzotti, C., Schmidt, E.M., Baker, J., Jeffery, L.E., Kaur, S., Briggs, Z., et al. (2011). Trans-endocytosis of CD80 and CD86: a molecular basis for the cell-extrinsic function of CTLA-4. Science 332, 600–603. 10.1126/science.1202947.

21. Xiao, Q., McAtee, C.K., and Su, X. (2022). Phase separation in immune signalling. Nat. Rev. Immunol. 22, 188–199. 10.1038/s41577-021-00572-5.

22. Su, X., Ditlev, J.A., Rosen, M.K., and Vale, R.D. (2017). Reconstitution of TCR Signaling Using Supported Lipid Bilayers. Methods Mol. Biol. 1584, 65–76. 10.1007/978-1-4939-6881-7_5.

23. Dustin, M.L., and Groves, J.T. (2012). Receptor signaling clusters in the immune synapse. Annu. Rev. Biophys. 41, 543–556. 10.1146/annurev-biophys-042910-155238.

24. Yokosuka, T., Kobayashi, W., Sakata-Sogawa, K., Takamatsu, M., Hashimoto-Tane, A., Dustin, M.L., Tokunaga, M., and Saito, T. (2008). Spatiotemporal regulation of T cell costimulation by TCR-CD28 microclusters and protein kinase C theta translocation. Immunity 29, 589–601. 10.1016/j.immuni.2008.08.011.

25. Lim, C.S., Lee, J., Kim, J.W., and Lee, J.O. (2025). Highly ordered clustering of TNFalpha and BAFF ligand-receptor-intracellular adaptor complexes on a lipid membrane. Nat. Commun. 16, 5551. 10.1038/s41467-025-61271-6.

26. Skrajna, A., Lenger, C., Robinson, E., Cannon, K., Sarsam, R., Ouellette, R.G., Abotsi, A.M., Brennwald, P., McGinty, R.K., Strauss, J.D., and Baker, R.W. (2025). Nickel-NTA lipid-monolayer affinity grids allow for high-resolution structure determination by cryo-EM. J. Struct. Biol. 217, 108253. 10.1016/j.jsb.2025.108253.

27. Kelly, D.F., Dukovski, D., and Walz, T. (2010). A practical guide to the use of monolayer purification and affinity grids. Methods Enzymol. 481, 83–107. 10.1016/S0076-6879(10)81004-3.

28. Truong, C.D., Williams, D.R., Zhu, M., Wang, J.C., and Chiu, P.L. (2021). Sample Preparation using a Lipid Monolayer Method for Electron Crystallographic Studies. J. Vis. Exp. 10.3791/63015.

29. Pentcheva-Hoang, T., Egen, J.G., Wojnoonski, K., and Allison, J.P. (2004). B7-1 and B7-2 selectively recruit CTLA-4 and CD28 to the immunological synapse. Immunity 21, 401–413. 10.1016/j.immuni.2004.06.017.

30. Peach, R.J., Bajorath, J., Naemura, J., Leytze, G., Greene, J., Aruffo, A., and Linsley, P.S. (1995). Both extracellular immunoglobin-like domains of CD80 contain residues critical for binding T cell surface receptors CTLA-4 and CD28. J. Biol. Chem. 270, 21181–21187. 10.1074/jbc.270.36.21181.

31. Rohr, J., Guo, S., Huo, J., Bouska, A., Lachel, C., Li, Y., Simone, P.D., Zhang, W., Gong, Q., Wang, C., et al. (2016). Recurrent activating mutations of CD28 in peripheral T-cell lymphomas. Leukemia 30, 1062–1070. 10.1038/leu.2015.357.

32. Abramson, J., Adler, J., Dunger, J., Evans, R., Green, T., Pritzel, A., Ronneberger, O., Willmore, L., Ballard, A.J., Bambrick, J., et al. (2024). Accurate structure prediction of biomolecular interactions with AlphaFold 3. Nature 630, 493–500. 10.1038/s41586-024-07487-w.

33. Kennedy, A., Waters, E., Rowshanravan, B., Hinze, C., Williams, C., Janman, D., Fox, T.A., Booth, C., Pesenacker, A.M., Halliday, N., et al. (2022). Differences in CD80 and CD86 transendocytosis reveal CD86 as a key target for CTLA-4 immune regulation. Nat. Immunol. 23, 1365–1378. 10.1038/s41590-022-01289-w.

34. Harris, N., Peach, R., Naemura, J., Linsley, P.S., Le Gros, G., and Ronchese, F. (1997). CD80 costimulation is essential for the induction of airway eosinophilia. J. Exp. Med. 185, 177–182. 10.1084/jem.185.1.177.

35. Reynolds, J., Tam, F.W., Chandraker, A., Smith, J., Karkar, A.M., Cross, J., Peach, R., Sayegh, M.H., and Pusey, C.D. (2000). CD28-B7 blockade prevents the development of experimental autoimmune glomerulonephritis. J. Clin. Invest. 105, 643–651. 10.1172/JCI6710.

36. Morton, P.A., Fu, X.T., Stewart, J.A., Giacoletto, K.S., White, S.L., Leysath, C.E., Evans, R.J., Shieh, J.J., and Karr, R.W. (1996). Differential effects of CTLA-4 substitutions on the binding of human CD80 (B7-1) and CD86 (B7-2). J. Immunol. 156, 1047–1054.

37. Peach, R.J., Bajorath, J., Brady, W., Leytze, G., Greene, J., Naemura, J., and Linsley, P.S. (1994). Complementarity determining region 1 (CDR1)- and CDR3-analogous regions in CTLA-4 and CD28 determine the binding to B7-1. J. Exp. Med. 180, 2049–2058. 10.1084/jem.180.6.2049.

38. Kuehn, H.S., Ouyang, W., Lo, B., Deenick, E.K., Niemela, J.E., Avery, D.T., Schickel, J.N., Tran, D.Q., Stoddard, J., Zhang, Y., et al. (2014). Immune dysregulation in human subjects with heterozygous germline mutations in CTLA4. Science 345, 1623–1627. 10.1126/science.1255904.

39. Mitsuiki, N., Schwab, C., and Grimbacher, B. (2019). What did we learn from CTLA-4 insufficiency on the human immune system? Immunol. Rev. 287, 33–49. 10.1111/imr.12721.

40. Schubert, D., Bode, C., Kenefeck, R., Hou, T.Z., Wing, J.B., Kennedy, A., Bulashevska, A., Petersen, B.S., Schaffer, A.A., Gruning, B.A., et al. (2014). Autosomal dominant immune dysregulation syndrome in humans with CTLA4 mutations. Nat. Med. 20, 1410–1416. 10.1038/nm.3746.

41. Schwab, C., Gabrysch, A., Olbrich, P., Patino, V., Warnatz, K., Wolff, D., Hoshino, A., Kobayashi, M., Imai, K., Takagi, M., et al. (2018). Phenotype, penetrance, and treatment of 133 cytotoxic T-lymphocyte antigen 4-insufficient subjects. J. Allergy Clin. Immunol. 142, 1932–1946. 10.1016/j.jaci.2018.02.055.

42. Chaudhri, A., Xiao, Y., Klee, A.N., Wang, X., Zhu, B., and Freeman, G.J. (2018). PD-L1 Binds to B7-1 Only In Cis on the Same Cell Surface. Cancer Immunol. Res. 6, 921–929. 10.1158/2326-6066.CIR-17-0316.

43. Maurer, M.F., Lewis, K.E., Kuijper, J.L., Ardourel, D., Gudgeon, C.J., Chandrasekaran, S., Mudri, S.L., Kleist, K.N., Navas, C., Wolfson, M.F., et al. (2022). The engineered CD80 variant fusion therapeutic davoceticept combines checkpoint antagonism with conditional CD28 costimulation for anti-tumor immunity. Nat. Commun. 13, 1790. 10.1038/s41467-022-29286-5.

44. Adam, G.D., M. (1968). Reduction of dimensionality in biological diffusion processes (W. H. Freeman).

45. Grebenkov, D.M., Metzler, R., and Oshanin, G. (2022). Search efficiency in the Adam-Delbruck reduction-of-dimensionality scenario versus direct diffusive search. New J. Phys. 24, 083035.

46. Kim, H.M., Yu, K.S., Lee, M.E., Shin, D.R., Kim, Y.S., Paik, S.G., Yoo, O.J., Lee, H., and Lee, J.O. (2003). Crystal structure of the BAFF-BAFF-R complex and its implications for receptor activation. Nat. Struct. Biol. 10, 342–348. 10.1038/nsb925.

47. Liu, Y., Hong, X., Kappler, J., Jiang, L., Zhang, R., Xu, L., Pan, C.H., Martin, W.E., Murphy, R.C., Shu, H.B., et al. (2003). Ligand-receptor binding revealed by the TNF family member TALL-1. Nature 423, 49–56. 10.1038/nature01543.

48. Zhang, Y., Song, Q., Cassady, K., Lee, M., Tang, H., Zheng, M., Wang, B., Schones, D.E., Fu, Y.X., Riggs, A.D., et al. (2023). Blockade of trans PD-L1 interaction with CD80 augments antitumor immunity. Proc. Natl. Acad. Sci. U S A 120, e2205085120. 10.1073/pnas.2205085120.

49. Sugiura, D., Okazaki, I.M., Maeda, T.K., Maruhashi, T., Shimizu, K., Arakaki, R., Takemoto, T., Ishimaru, N., and Okazaki, T. (2022). PD-1 agonism by anti-CD80 inhibits T cell activation and alleviates autoimmunity. Nat. Immunol. 23, 399–410. 10.1038/s41590-021-01125-7.

50. Goehring, A., Lee, C.H., Wang, K.H., Michel, J.C., Claxton, D.P., Baconguis, I., Althoff, T., Fischer, S., Garcia, K.C., and Gouaux, E. (2014). Screening and large-scale expression of membrane proteins in mammalian cells for structural studies. Nat. Protoc. 9, 2574–2585. 10.1038/nprot.2014.173.

51. Punjani, A., Rubinstein, J.L., Fleet, D.J., and Brubaker, M.A. (2017). cryoSPARC: algorithms for rapid unsupervised cryo-EM structure determination. Nat. Methods 14, 290–296. 10.1038/nmeth.4169.

52. Bepler, T., Morin, A., Rapp, M., Brasch, J., Shapiro, L., Noble, A.J., and Berger, B. (2019). Positive-unlabeled convolutional neural networks for particle picking in cryo-electron micrographs. Nat. Methods 16, 1153–1160. 10.1038/s41592-019-0575-8.

53. Sanchez-Garcia, R., Gomez-Blanco, J., Cuervo, A., Carazo, J.M., Sorzano, C.O.S., and Vargas, J. (2021). DeepEMhancer: a deep learning solution for cryo-EM volume post-processing. Commun. Biol. 4, 874. 10.1038/s42003-021-02399-1.

54. Meng, E.C., Goddard, T.D., Pettersen, E.F., Couch, G.S., Pearson, Z.J., Morris, J.H., and Ferrin, T.E. (2023). UCSF ChimeraX: Tools for structure building and analysis. Protein Sci. 32, e4792. 10.1002/pro.4792.

55. Liebschner, D., Afonine, P.V., Baker, M.L., Bunkoczi, G., Chen, V.B., Croll, T.I., Hintze, B., Hung, L.W., Jain, S., McCoy, A.J., et al. (2019). Macromolecular structure determination using X-rays, neutrons and electrons: recent developments in Phenix. Acta Crystallogr. D Struct. Biol. 75, 861–877. 10.1107/S2059798319011471.

