## Supplemental Tables and Figures for "Structural basis of CD28 and CTLA-4 interactions with CD80, CD86, and the CD80-PD-L1 heterodimer on artificial and cellular membranes"

### Documents S1: Tables S1-S4, Figures S1-S16

**Table S1. CD28, CTLA-4, CD80, CD86 and PD-L1 protein sequences**

| Protein name | Protein sequence |
|---|---|
| CD80<br>(as encoded<br>in the plasmid) | MGHTRRQGTSPSKCPYLNFFQLLVLAGLSHFCSG<br>VIHVTKEVKEVATLSCGHNVSVEELAQTRIYWQKEKKMVLTMMSGDMNIWPEYKNR<br>TIFDITNNLSIVILALRPSDEGTYECVVLKYEKDAFKREHLAEVTLSVKADFPTPSISDF<br>EIPTSNIRRIICSTSGGFPEPHLSWLENGEELNAINTTVSQDPETELYAVSSKLDFNMT<br>TNHSFMCLIKYGHLRVNQTFNWNTTKQEHFPDN<br>GGGGSGGGGSHHHHHHHH |
| CD80<br>(after purification) | VIHVTKEVKEVATLSCGHNVSVEELAQTRIYWQKEKKMVLTMMSGDMNIWPEYKNR<br>TIFDITNNLSIVILALRPSDEGTYECVVLKYEKDAFKREHLAEVTLSVKADFPTPSISDF<br>EIPTSNIRRIICSTSGGFPEPHLSWLENGEELNAINTTVSQDPETELYAVSSKLDFNMT<br>TNHSFMCLIKYGHLRVNQTFNWNTTKQEHFPDN<br>GGGGSGGGGSHHHHHHHH |
| CD86<br>(as encoded in<br>the plasmid) | MDPQCTMGLSNILFVMAFLLSGA<br>APLKIQAYFNETADLPCQFANSQNQSLSELVVFWQDQENLVLNEVYLGKEKFDSVHS<br>KYMGRTSFDSDSWTLRLHNLQIKDKGLYQCIIHHKKPTGMIRIHQMNSELSVLANFSQ<br>PEIVPISNITENVYINLTCSSIHGYPEPKKMSVLLRTKNSTIEYDGVMQKSQDNVTELY<br>DVSISLSVSFPDVTSNMTIFCILETDKTRLLSSPFSIELEDPQPPPDHIP<br>GGGGSGGGGSHHHHHHHH |
| CD86<br>(after purification) | APLKIQAYFNETADLPCQFANSQNQSLSELVVFWQDQENLVLNEVYLGKEKFDSVHS<br>KYMGRTSFDSDSWTLRLHNLQIKDKGLYQCIIHHKKPTGMIRIHQMNSELSVLANFSQ<br>PEIVPISNITENVYINLTCSSIHGYPEPKKMSVLLRTKNSTIEYDGVMQKSQDNVTELY<br>DVSISLSVSFPDVTSNMTIFCILETDKTRLLSSPFSIELEDPQPPPDHIP<br>GGGGSGGGGSHHHHHHHH |
| CD28<br>(as encoded in<br>the plasmid) | MVSAIVLYVLLAAAAHSAFAADP<br>NKILVKQSPMLVAYDNAVNLSCKYSYNLFSREFRASLHKGLDSAVEVCVVYGNYSQ<br>QLQVYSKTGFNCDGKLGNESVTFYLQNLYVNQTDIYFCKIEVMYPPPYLDNEKSNGT<br>IIHVKGKHLCPSPLFPGPSKP<br>GGSLVPRGSGHHHHHHHHESKYGPPCPSCPAPEFLGGPSVFLFPPKPKDTLMISRT<br>PEVTCVVVDVSQEDPEVQFNWYVDGVEVHNAKTKPREEQFNSTYRVVSVLTVLHQ<br>DWLNGKEYKCKVSNKGLPSSIEKTISKAKGQPREPQVYTLPPSQEEMTKNQVSLTCL<br>VKGFYPSDIAVEWESNGQPENNYKTTPPVLDSDGSFFLYSRLTVDKSRWQEGNVFS<br>CSVMHEALHNHYTQKSLSLSLGK |
| CD28<br>(after purification) | NKILVKQSPMLVAYDNAVNLSCKYSYNLFSREFRASLHKGLDSAVEVCVVYGNYSQ<br>QLQVYSKTGFNCDGKLGNESVTFYLQNLYVNQTDIYFCKIEVMYPPPYLDNEKSNGT<br>IIHVKGKHLCPSPLFPGPSKP<br>GGSLVPR |
| CTLA-4<br>(as encoded in<br>the plasmid) | MACLGFQRHKAQLNLATRTWPCTLLFFLLFIPVFCKA<br>MHVAQPAVVLASSRGIASFVCEYASPGKATEVRVTVLRQADSQVTEVCAATYMMGN<br>ELTFLDDSICTGTSSGNQVNLTIQGLRAMDTGLYICKVELMYPPPYYLGIGNGTQIYVI<br>DPEPCPDSD<br>GGGGSLVPRGSHHHHHHHH |
| CTLA-4<br>(after purification) | MHVAQPAVVLASSRGIASFVCEYASPGKATEVRVTVLRQADSQVTEVCAATYMMGN<br>ELTFLDDSICTGTSSGNQVNLTIQGLRAMDTGLYICKVELMYPPPYYLGIGNGTQIYVI<br>DPEPCPDSD<br>GGGGSLVPR |
| CTLA-4<br>(after purification<br>without thrombin<br>cleavage) | MHVAQPAVVLASSRGIASFVCEYASPGKATEVRVTVLRQADSQVTEVCAATYMMGN<br>ELTFLDDSICTGTSSGNQVNLTIQGLRAMDTGLYICKVELMYPPPYYLGIGNGTQIYVI<br>DPEPCPDSD<br>GGGGSLVPRGSHHHHHHHH |

| | |
|---|---|
| CTLA-4fl<br>(as encoded in<br>the transfected plasmid) | MACLGFQRHKAQLNLATRTWPCTLLFFLLFIPVFCKA<br>MHVAQPAVVLASSRGIASFVCEYASPGKATEVRVTVLRQADSQVTEVCAATYMMGN<br>ELTFLDDSICTGTSSGNQVNLTIQGLRAMDTGLYICKVELMYPPPYYLGIGNGTQIYVI<br>DPEPCPDSDFLLWILAAVSSGLFFYSFLLTAVSLSKMLKKRSPLTTGVYVKMPPTEPE<br>CEKQFQPYFIPIN<br>GGSLVPRGSASAPSRLEEELRRRLTE |
| PD-L1<br>(as encoded in the plasmid) | MRIFAVFIFMTYWHLLNA<br>FTVTVPKDLYVVEYGSNMTIECKFPVEKQLDLAALIVYWEMEDKNIIQFVHGEEDLKV<br>QHSSYRQRARLLKDQLSLGNAALQITDVKLQDAGVYRCMISYGGADYKRITVKVNAP<br>YNKINQRILVVDPVTSEHELTCQAEGYPKAEVIWTSSDHQVLSGKTTTTNSKREEKLF<br>NVTSTLRINTTTNEIFYCTFRRLDPEENHTAELVIPELPLAHPPNER<br>GGGGSGGGGSHHHHHHHH |
| PD-L1<br>(after purification) | FTVTVPKDLYVVEYGSNMTIECKFPVEKQLDLAALIVYWEMEDKNIIQFVHGEEDLKV<br>QHSSYRQRARLLKDQLSLGNAALQITDVKLQDAGVYRCMISYGGADYKRITVKVNAP<br>YNKINQRILVVDPVTSEHELTCQAEGYPKAEVIWTSSDHQVLSGKTTTTNSKREEKLF<br>NVTSTLRINTTTNEIFYCTFRRLDPEENHTAELVIPELPLAHPPNER<br>GGGGSGGGGSHHHHHHHH |

Footnote: Non-native sequences, including signal peptides, flexible linkers, histidine tags, an ALFA tag, thrombin cleavage sites, and an Fc tag, are written in purple.

**Table S2. Cryo-EM data statistics of the CD28-CD86, CTLA-4-CD86 and CD28-CD80**

| Receptor complexes | CD28-CD86 | CTLA-4-CD86 | CD28-CD80 |
|---|---|---|---|
| Lipid layer | Lipid monolayer | Lipid monolayer | Lipid monolayer |
| Complex molar ratio | 2:2 | 2:2 | 2:4 |
| EMDB code | EMD-68399 | EMD-68397 | EMD-68395 |
| PDB code | 22KL | 22KJ | 22KI |
| **Data** | | | |
| Magnification | 200,000 x | 105,000 x | 130,000 x |
| Voltage (kV) | 300 | 300 | 300 |
| Electron exposure ($e^{-}/Å^2$) | 40 | 40 | 40 |
| Defocus range (μm) | -0.8 to -2.0 | -0.8 to -2.0 | -0.8 to -2.0 |
| Pixel size (Å) | 0.6 | 0.8265 | 0.9388 |
| Tilt angles (˚) | 0 | 0 | 0, 30 |
| Box size (pixels) | 360 | 360 | 360 |
| Symmetry imposed | C2 | C2 | C2 |
| Initial particle images (no.) | 231,209 | 420,286 | 436,027 |
| Final particle images (no.) | 132,749 | 261,391 | 196,933 |
| Map resolution (Å) | 3.39 | 3.08 | 3.47 |
| FSC threshold | 0.143 | 0.143 | 0.143 |
| **Refinement** | | | |
| Initial structure templates (PDB code) | 1YJD (CD28)<br>22KJ (CD86, this study) | 1I8L (CTLA-4)<br>AlphaFold3 (CD86) | 1YJD (CD28)<br>1I8L (CD80) |
| Model resolution (Å) | 4.0 | 3.23 | 3.9 |
| FSC threshold | 0.5 | 0.5 | 0.5 |
| Map sharpening B factor ($Å^2$) | 137.9 | 115.5 | 135.2 |
| Model composition | | | |
| Non-hydrogen atoms | 5,062 | 5,164 | 6,728 |
| Protein residues | 652 | 660 | 858 |
| B factors ($Å^2$) | | | |
| Protein | 98.09 | 82.86 | 83.14 |
| R.m.s. deviations | | | |
| Bond lengths (Å) | 0.002 | 0.004 | 0.004 |
| Bond angles (°) | 0.495 | 0.563 | 0.706 |
| Validation | | | |
| MolProbity score | 1.82 | 1.50 | 1.94 |
| Clashscore | 7.86 | 7.32 | 8.79 |
| Poor rotamers (%) | 1.22 | 0.83 | 1.61 |
| Ramachandran plot | | | |
| Favored (%) | 95.31 | 97.55 | 95.47 |
| Allowed (%) | 4.69 | 2.45 | 4.53 |
| Disallowed (%) | 0 | 0 | 0 |

**Table S3. Cryo-EM data statistics of the CTLA-4-CD80 clusters**

| Receptor complexes | CTLA-4-CD80 (CTLA-4 focused) | CTLA-4-CD80 (CD80 focused) | CTLA-4-CD80 (linear cluster) | CTLA-4-CD80 (2D cluster) |
|---|---|---|---|---|
| Lipid layer | Lipid monolayer | Lipid monolayer | Lipid monolayer | Lipid monolayer |
| Complex molar ratio | 2:2 | 2:2 | Linear cluster | 2D Lattice |
| EMDB code | EMD-68440 | EMD-68437 | EMD-68444 | EMD-68446 |
| PDB code | 22LR | 22LQ | 22LT | 22LU |
| **Data** | | | | |
| Magnification | 130,000 x | 130,000 x | 130,000 x | 130,000 x |
| Voltage (kV) | 300 | 300 | 300 | 300 |
| Electron exposure ($e^{-}/Å^2$) | 40 | 40 | 40 | 40 |
| Defocus range (μm) | -0.8 to -2.0 | -0.8 to -2.0 | -0.8 to -2.0 | -0.8 to -2.0 |
| Pixel size (Å) | 0.9388 | 0.9388 | 0.9388 | 0.9388 |
| Tilt angles (˚) | 0, 30 | 0, 30 | 0, 30 | 0, 30 |
| Box size (pixels) | 720 | 800 | 1,600 | 1,200 |
| Symmetry imposed | C2 | C2 | C1 | C1 |
| Initial particle images (no.) | 632,183 | 2,206,082 | 242,638 | 292,140 |
| Final particle images (no.) | 293,924 | 1,019,017 | 78,088 | 95,478 |
| Map resolution (Å) | 3.54 | 3.29 | 8.33 | 7.65 |
| FSC threshold | 0.143 | 0.143 | 0.143 | 0.143 |
| **Refinement** | | | | |
| Initial structure templates (PDB code) | 1I8L | 1I8L | 22LR (this study) | 22LR (this study) |
| Model resolution (Å) | 3.82 | 3.8 | 11.6 | 11.0 |
| FSC threshold | 0.5 | 0.5 | 0.5 | 0.5 |
| Map sharpening B factor ($Å^2$) | 74.9 | 76.7 | 604.1 | 426.7 |
| Model composition | | | | |
| Non-hydrogen atoms | 4,905 | 5,040 | 14,715 | 41,706 |
| Protein residues | 634 | 644 | 1,902 | 5,389 |
| B factors ($Å^2$) | | | | |
| Protein | 100.99 | 98.79 | 100.99 | 100.98 |
| R.m.s. deviations | | | | |
| Bond lengths (Å) | 0.003 | 0.003 | 0.003 | 0.003 |
| Bond angles (°) | 0.629 | 0.589 | 0.650 | 0.649 |
| Validation | | | | |
| MolProbity score | 1.57 | 1.62 | 1.73 | 1.74 |
| Clashscore | 8.05 | 8.71 | 8.60 | 9.18 |
| Poor rotamers (%) | 0.73 | 0.52 | 1.52 | 1.41 |
| Ramachandran plot | | | | |
| Favored (%) | 97.28 | 97.17 | 97.34 | 97.29 |
| Allowed (%) | 2.72 | 2.83 | 2.66 | 2.71 |
| Disallowed (%) | 0 | 0 | 0 | 0 |

**Table S4. Cryo-EM data statistics of CTLA-4fl-CD80 2D cluster on native membrane, CTLA-4-CD80-PD-L1 and CD28-CD80-PD-L1**

| Receptor complexes | CTLA-4fl-CD80 | CTLA-4-CD80-PD-L1 | CD28-CD80-PD-L1 |
|---|---|---|---|
| Lipid layer | Cellular membrane fragment-<br>Lipid monolayer | Lipid monolayer | Lipid monolayer |
| Complex molar ratio | 2:2 | 2:2:2 | 2:2:2 |
| EMDB code | EMD-69318 | EMD-68442 | EMD-68411 |
| PDB code | N/A | 22LS | 22KY |
| **Data** | | | |
| Magnification | 105,000 x | 105,000 x | 105,000 x |
| Voltage (kV) | 300 | 300 | 300 |
| Electron exposure ($e^{-}/Å^2$) | 35 | 40 | 50 |
| Defocus range (μm) | -1.2 to -3.0 | -0.8 to -2.0 | -0.8 to -2.0 |
| Pixel size (Å) | 1.2 | 1.1972 | 0.8248 |
| Tilt angles (˚) | 0, 20 | 0, 30 | 0, 30 |
| Box size (pixels) | 400 | 480 | 400 |
| Symmetry imposed | C2 | C2 | C2 |
| Initial particle images (no.) | 112,317 | 477,937 | 1,385,607 |
| Final particle images (no.) | 29,581 | 385,964 | 728,612 |
| Map resolution (Å) | 10.11 | 3.66 | 3.33 |
| FSC threshold | 0.143 | 0.143 | 0.143 |
| **Refinement** | | | |
| Initial structure templates (PDB code) | | 22LR (CTLA-4, CD80, this study)<br>7TPS (PD-L1) | 1YJD (CD28)<br>22LQ (CD80, this study)<br>7TPS (PD-L1) |
| Model resolution (Å) | | 4.1 | 3.9 |
| FSC threshold | | 0.5 | 0.5 |
| Map sharpening B factor ($Å^2$) | | 74.2 | 145.1 |
| Model composition | | | |
| Non-hydrogen atoms | | 8,310 | 8,088 |
| Protein residues | | 1,060 | 1,040 |
| B factors ($Å^2$) | | | |
| Protein | | 123.47 | 98.54 |
| R.m.s. deviations | | | |
| Bond lengths (Å) | | 0.004 | 0.003 |
| Bond angles (°) | | 0.666 | 0.617 |
| Validation | | | |
| MolProbity score | | 2.04 | 1.96 |
| Clashscore | | 12.03 | 8.71 |
| Poor rotamers (%) | | 1.40 | 1.58 |
| Ramachandran plot | | | |
| Favored (%) | | 95.23 | 95.10 |
| Allowed (%) | | 4.77 | 4.90 |
| Disallowed (%) | | 0 | 0 |

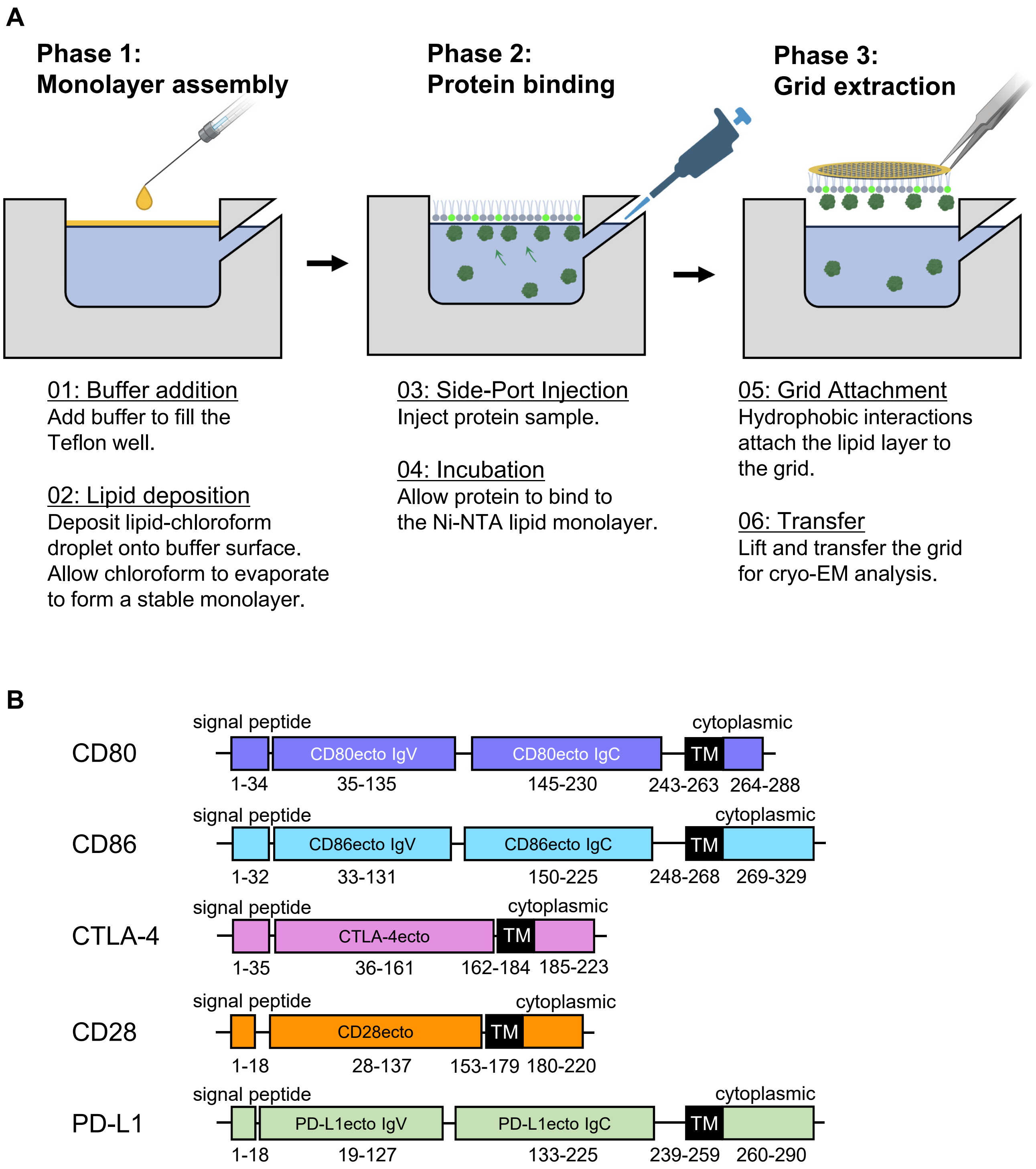

**Figure S1. Preparation of lipid monolayer grids and protein domain organization.**
(A) Grid preparation procedure.
(B) Schematic diagrams illustrating the domain organization of the proteins. Amino acid residues are numbered according to the UniProt database (https://www.uniprot.org).

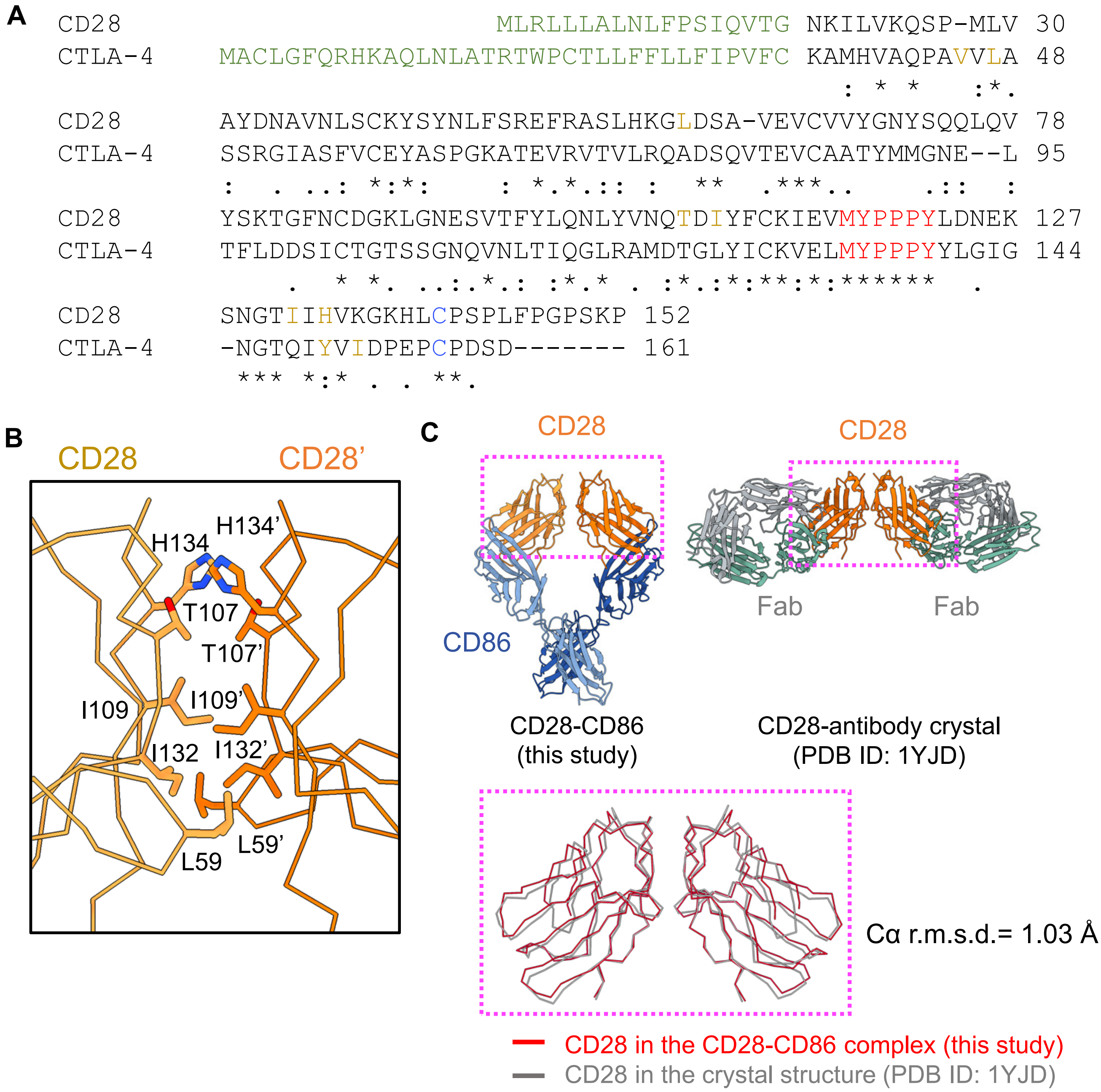

**Figure S2. Structure of the homodimerization interface of CD28.**
(A) Sequence alignment of the ectodomains of CD28 and CTLA-4. Signal peptides, MYPPPY motifs, residues involved in the homodimerization interface, and cysteines forming conserved inter-subunit disulfide bridges are colored green, red, orange, and blue, respectively. Asterisks, colons, and periods below the alignment indicate identical, highly conserved, and conserved residues, respectively.
(B) Close-up view of the CD28 homodimerization interface. The two CD28 monomeric subunits are shown in dark and light orange, respectively. Amino acid residues at the dimerization interface are labeled. Residue numbers for the second monomer in the CD28 dimer are indicated by prime symbols.
(C) Structural comparison of CD28 dimers. CD28 dimers from the cryo-EM structure of the CD28-CD86 complex (upper left) and the previously reported CD28-antibody complex (upper right) are highlighted with magenta boxes and superimposed in the lower panel.

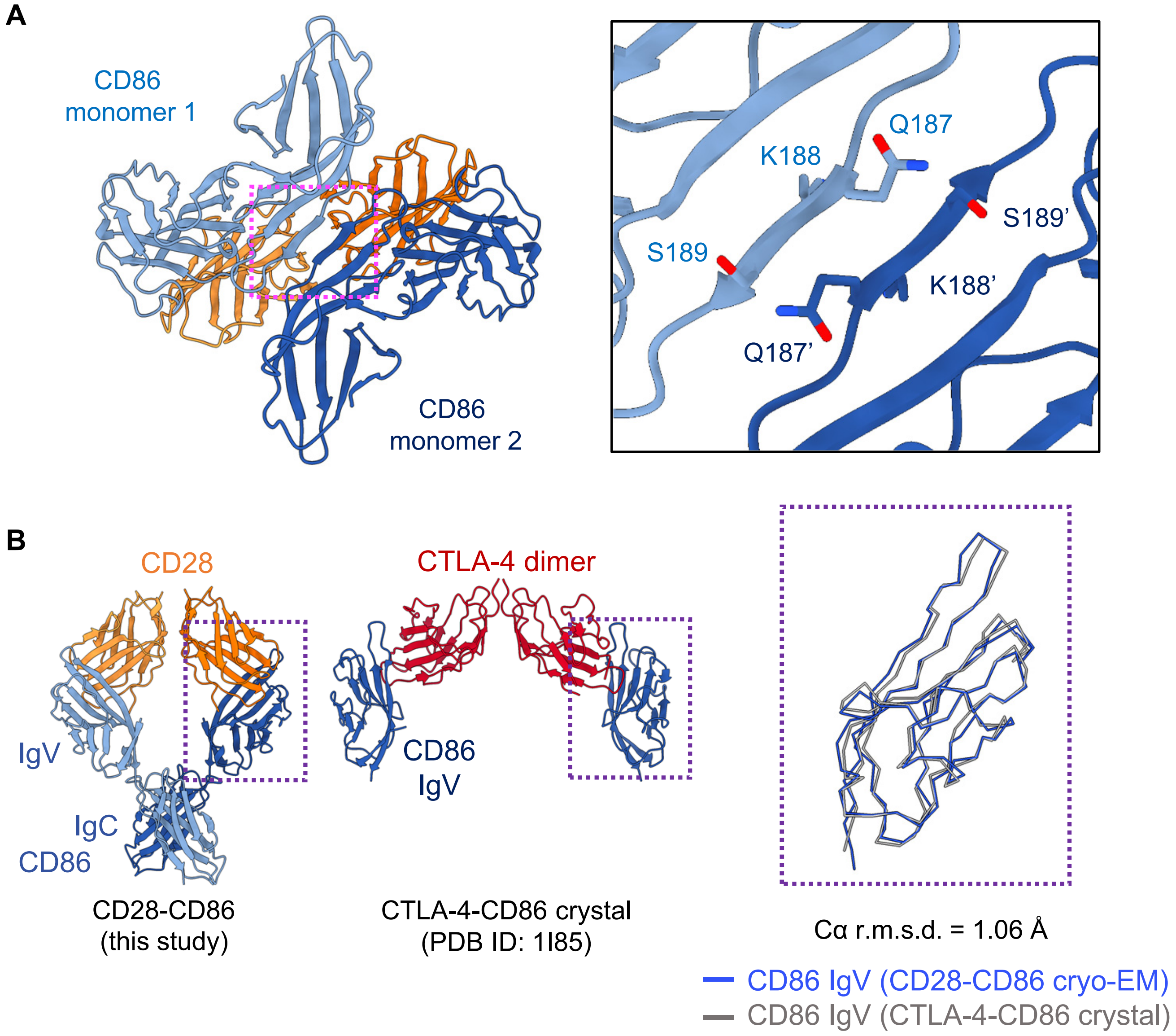

**Figure S3. Structure of the homodimerization interface of CD86.**
(A) Structure of the CD86 homodimeric interface. CD28 dimers are colored dark and light blue, and CD86 homodimers are colored dark and light orange. The CD86 homodimeric interface, highlighted by a dashed magenta box, is shown at higher magnification in the right panel. Amino acid residues from the second CD86 subunit are indicated by prime symbols.
(B) Structural comparison of the CD86 IgV domain. The CD86 IgV domains from the cryo-EM structure of the CD28-CD86 complex (left, Figure 1D) and the previously reported crystal structure of the CTLA-4-CD86 complex (middle) are superimposed in the right panel. The structures of the CD86 IgV domains used for the comparison in the left and middle panels are indicated by dashed purple boxes.

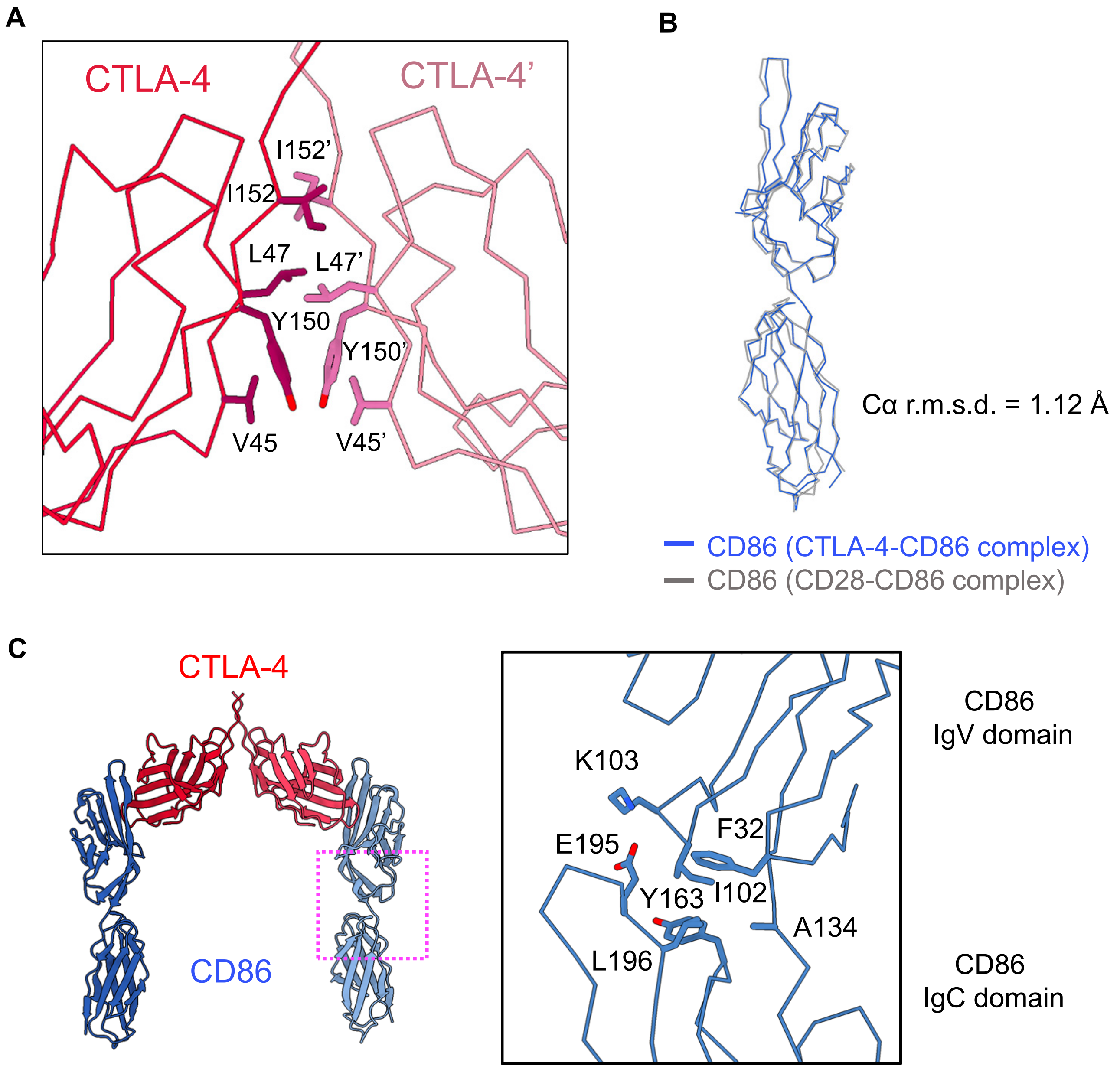

**Figure S4. Structure of the homodimerization interface of CTLA-4.**
(A) Structure of the CTLA-4 homodimerization interface in the CTLA-4-CD86 complex. The two CTLA-4 subunits are colored dark and light red, respectively. Amino acid residues of the second CTLA-4 subunit are indicated by prime symbols. The view is the same as in the lower left panel of Figure 2D.
(B) Structural comparison of CD86 in the CTLA-4-CD86 complex. The CD86 molecules from the cryo-EM structures of the CD28-CD86 and CTLA-4-CD86 complexes are colored grey and blue, respectively, and superimposed. The view is the same as in the lower left panel of Figure 2D.
(C) Structure of the IgV-IgC domain interface of CD86. The region highlighted by the magenta box (left) is shown enlarged in the right panel.

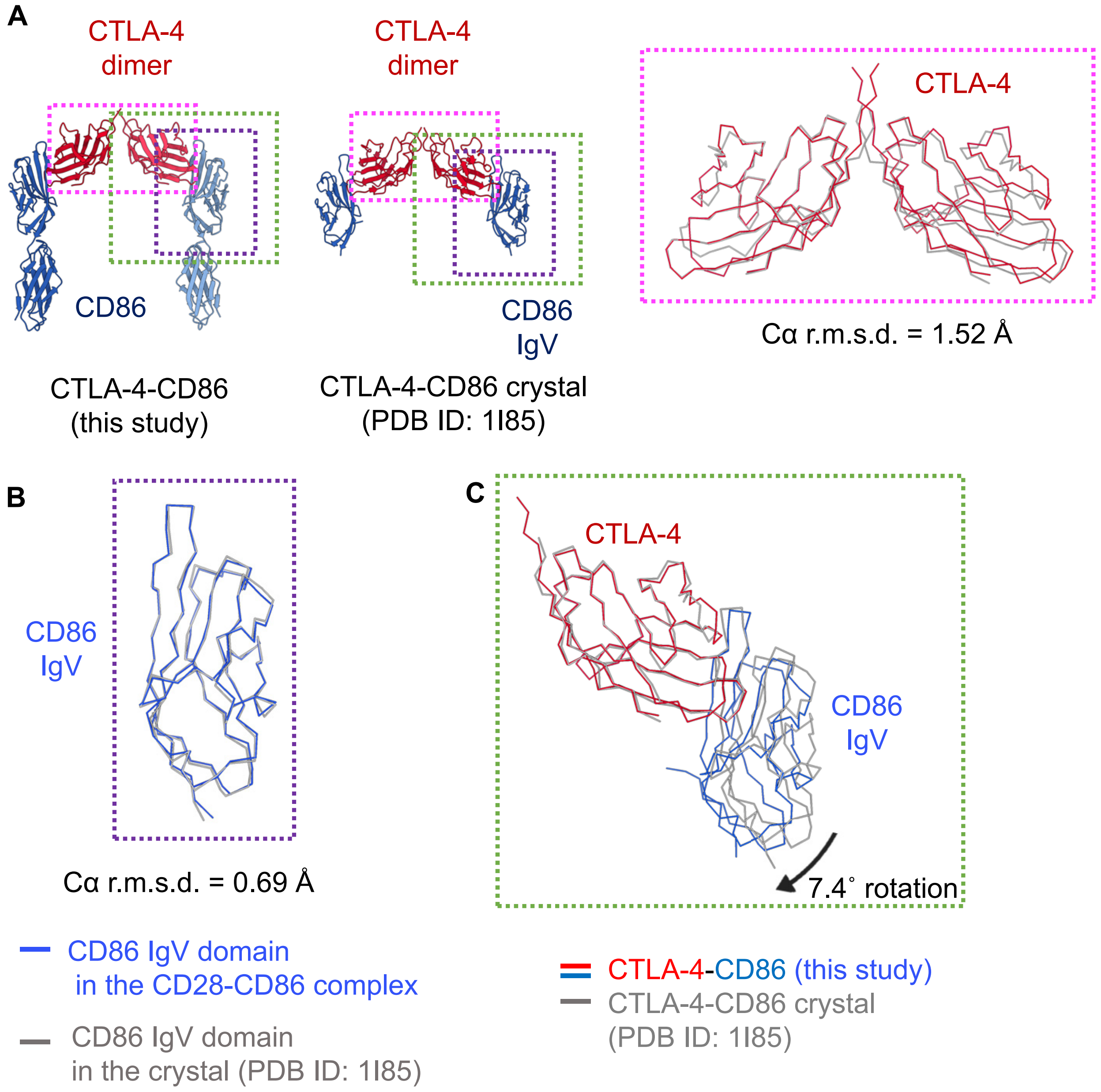

**Figure S5. Structural comparison of CTLA-4-CD86 complexes.**
(A) Structural comparison of CTLA-4 homodimers in the CTLA-4-CD86 complex determined by cryo-EM (left, Figure 2D) and X-ray crystallography (middle). The CTLA-4 dimers from our cryo-EM structure (left) and the previously reported crystal structure (middle) are superimposed in the right panel. The CTLA-4 structures used for the comparison in the left and middle panels are indicated by dashed magenta boxes.
(B) Structural comparison of CD86 in the CTLA-4-CD86 complexes determined by cryo-EM (blue) and X-ray crystallography (grey). The CD86 structures used for the comparison correspond to those indicated by dashed purple boxes in the left and middle panels of (A).
(C) Structural differences at the CTLA-4-CD86 interaction interface between the cryo-EM and X-ray crystal structures. When the CTLA-4 monomers are structurally aligned, the CD86 domain is rotated by ~7.4°. The CTLA-4-CD86 complexes used for the comparison are indicated by dashed green boxes in the left and middle panels of (A).

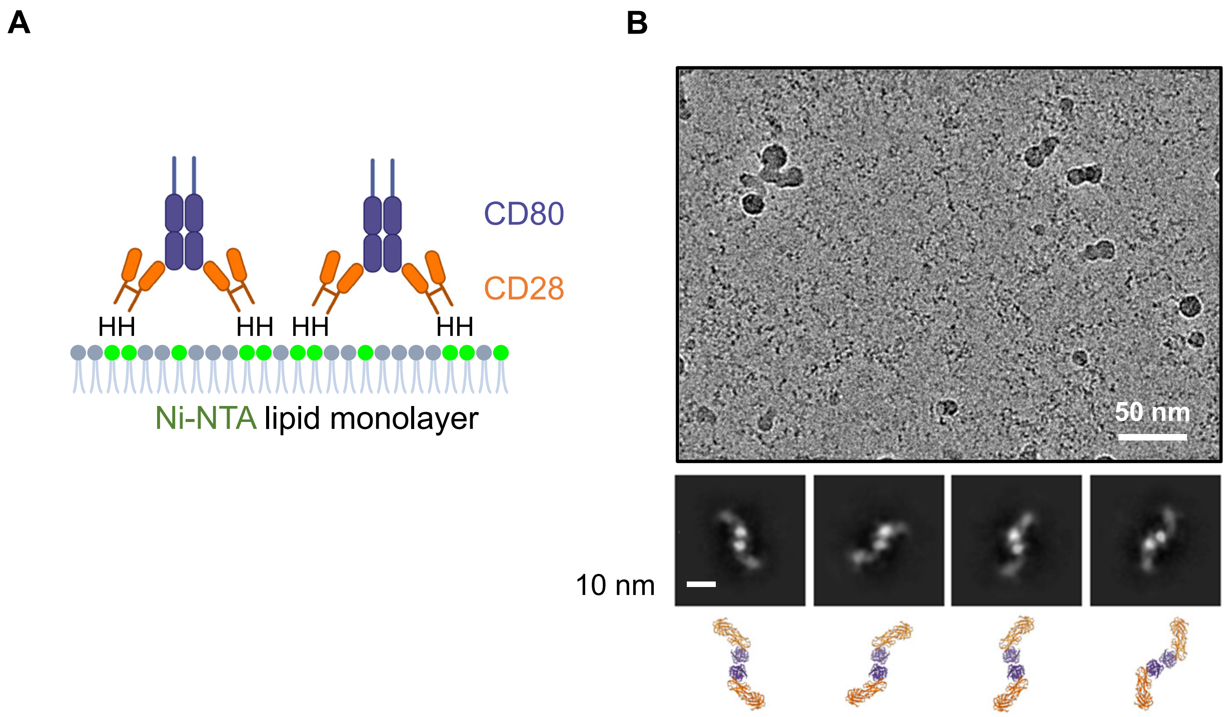

**Figure S6. Inverse attachment of the CD28-CD80 complex to the lipid layer.**
(A) Schematic of the lipid monolayer experiment. An octa-histidine tag was fused to the C-terminus of the CD28 ectodomain, enabling tethering of the CD28-CD80 complex to the $Ni^{2+}$-NTA lipid monolayer. "H" denotes the octa-histidine tag, and Ni2+-NTA lipids are shown in green.
(B) Representative cryo-EM micrograph, with the corresponding 2D class averages shown below. The approximate orientations of the CD28-CD80 complexes are schematically indicated beneath the 2D class averages.

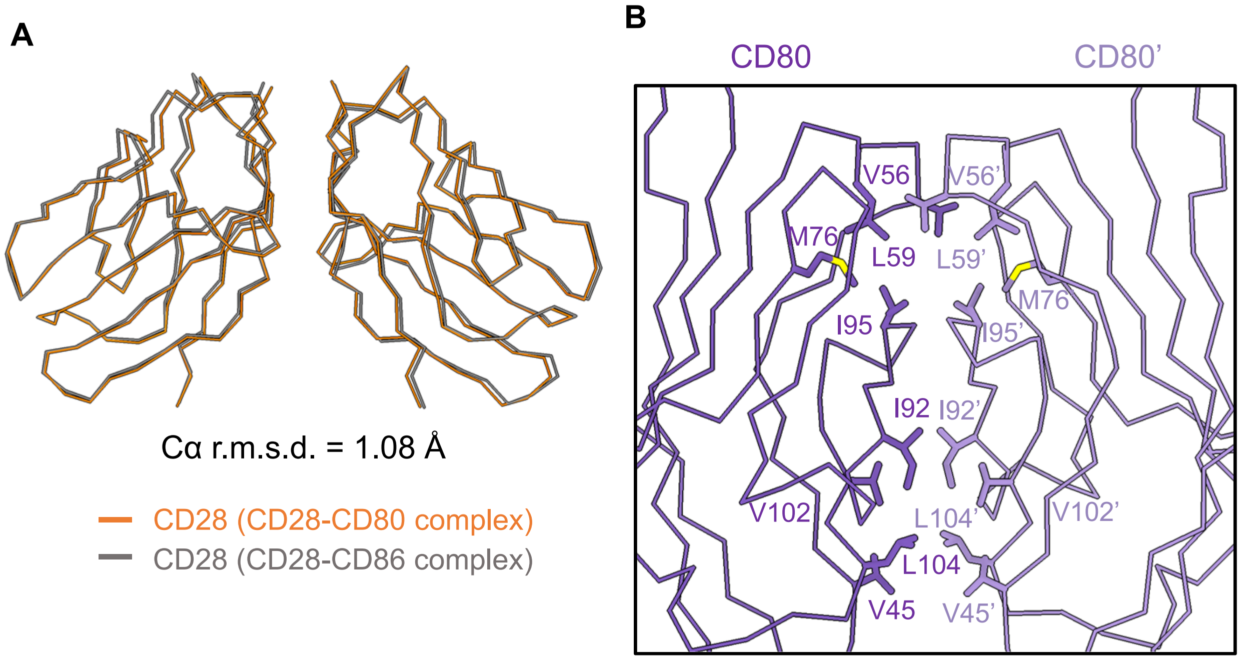

**Figure S7. Structure of the homodimerization interfaces of CD28 and CD80 in the CD28-CD80 complex.**

(A) Structural comparison of CD28 homodimers in the CD28-CD80 (Figure 3D) and CD28-CD86 (Figure 1D) complexes. The view is the same as in the lower panel of Figure 1D.

(B) Close-up view of the CD80 homodimerization interface in the CD28-CD80 complex. The view is the same as in the lower panel of Figure 3D. Residue numbers for the second monomer in the CD80 dimer are indicated by prime symbols.

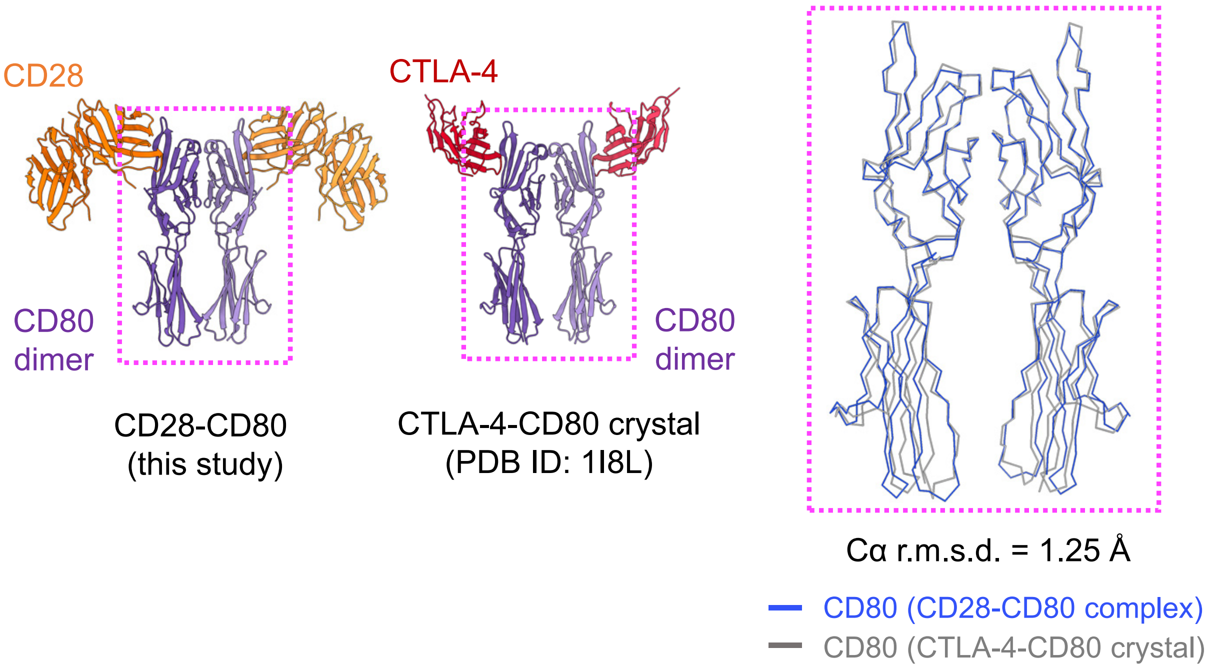

**Figure S8. Structural comparison of CD80 homodimers.**
The structures of the CD80 homodimers in the CD28-CD80 complex (left, Figure 2D) and in the CTLA-4-CD80 complex previously determined by X-ray crystallography (middle) are superimposed in the right panel. The structures of the CD80 homodimers used for the structural comparison are marked with dashed magenta boxes in the left and middle panels.

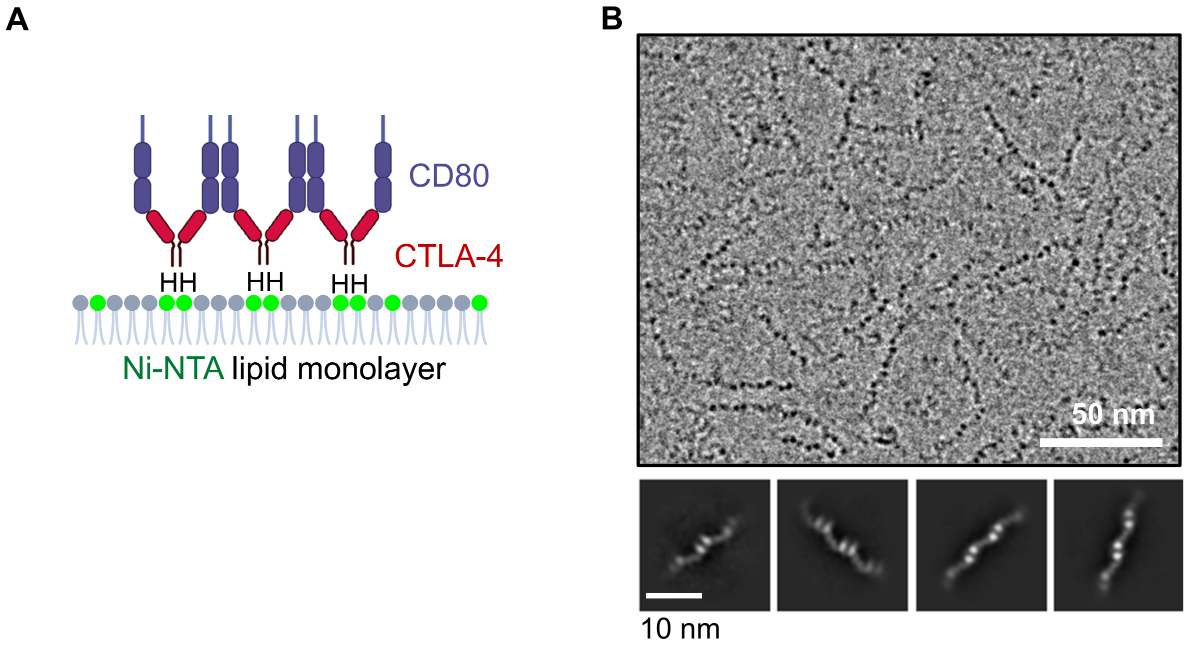

**Figure S9. Inverse attachment of the CTLA-4-CD80 complex to the lipid monolayer**
(A) Schematic of the lipid monolayer experiment. An octa-histidine tag was fused to the C terminus of the CTLA-4 ectodomain, enabling tethering of the CTLA-4-CD80 complex to the $Ni^{2+}$-NTA lipid monolayer. “H” denotes the octa-histidine tag, and $Ni^{2+}$-NTA lipids are shown in green.
(B) Representative cryo-EM micrograph, with corresponding 2D class averages shown below.

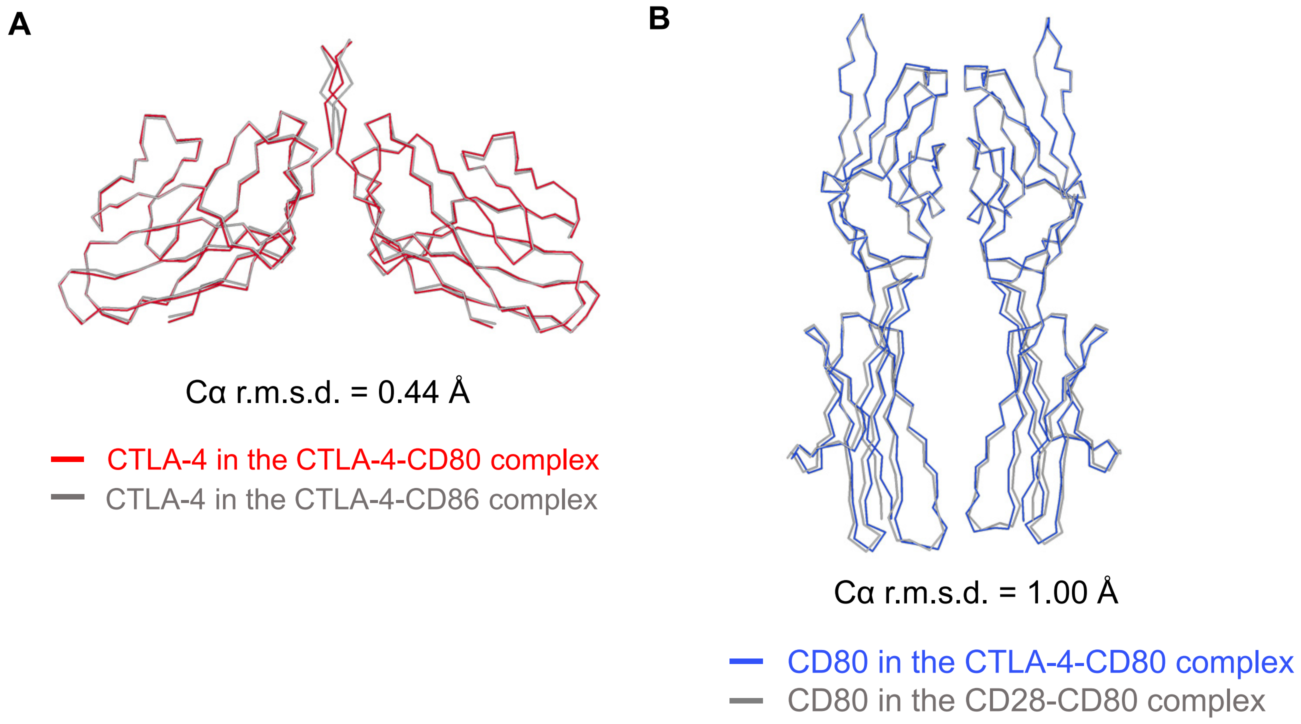

**Figure S10. Structural comparison of CTLA-4 and CD80 homodimers in the CTLA-4-CD80 linear cluster**

(A) Superimposition of CTLA-4 homodimers in the CTLA-4-CD80 and CTLA-4-CD86 complexes.

(B) Superimposition of CD80 homodimers in the CTLA-4-CD80 and CD28-CD80 complexes.

A

B

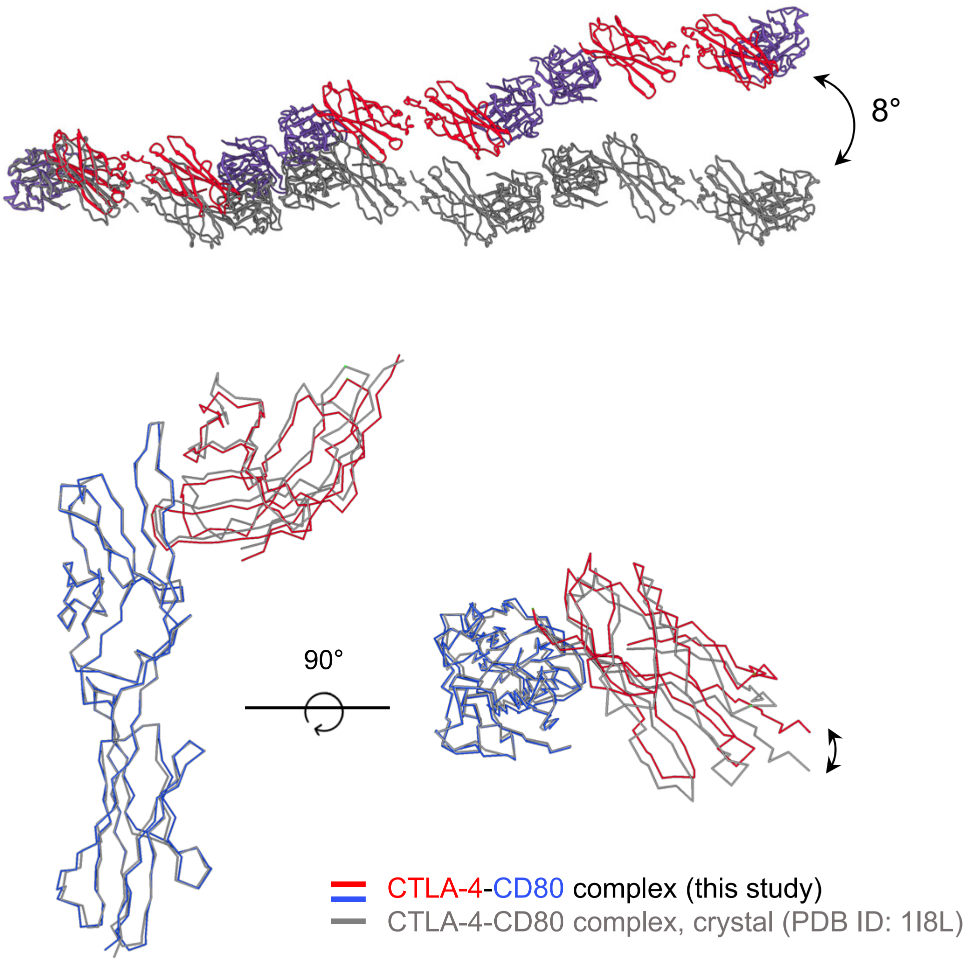

**Figure S11. Comparison of the CTLA-4-CD80 structure in the linear cluster with the crystal structure**

(A) Structural comparison of the CTLA-4-CD80 linear cluster determined in this study with that predicted from the crystal structure. The first CD80 molecules in the two clusters are superimposed to highlight structural differences. The cluster structure derived from the crystal data was generated by assembling the complexes present in the asymmetric unit using crystal symmetries.

(B) Structural differences between the CTLA-4-CD80 binding interfaces in the linear cluster and the crystal structure. The CD80 monomers in the two structures are aligned for comparison.

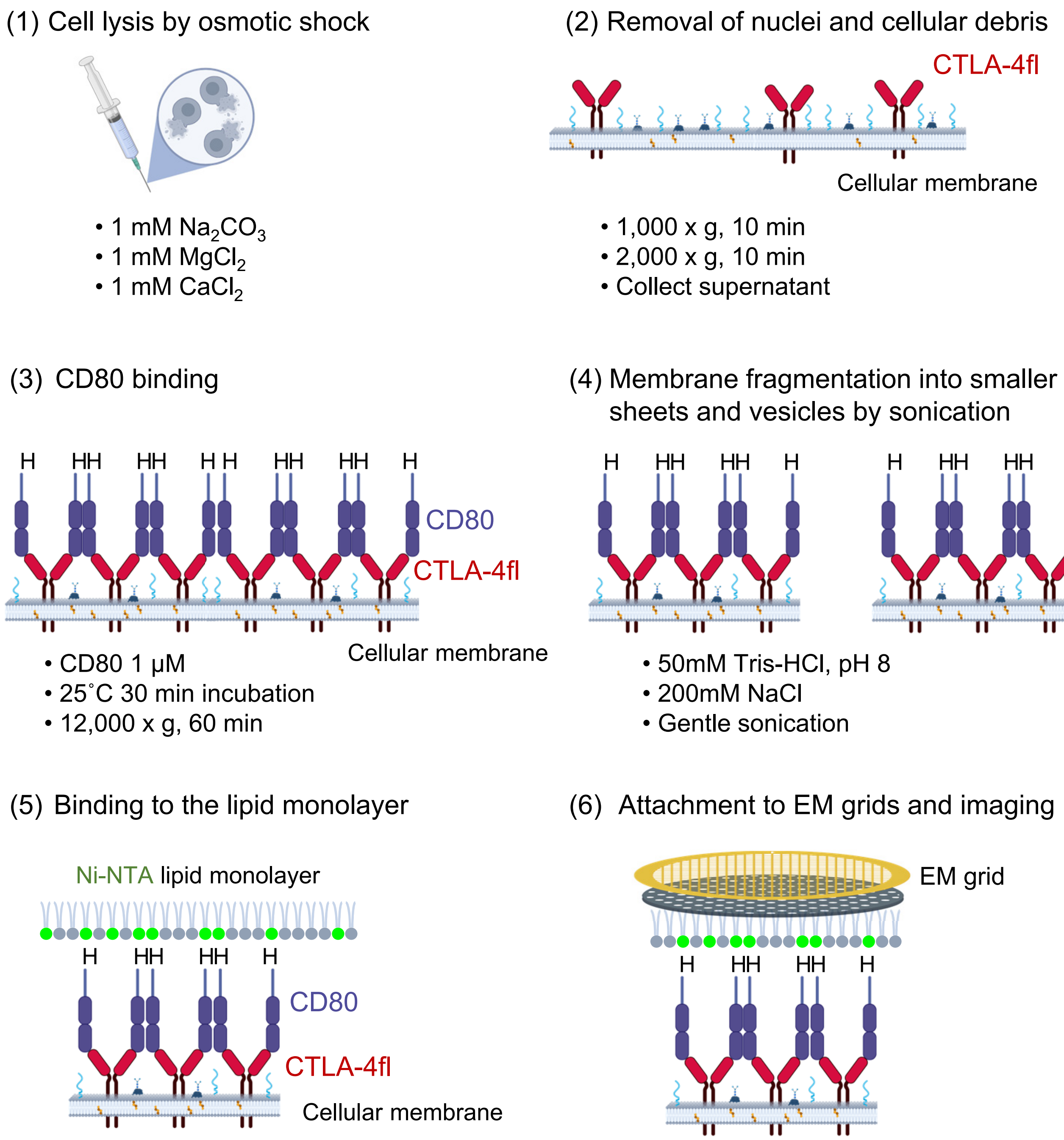

**Figure S12. Preparation of lipid monolayer grids with cellular membrane fragments.** Cells transfected with full-length human CTLA-4 (CTLA-4fl) were lysed, and membrane fragments generated by sonication were tethered to a lipid monolayer. CTLA-4fl and CD80 are schematically depicted in red and blue, respectively. "H" denotes the octa-histidine tag, and $Ni^{2+}$–NTA lipids are shown in green. Other proteins and carbohydrates present on the cellular membrane are illustrated as short curved lines in cyan and blue, respectively. Cholesterol is represented by an orange line.

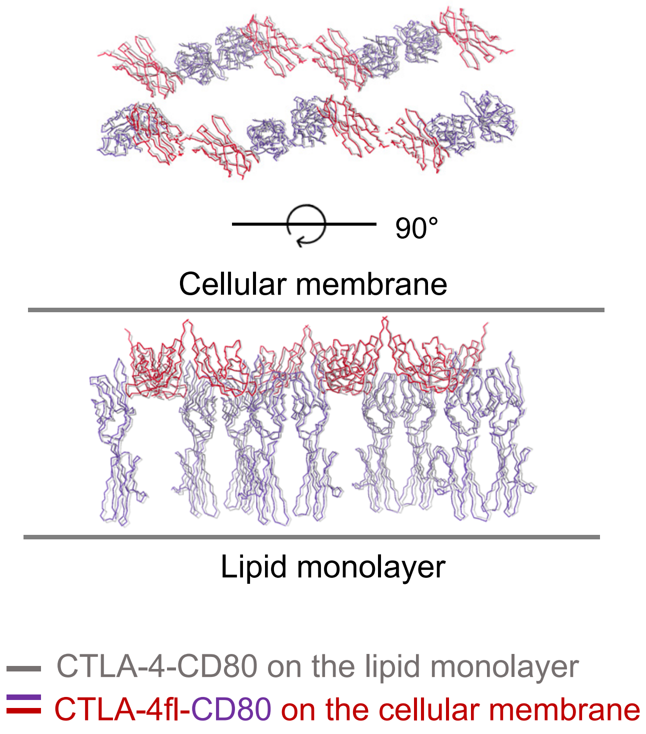

**Figure S13. Structural comparison of the two-dimensional lattices of the CTLA-4-CD80 complexes.** Alignment of the CTLA-4-CD80 complex formed on the $Ni^{2+}$-NTA lipid monolayer (Figure 5C) with the CTLA-4fl-CD80 complex formed on the cellular membrane (Figure 6D).

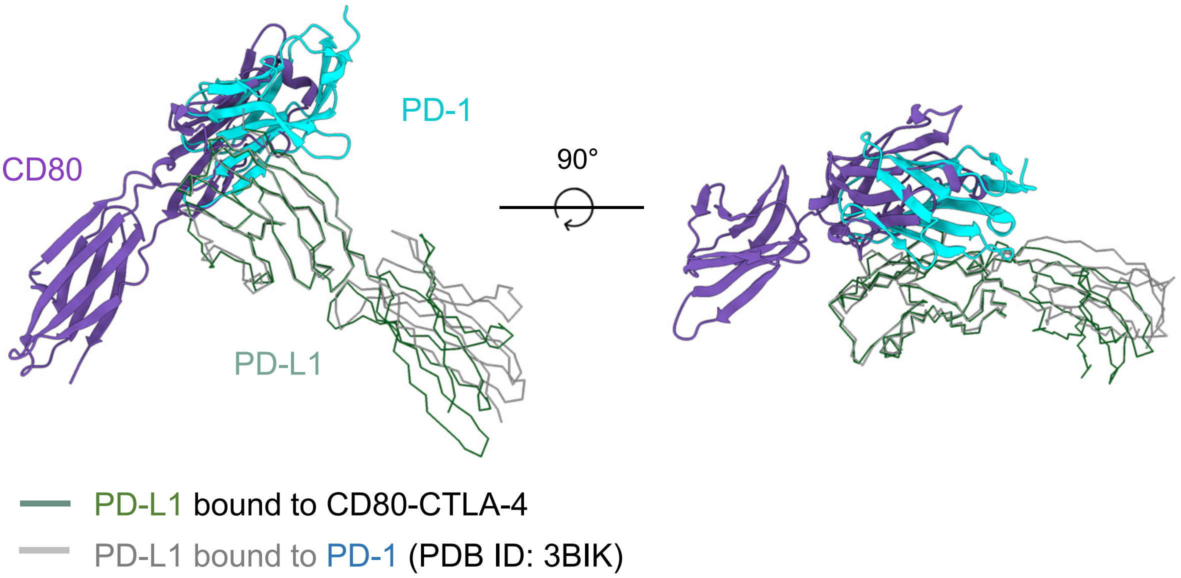

**Figure S14. Comparison of the PD-1 and CD80 binding sites on PD-L1**
Superimposition of the CD80-PD-L1 and PD-1-PD-L1 structures, with the IgV domains of PD-L1 aligned.

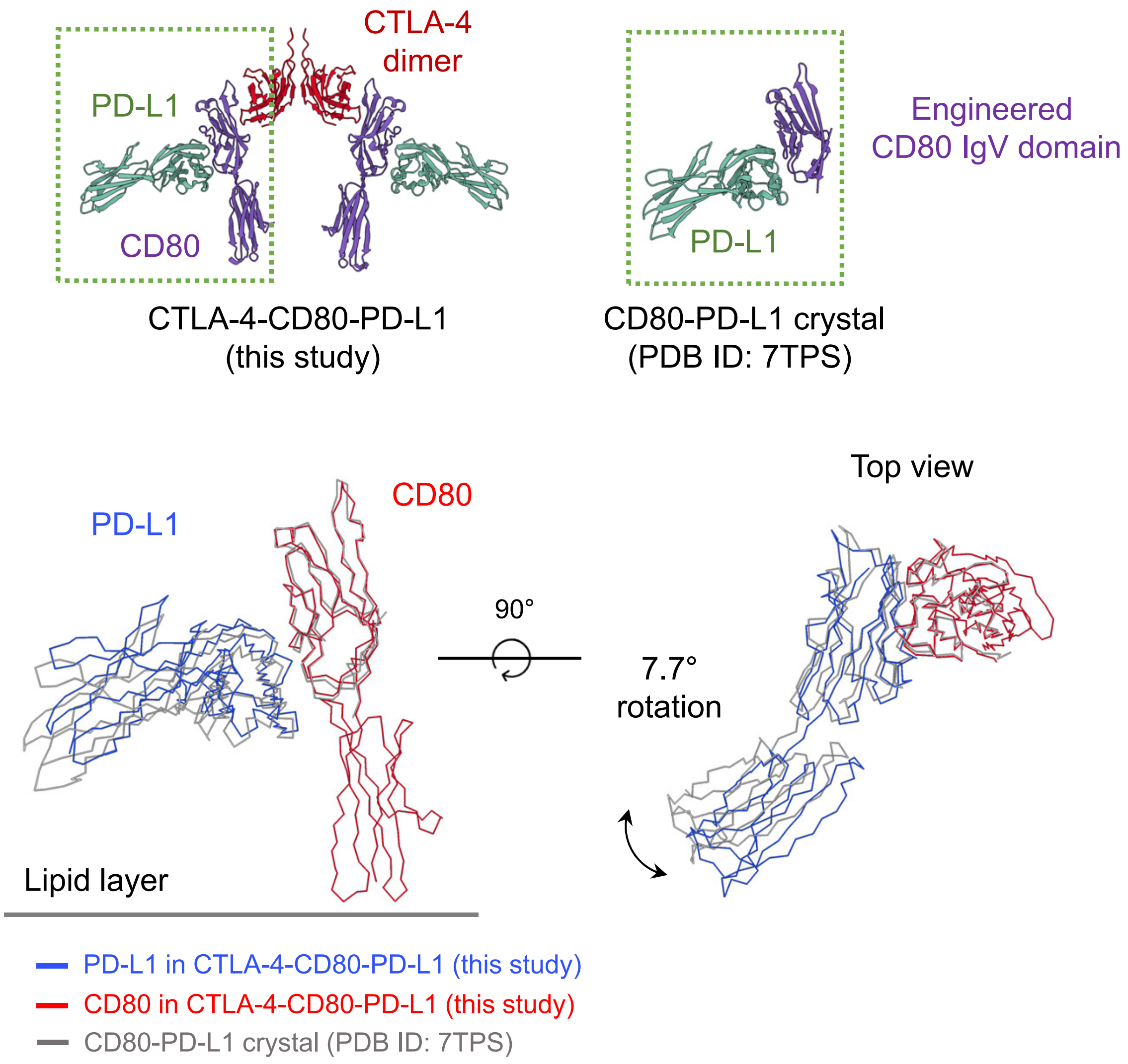

**Figure S15. Structural comparison of the PD-L1-CD80 complexes**

The IgV domain structures of CD80 from the CTLA-4-CD80-PD-L1 complex (upper left; Figure 6E) and the previously reported PD-L1-CD80 crystal structure (upper right; PDB ID: 7TPS) are superimposed in the lower panels. The PD-L1-CD80 structures used for comparison in the upper left and right panels are indicated by dashed green boxes. Note that CD80 in the crystal structure includes only the IgV domain, which features seven mutations to enhance binding affinity.

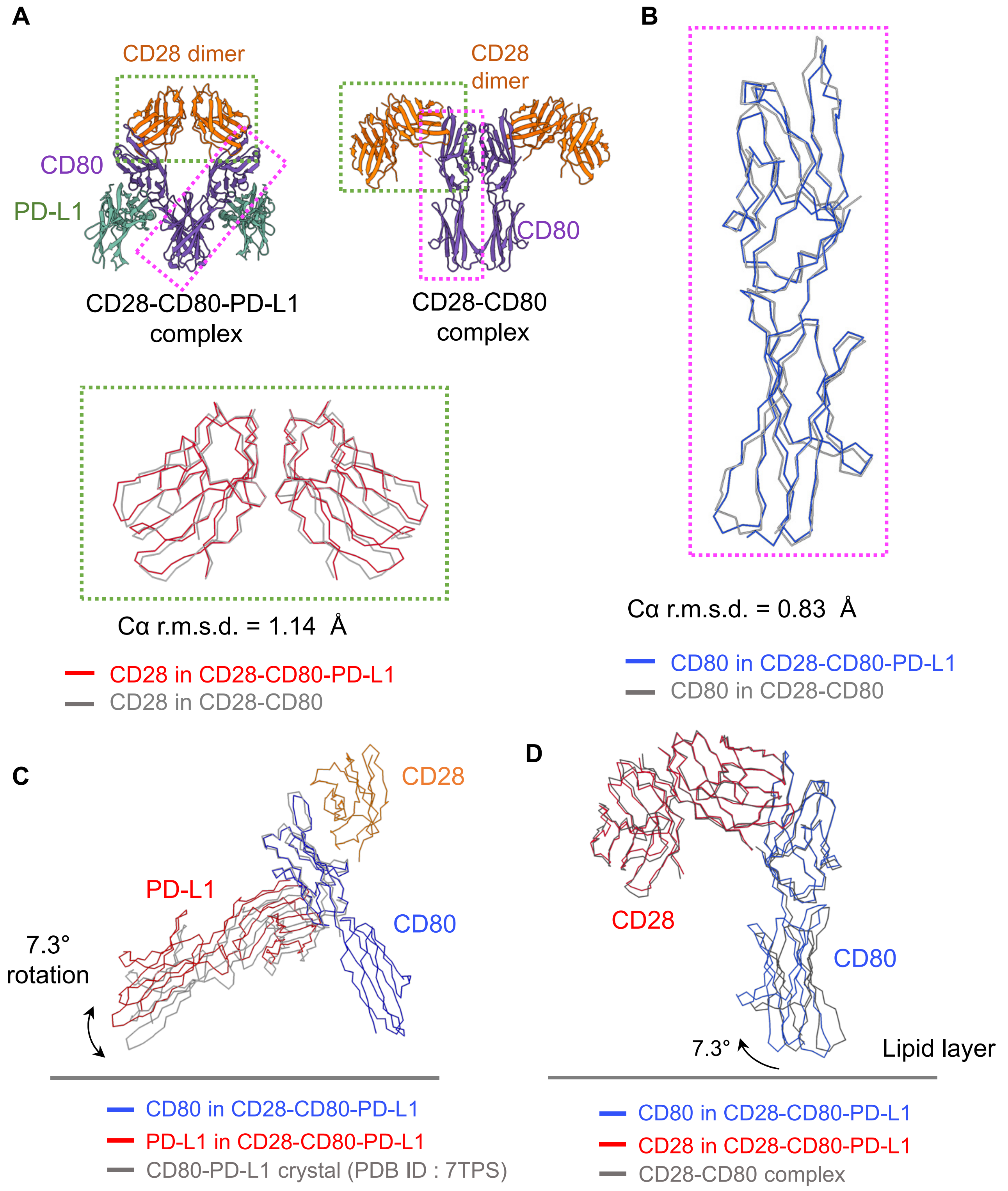

**Figure S16. Structural comparison of CD28 and CD80 in the CD28-CD80-PD-L1 and CD28-CD80 complexes**

(A) Structural comparison of CD28 homodimers. CD28 dimers from the CD28-CD80-PD-L1 complex (upper left; Figure 7D) and the CD28-CD80 complex (upper right; Figure 3D) are superimposed in the lower panel. The CD28 structures used for comparison in the upper left and right panels are indicated by dashed green boxes.

(B) Structural comparison of CD80. CD80 structures from the CD28-CD80-PD-L1 and CD28-CD80 complexes are superimposed. The CD80 structures used for the comparison in (A) are indicated by dashed magenta boxes.

(C) Structural comparison of the CD80-PD-L1 interface. PD-L1 structures are compared after superposition of the CD80 IgV domains from the CD28-CD80-PD-L1 complex and the CD80-PD-L1 crystal structure (PDB ID: 7TPS). Note that CD80 in the crystal structure contains seven mutations.

(D) Structural comparison of the CD28-CD80 interface. CD80 structures are compared after superposition of the right monomers of the CD28 dimers from the CD28-CD80-PD-L1 and CD28-CD80 complexes.

**Methods S1: Cryo-EM processing schematics, related to Figures 1-8**

Structural basis of CD28 and CTLA-4 interactions with CD80, CD86, and the CD80-PD-L1 heterodimer on artificial and cellular membranes

Gu Min Han, Chan Seok Lim and Jie-Oh Lee

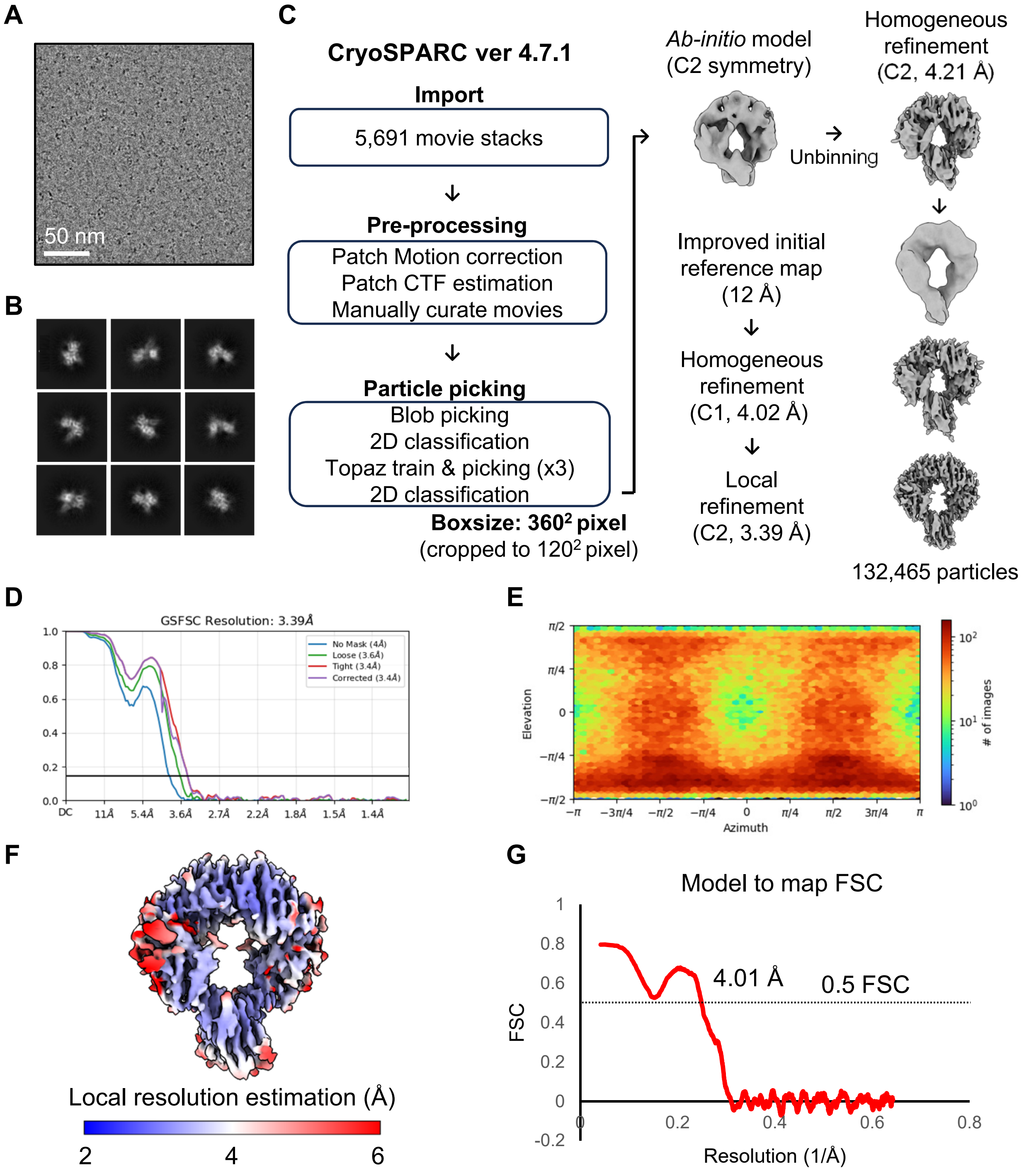

**Processing Schematic 1. Cryo-EM data processing of the CD28-CD86 complex. Related to Figure 1.**

(A) A representative micrograph image. (B) Selected 2D class averages. (C) Summary of cryo-EM data processing. (D) Map Fourier Shell Correlation (FSC) curve. (E) Orientation distribution of the particles used for 3D electron density reconstruction. (F) Electron density map colored according to the local resolution estimation. (G) Model to map FSC curve.

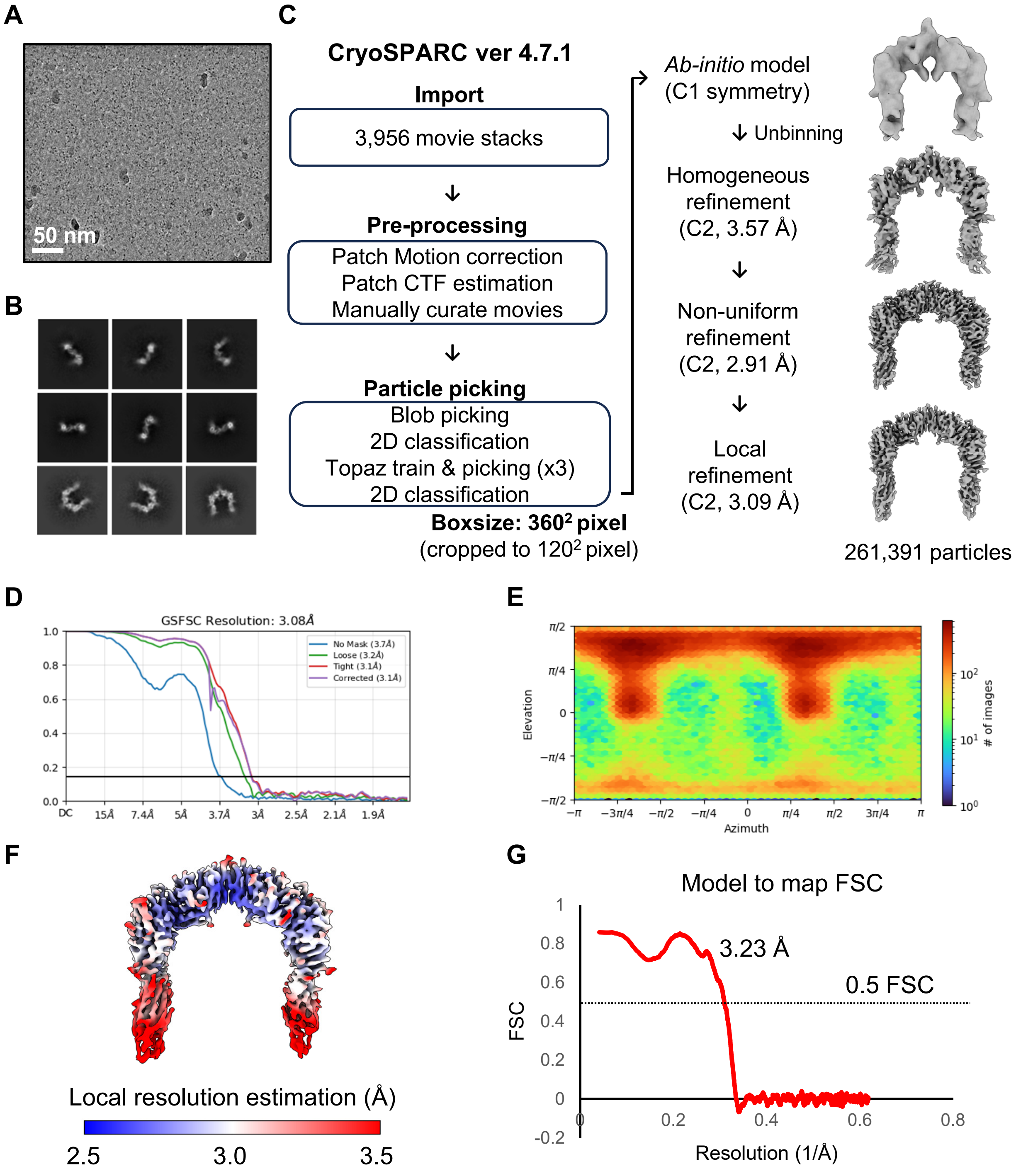

**Processing Schematic 2. Cryo-EM data processing of the CTLA-4-CD86 complex. Related to Figure 2.**

(A) A representative micrograph image. (B) Selected 2D class averages. (C) Summary of cryo-EM data processing. (D) Map Fourier Shell Correlation (FSC) curve. (E) Orientation distribution of the particles used for 3D electron density reconstruction. (F) Electron density map colored according to the local resolution estimation. (G) Model to map FSC curve.

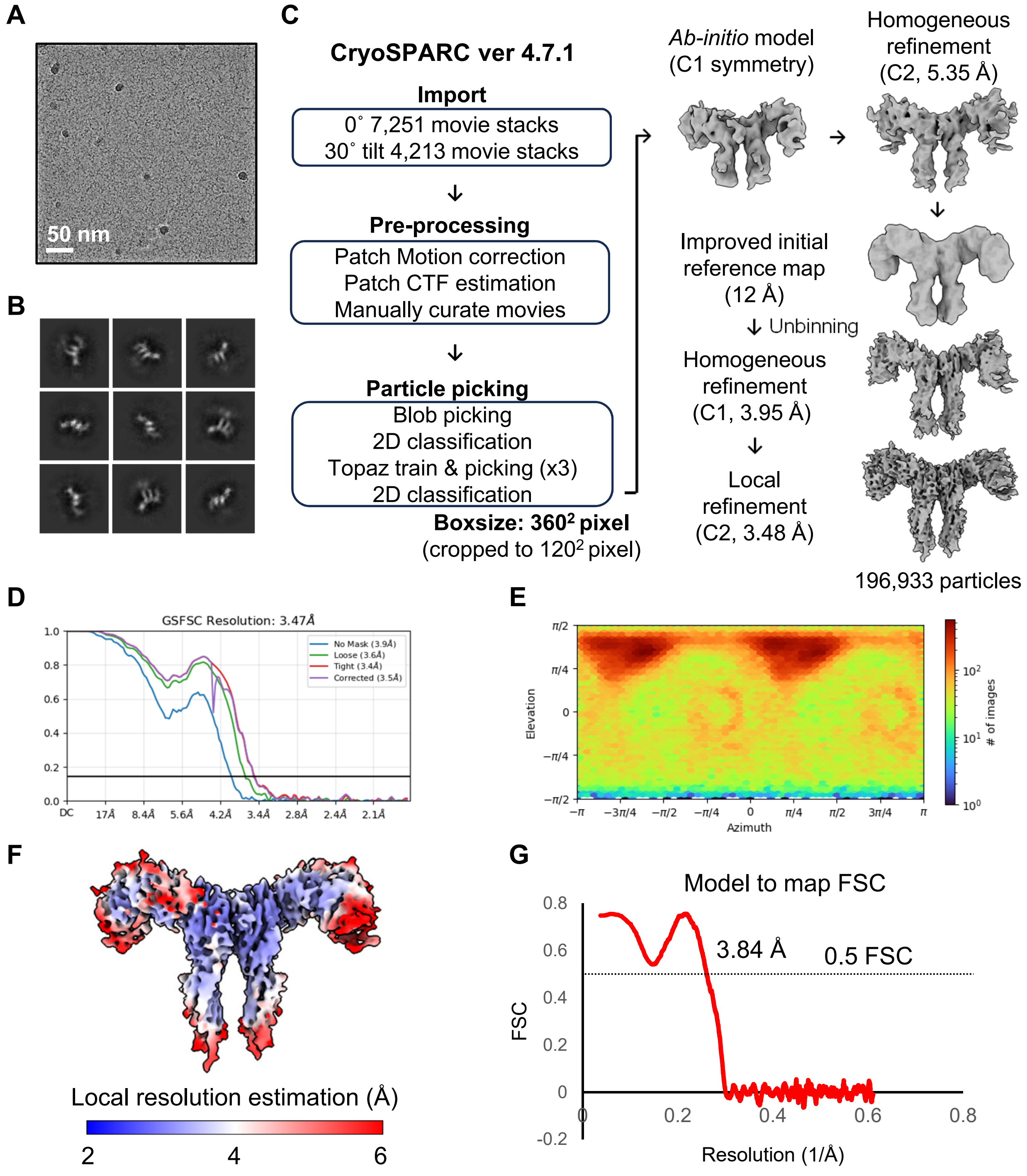

**Processing Schematic 3. Cryo-EM data processing of the CD28-CD80 complex. Related to Figure 3.**
(A) A representative micrograph image. (B) Selected 2D class averages. (C) Summary of cryo-EM data processing. (D) Map Fourier Shell Correlation (FSC) curve. (E) Orientation distribution of the particles used for 3D electron density reconstruction. (F) Electron density map colored according to the local resolution estimation. (G) Model to map FSC curve.

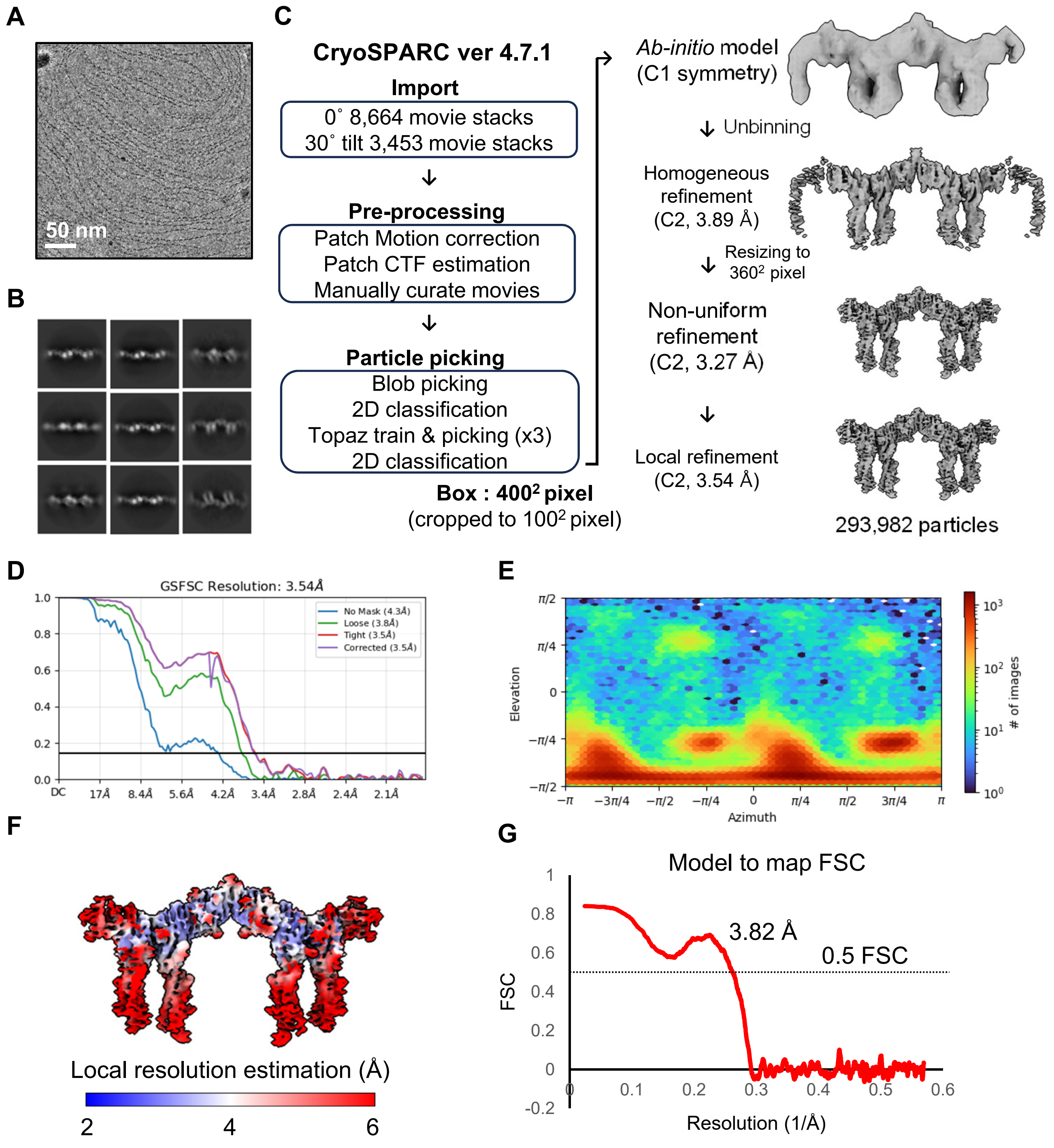

**Processing Schematic 4. Cryo-EM data processing of the CTLA-4-CD80 complex (CTLA-4 focused). Related to Figure 4.**

(A) A representative micrograph image. (B) Selected 2D class averages. (C) Summary of cryo-EM data processing. (D) Map Fourier Shell Correlation (FSC) curve. (E) Orientation distribution of the particles used for 3D electron density reconstruction. (F) Electron density map colored according to the local resolution estimation. (G) Model to map FSC curve.

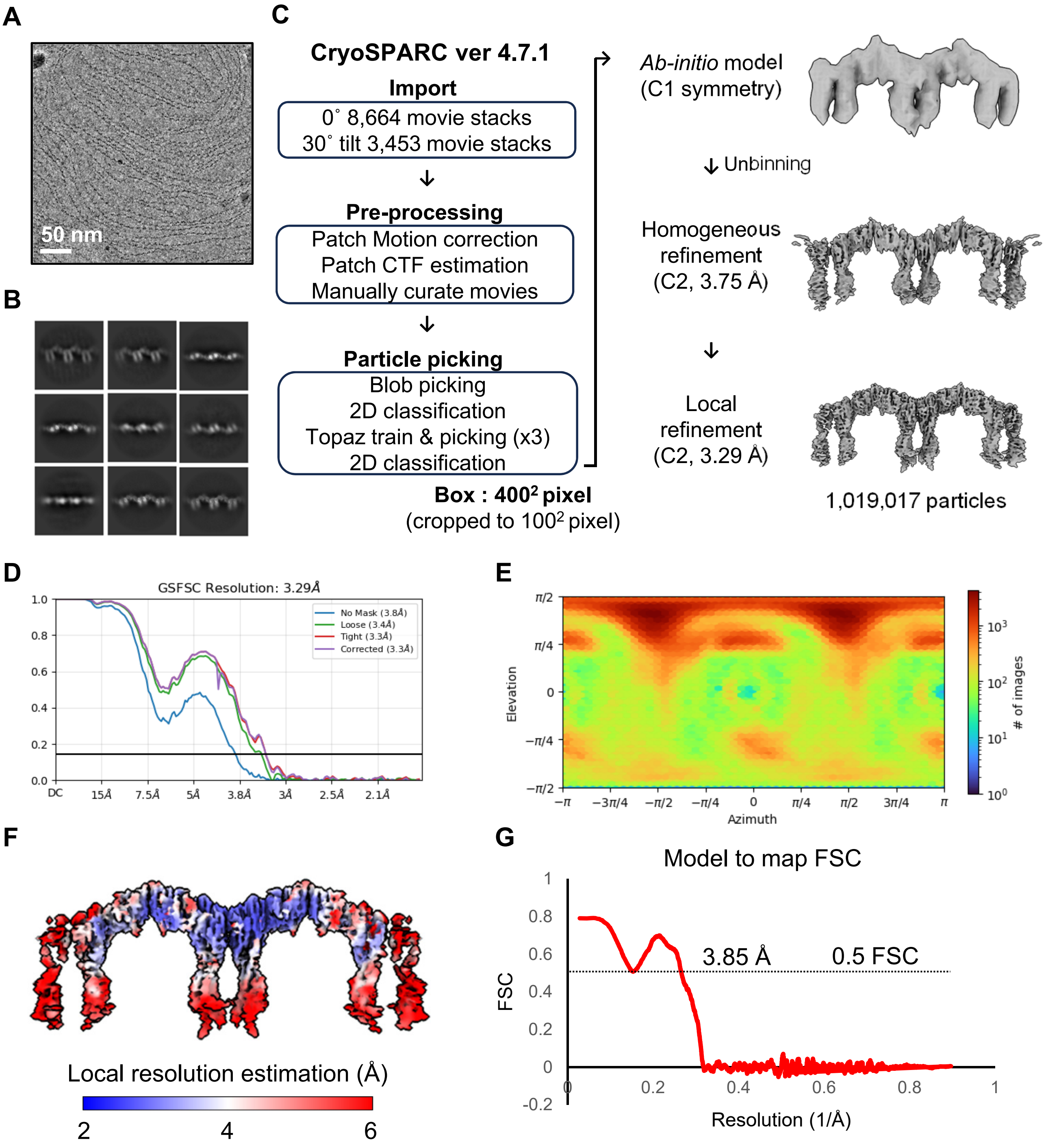

**Processing Schematic 5. Cryo-EM data processing of the CTLA-4-CD80 complex (CD80 focused). Related to Figure 4.**

(A) A representative micrograph image. (B) Selected 2D class averages. (C) Summary of cryo-EM data processing. (D) Map Fourier Shell Correlation (FSC) curve. (E) Orientation distribution of the particles used for 3D electron density reconstruction. (F) Electron density map colored according to the local resolution estimation. (G) Model to map FSC curve.

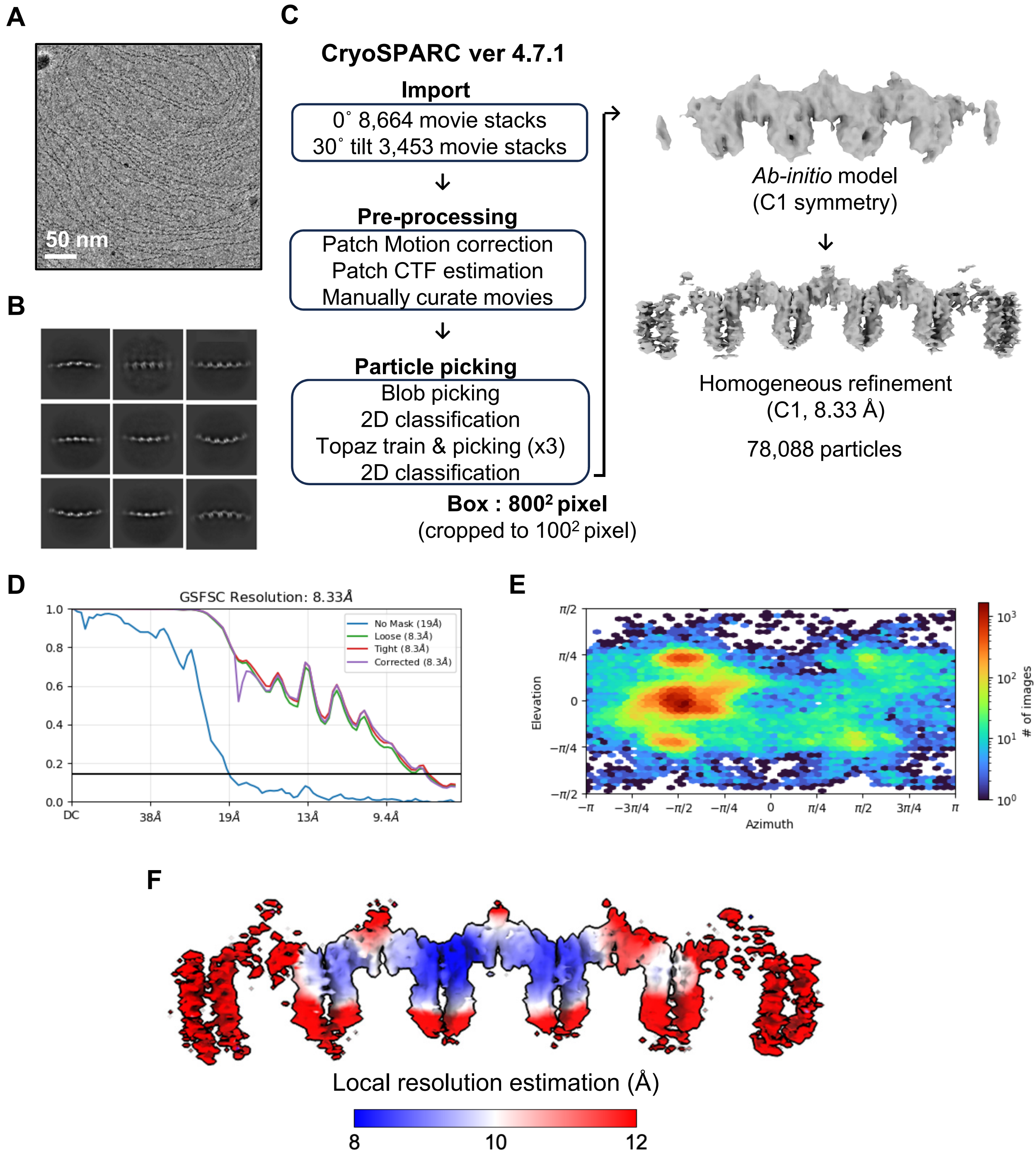

**Processing Schematic 6. Cryo-EM data processing of the CTLA-4-CD80 complex (linear cluster). Related to Figure 4.**

(A) A representative micrograph image. (B) Selected 2D class averages. (C) Summary of cryo-EM data processing. (D) Map Fourier Shell Correlation (FSC) curve. (E) Orientation distribution of the particles used for 3D electron density reconstruction. (F) Electron density map colored according to the local resolution estimation.

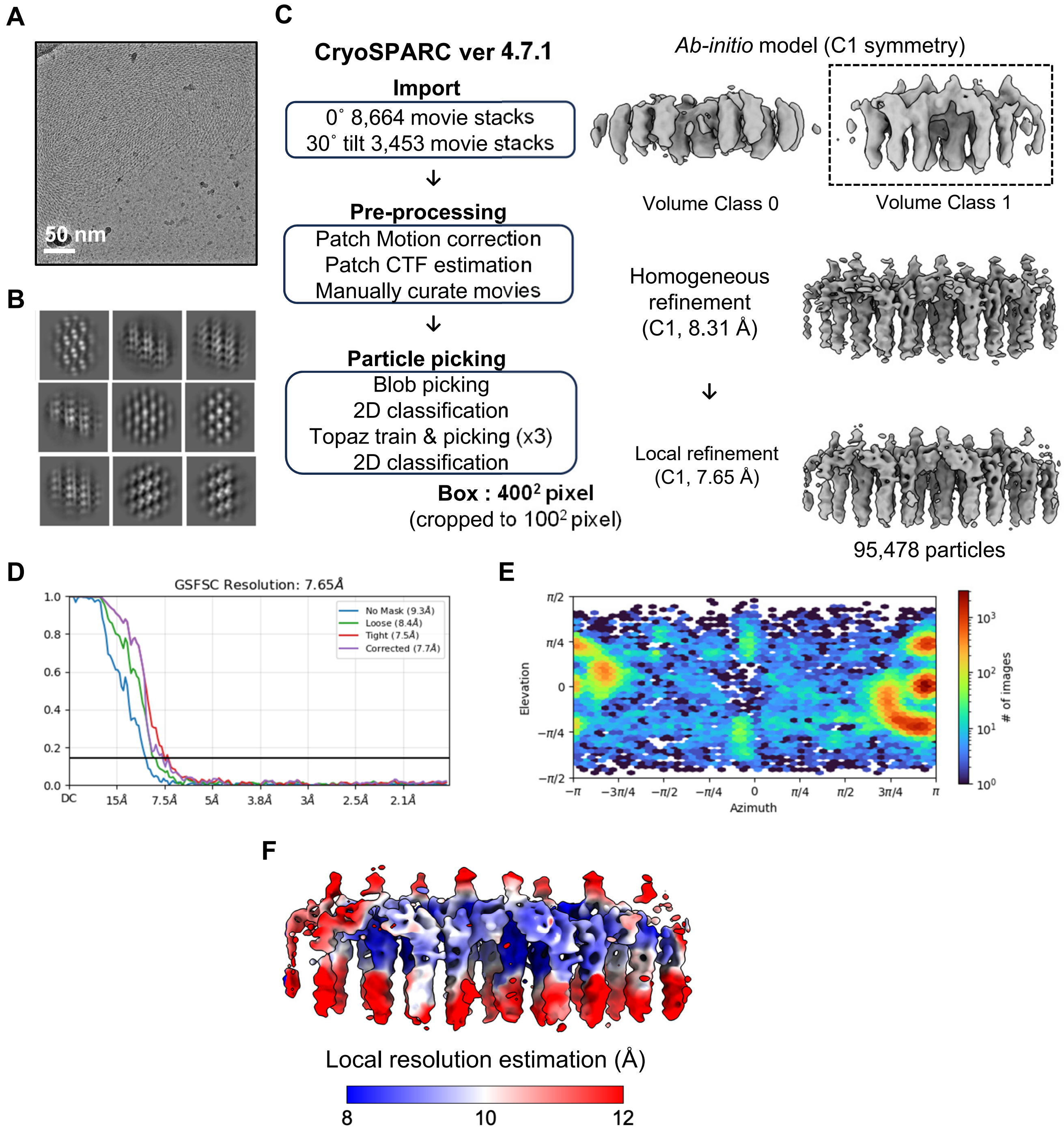

**Processing Schematic 7. Cryo-EM data processing of the CTLA-4-CD80 complex (2D cluster). Related to Figure 5.**

(A) A representative micrograph image. (B) Selected 2D class averages. (C) Summary of cryo-EM data processing. (D) Map Fourier Shell Correlation (FSC) curve. (E) Orientation distribution of the particles used for 3D electron density reconstruction. (F) Electron density map colored according to the local resolution estimation.

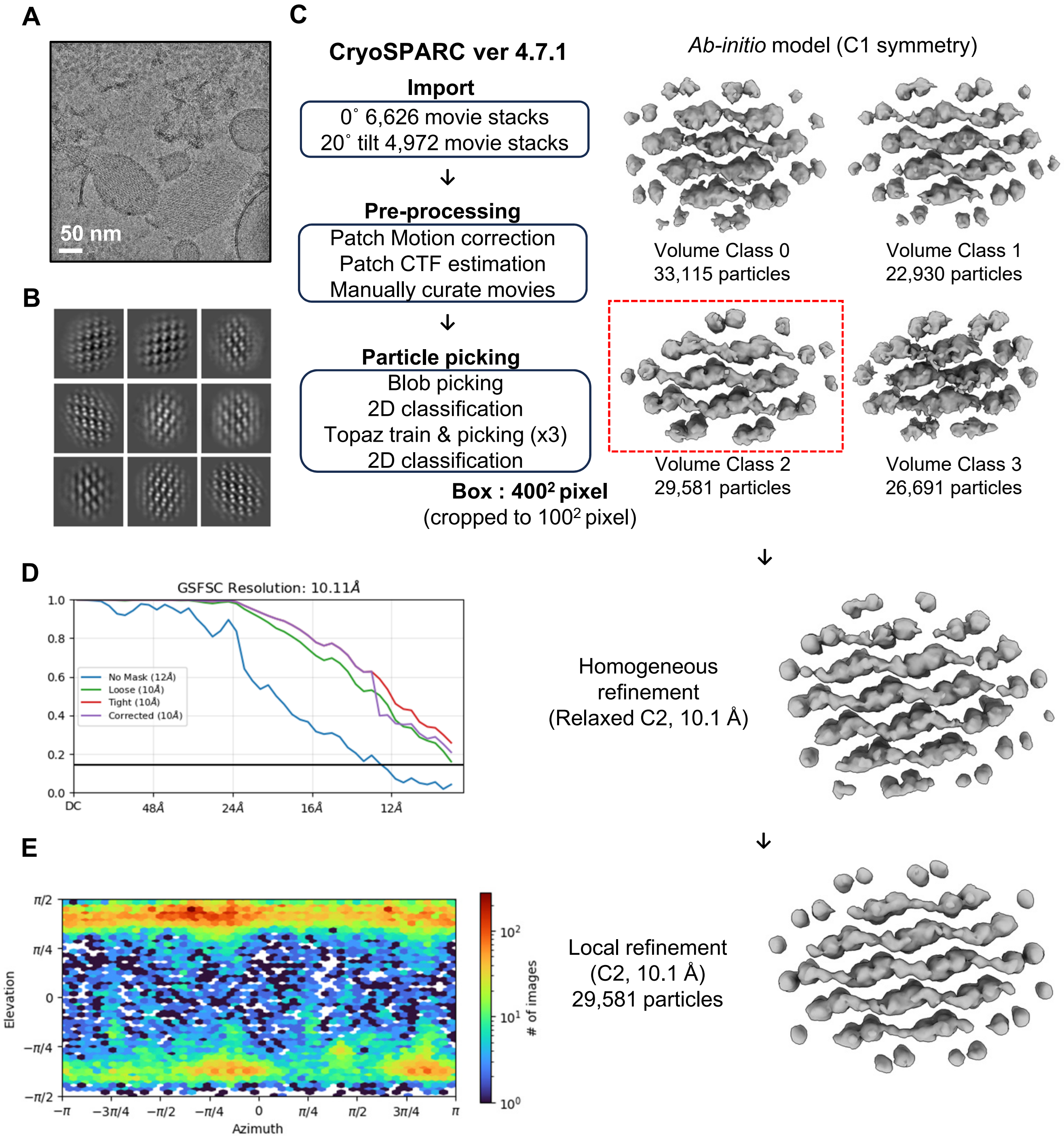

**Processing Schematic 8. Cryo-EM data processing of the CTLA-4-CD80 complex (2D cluster) on the cellular membrane fragment. Related to Figure 6.**

(A) A representative micrograph image. (B) Selected 2D class averages. (C) Summary of cryo-EM data processing. (D) Map Fourier Shell Correlation (FSC) curve. (E) Orientation distribution of the particles used for 3D electron density reconstruction.

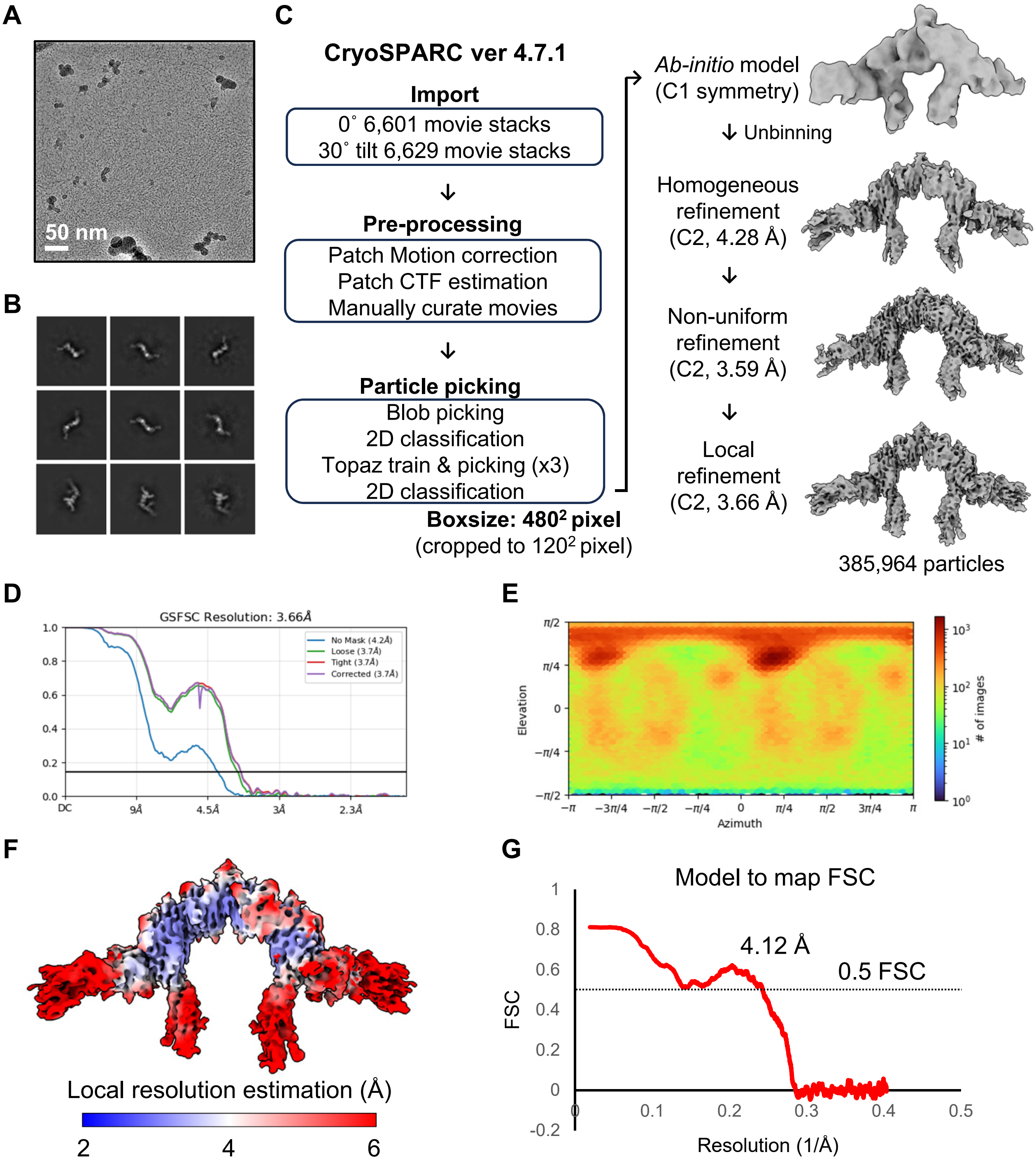

**Processing Schematic 9. Cryo-EM data processing of the CTLA-4-CD80-PD-L1 complex. Related to Figure 7.**

(A) A representative micrograph image. (B) Selected 2D class averages. (C) Summary of cryo-EM data processing. (D) Map Fourier Shell Correlation (FSC) curve. (E) Orientation distribution of the particles used for 3D electron density reconstruction. (F) Electron density map colored according to the local resolution estimation. (G) Model to map FSC curve.

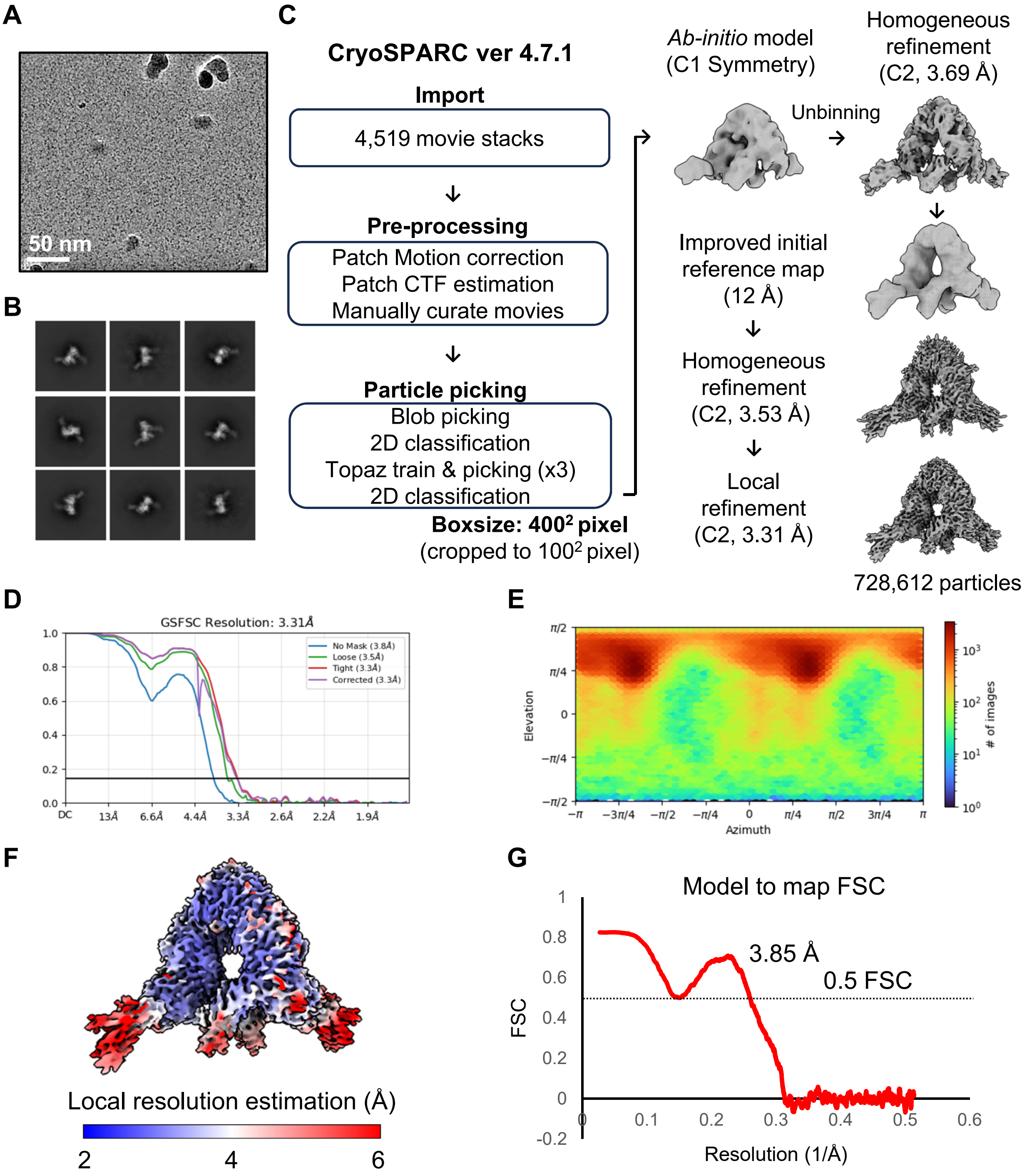

**Processing Schematic 10. Cryo-EM data processing of the CD28-CD80-PD-L1 complex. Related to Figure 8.**

(A) A representative micrograph image. (B) Selected 2D class averages. (C) Summary of cryo-EM data processing. (D) Map Fourier Shell Correlation (FSC) curve. (E) Orientation distribution of the particles used for 3D electron density reconstruction. (F) Electron density map colored according to the local resolution estimation. (G) Model to map FSC curve.
